# Bringing TrackMate into the era of machine-learning and deep-learning

**DOI:** 10.1101/2021.09.03.458852

**Authors:** Dmitry Ershov, Minh-Son Phan, Joanna W. Pylvänäinen, Stéphane U. Rigaud, Laure Le Blanc, Arthur Charles-Orszag, James R. W. Conway, Romain F. Laine, Nathan H. Roy, Daria Bonazzi, Guillaume Duménil, Guillaume Jacquemet, Jean-Yves Tinevez

## Abstract

TrackMate is an automated tracking software used to analyze bioimages and distributed as a Fiji plugin. Here we introduce a new version of TrackMate rewritten to improve performance and usability, and integrating several popular machine and deep learning algorithms to improve versatility. We illustrate how these new components can be used to efficiently track objects from brightfield and fluorescence microscopy images across a wide range of bio-imaging experiments.

---

Object tracking is an essential image analysis technique used across biosciences to quantify dynamic processes. In life sciences, tracking is used for instance to track single particles, sub-cellular organelles, bacteria, cells, and whole animals. Due to the diversity of images to analyze, no single software can address every Life-Sciences research tracking challenges. This prompted for flexible and extensible software tracking platforms [1–5] that enable biologists to build automated tracking pipelines tailored for a specific problem.

Most tracking algorithms proceed in two steps. First, a detection algorithm detects or segments individual objects at each time point. Second, a linking algorithm links the detections across time points and builds tracks that follow each object over time. Importantly, accurately detecting or segmenting objects is crucial to yield accurate tracking results [6]. However, the low signal-to-noise ratio (SNR) typical of live-cell fluorescence microscopy often makes it challenging to accurately track all objects. Overlooked objects then result in the linking part generating short tracks that end prematurely, creating multiple short tracks for these objects. Also, objects at high density can be challenging to individualize as they often overlap or come in contact. Most detection algorithms will treat them as a single detection, resulting in breaks in tracks or in single tracks linking groups of objects.

Several linking algorithms can partly rescue these issues, but overall all tracking algorithms tested in [6] displayed a decreasing performance with increasing object density and decreasing SNR. Machine-learning (ML) and deep learning (DL) approaches can address these challenges. Indeed ML and DL approaches have been shown to excel at image segmentation tasks in low SNR and high-density images [7]. In fact, several groups already used DL to perform successfully complex tracking problems [7–11]. However, many of these approaches remain specific to a problem or challenging to employ as they often require dedicated coding skills and specialized hardware.

We previously developed TrackMate [4], a user-friendly Fiji [12] plugin for tracking objects in fluorescence microscopy images. TrackMate offers automated and semi-automated tracking algorithms, together with advanced visualization and analysis tools. TrackMate is interactive and enables users to filter and curate tracking results based on defined parameters, and as such, it can accommodate a wide range of tracking challenges. TrackMate has a strong focus on interoperability and can be scripted (Jython in Fiji, MATLAB via MIJ [13]), exchange data with other tracking software (Icy [14], MTrackJ [15]) or analysis packages (MATLAB, R, Python). TrackMate is also a software platform that can easily be extended by adding new tracking algorithms, analysis, and visualization features. Thanks to these features, TrackMate has built a large user and developer base [16–18].

Until now, TrackMate detectors were solely based on the Laplacian of Gaussian (LoG) filter. The LoG filter is efficient against sub-resolved particles [19] or other blob-like objects but performs poorly for textured objects, for objects with complex shapes, and other imaging modalities than fluorescence. These detectors are also limited to measuring the object’s position and not their shape.

Here we introduce a new version of TrackMate rewritten to improve performance, usability, and versatility. In particular, we developed a new API that allowed integrating the main segmentation tools based on ML and DL algorithms available in Java (Figure 1). For instance, ilastik [20], Weka [21] and StarDist [22] are now integrated into TrackMate as object detectors allowing the user to seamlessly use these methods from the same interface. We also integrated the morphological segmentation algorithm of MorphoLibJ [23]. For algorithms not implemented in Java, TrackMate can incorporate their segmentation results by directly loading label images, probability maps, or binary mask images. Importantly, the integrated segmentation algorithms can provide the shape of the object in addition to its position in every single frame. We therefore rewrote the TrackMate data model to store, display and analyze the object contours in 2D. TrackMate can now measure morphological features on the tracked objects. This allows for instance to correlate motility with object shape changes and fluorescent intensity over time.

**Fig. 1.**
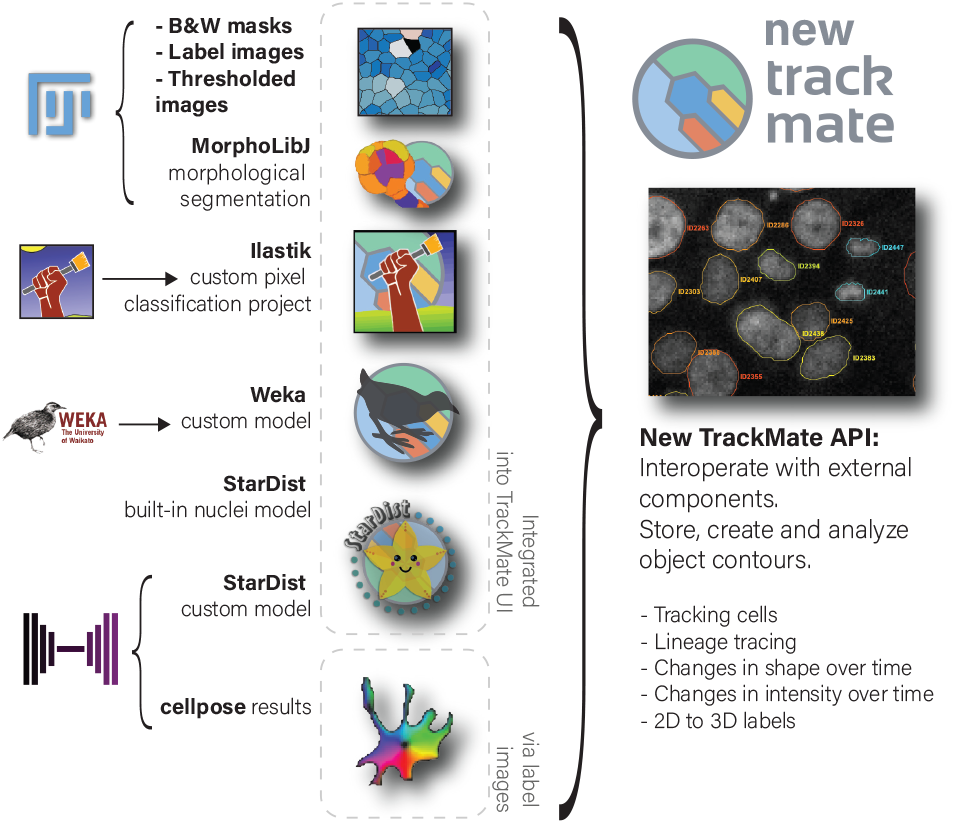
New TrackMate capabilities. TrackMate can now create, use, analyze and store object contours segmented from 2D images. These contours enable TrackMate to extract morphological features of the tracked objects over time. We also wrote a new application programming interface (API) to facilitate integrating external components to TrackMate. We use this API to incorporate popular segmentation tools available in Java, including ilastik, the Weka Trainable-Segmentation Fiji plugin, StarDist, and the morphological segmentation tool MorphoLibJ. TrackMate can now also track previously segmented objects by directly importing mask or label images generated, for instance, using cellpose.

We found that the new detectors offer better tracking accuracy than the LoG detector when the shape of the objects to track is more complex than a round spot (Supplementary Note 1, Supplementary Figure 1-3). Significantly the newly included detectors considerably increase the breadth of TrackMate applications and capabilities (Figure 2, Movie 1-10, Supplementary Figure 4-7, Supplementary manual, and tutorials).

**Fig. 2.**
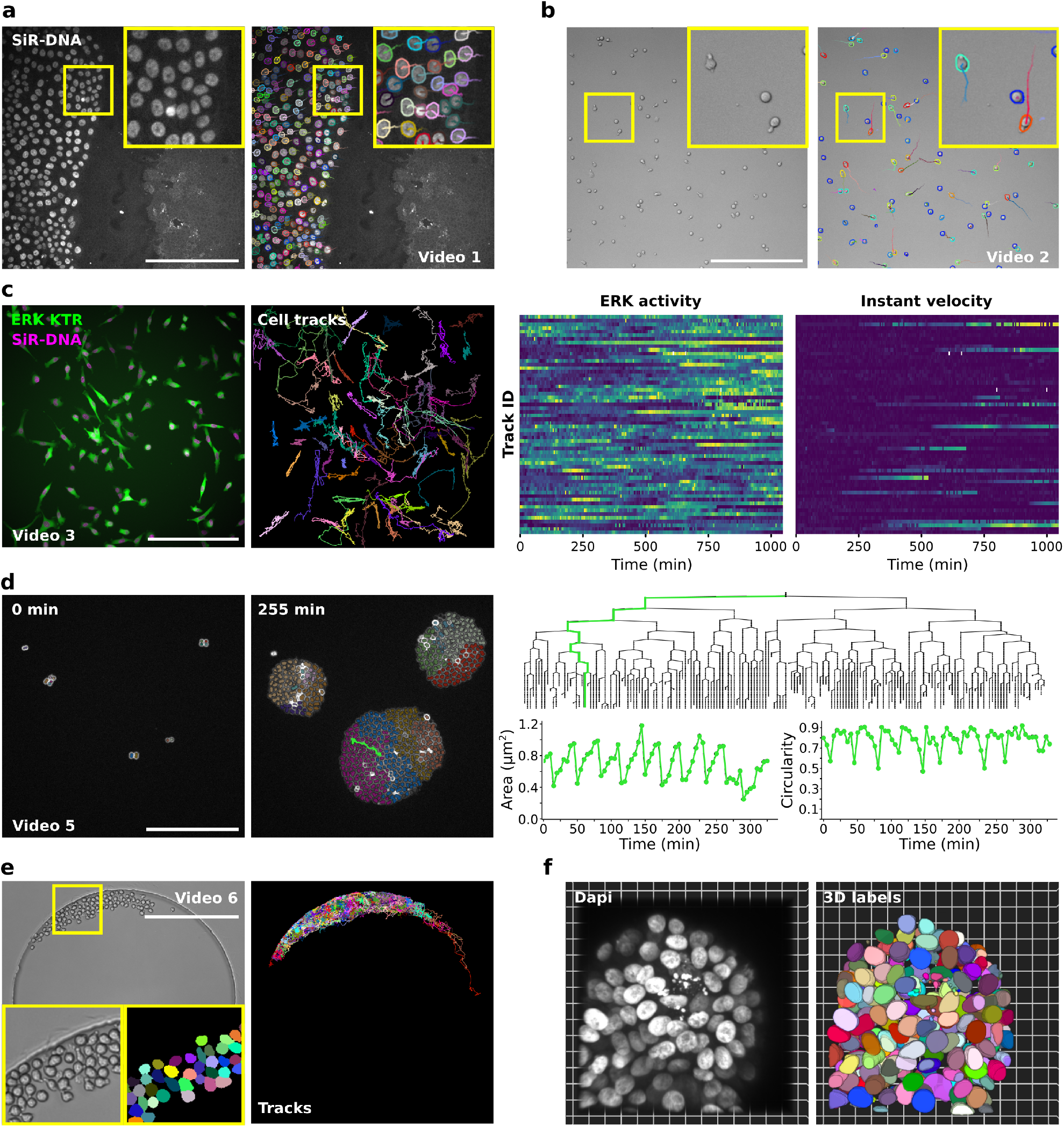
TrackMate can be used to track objects from a wide variety of bio-imaging experiments. **a**. Migration of MCF10DCIS.com cells, labeled with SiR-DNA, recorded using a spinning disk confocal microscope and automatically tracked using a custom StarDist model loaded in TrackMate (see also Movie 1). Detected cells and their local tracks (colors indicate track ID) are displayed. Scale bar = 250 μm. **b**. The migration of activated T cells plated on ICAM-1 was recorded using a brightfield microscope and automatically tracked using a custom StarDist model loaded in TrackMate (see also Movie 2). Detected cells (colors indicate the mean track speed; blue slow-moving cells, red fast-moving cells) and their local tracks (colors indicate track ID) are displayed. Scale bar = 250 μm. **c**. MDA-MB-231 cells stably expressing an ERK activity reporter (ERKKTRClover) and labeled using SiR-DNA were recorded live using a wide-field fluorescent microscope over 17 hours. Cell nuclei were automatically tracked over time using a StarDist model available in TrackMate (see Movie 3). For each tracked cell, the average intensity of the ERK reporter was measured in their nucleus over time (directly in TrackMate). Changes in ERK activity and in instant velocity are displayed as heatmaps (blue low, yellow high). **d**. The growth of *Neisseria meningitidis* expressing PilQ-mCherry was recorded using a spinning-disk confocal microscope. An ilastik pixel classifier trained to segment individual bacteria was loaded into TrackMate to follow bacteria growth. Representative field of views and the linage tree of the bacteria highlighted in green are displayed (see Movie 5). Changes in area and circularity of a bacterium over the tracking period are also highlighted (green track). Cell division events translate in sharp decreases in area, followed by a quasi-linear increase. The circularity roughly plateaus during cell growth then decreases before cell division. Scale bar = 25 μm. **e**. Mouse hematopoietic stem cells migrating in a hydrogel microwell were automatically segmented using cellpose implemented in the ZeroCostDL4Mic platform. The resulting label images were automatically tracked using TrackMate (see Supplementary Figure 1 and Movie 6). Example raw and label images, as well as cell tracks, are displayed. **f**. MCF10DCIS.com 3D spheroids were stained for Dapi and imaged using a spinning disk confocal microscope. Across the Z volume, nuclei were detected at each Z plane using StarDist and tracked (everything done in TrackMate). Tracked nuclei were then exported as a label image to create 3D labels (see Movie 8).

For instance, the StarDist integration offers efficient and versatile nuclei detection in fluorescence images via the built-in model (from image set BBBC038v1 in [24]). Our integration also provides an interface to use custom models trained *e.g*. with the ZeroCostDL4Mic platform [25]. To illustrate this, we used custom StarDist models to track fluorescently labelled nuclei of collectively migrating breast cancer cells or track rapidly migrating T cells from brightfield images (Figure 2a-b and Movie 1-2). Before this integration, fully automated tracking of label-free cells was difficult in TrackMate.

Thanks to TrackMate supporting multi-dimensional images, users can now track objects using one channel and measure changes of intensities within the track objects in separate channels over the tracking period. As an example, we tracked the nuclei of breast cancer cells expressing an ERK activity reporter and followed changes in ERK activity in single cells as they migrate (Figure 2c, Supplementary Figure 4 and Movie 3).

Users can also train a custom ML classifier using the Fiji Weka Trainable-Segmentation plugin or ilastik and use them subsequently in TrackMate. For instance, we used Weka combined with TrackMate to track focal adhesions in endothelial cells (Supplementary Figure 5 and Movie 4). We also used an ilastik pixel classifier to follow *Neisseria meningitidis* growth and correlate lineage information to single bacteria morphological measurements (Figure 2d and Movie 5). This new version of TrackMate makes it possible to track objects segmented by an extensive range of segmentation algorithms. Indeed TrackMate can now import binary mask images, segmentation probability maps, or label images directly, then track the imported objects. As examples, we tracked migrating cancer cells (fluorescent images), and hematopoietic stem cells ([26], brightfield images, Figure 2e, Supplementary Figure 6 and Movie 6-7) previously segmented using cellpose [27].

The new detectors work for 2D and 3D images when possible, but the display and analysis of object contours are currently limited to 2D images. However we can exploit TrackMate to segment 3D objects, using a slice-by-slice approach. Instead of tracking objects over time, we tracked them across subsequent axial planes and linked them to recover 3D objects (Figure 2f, Supplementary Figure 7 and Movie 8-10). With this approach, TrackMate can accelerate the segmentation of challenging 3D datasets by using a custom DL model trained on 2D images, for instance, using interactive annotation tools such as Kaibu [28].

We provide a detailed manual and step-by-step tutorials to facilitate the use of the new TrackMate detectors (Supplementary manual and Supplementary Figure 8). We also provide documentation aimed for developers to integrate their own DL and ML algorithms in TrackMate. Altogether, TrackMate now enables powerful segmentation approaches for tracking purposes directly in Fiji within a user interface already familiar to many. We envision that by making complex tracking problems more easily solvable by scientists, this new version of TrackMate will accelerate discoveries made in Life-Sciences.

## Supporting information

Supplemental manual

## Online methods and data availability statement

The version of TrackMate described here is available in the Fiji [12] software by simply updating it. TrackMate is documented on the ImageJ wiki: https://imagej.net/plugins/trackmate/ and the documentation for the new features can be accessed from https://imagej.net/plugins/trackmate/trackmate-v7-detectors. We also provide 11 test datasets that are made available via a dedicated Zenodo collection [29].

## Author contributions

GJ and JYT conceived the project; JYT wrote source code; GJ, JWP, NHR, and LLB performed the image acquisition of the test and example data; GJ, JWP, RFL, JYT, MSP, DE and SUR tested the code; JRWC, DB, GD and ACO provided critical reagents; GJ, JWP, JYT, MSP, DE, SUR and JYT redacted the documentation and tutorials.; GJ and JYT wrote the manuscript with input from all co-authors.

## Declaration of Interests

The authors declare no competing interests.

## Acknowledgments

The integration of existing algorithms as new detectors in TrackMate has been made possible thanks to the high quality of the code, documentation, and support provided by their respective authors. In particular, we would like to thank Anna Kreshuk, David Legland, Dominik Kutra, Ignacio Arganda-Carreras, Martin Weigert, Siân Culley, and Uwe Schmidt. We can only hope for TrackMate to reach such a standard of quality to become a better tool of Science. We are also grateful for the support and help of the bioimage analysis community, in particular Curtis Rueden, Jan Eglinger, Nicolas Chiaruttini, Tobias Pietzsch and Pavel Tomancak. We thanks Helen Blau for giving us the permission to use the “Mouse hematopoietic stem cells in hydrogel microwells” dataset made available on the Cell Tracking Challenge website.

This study was supported by France BioImaging (Investissement d’Avenir; ANR-10-INBS-04, JYT), the Academy of Finland (GJ), the Sigrid Juselius Foundation (GJ), the Cancer Society of Finland (GJ), the Åbo Akademi University Research Foundation (GJ, CoE CellMech), the Drug Discovery and Diagnostics strategic funding to Åbo Akademi University (GJ) and the European Union’s Horizon 2020 research and innovation program under Marie Sklodowska-Curie grant agreement 841973 (JRWC). JWP was supported by Health Campus Turku 2.0 funded by the Academy of Finland. RFL was supported by an MRC Skills development fellowship (MR/T027924/1). The Cell Imaging and Cytometry Core facility (Turku Bioscience, University of Turku, Åbo Akademi University and Biocenter Finland) is acknowledged for services, instrumentation, and expertise.

## Notes

https://zenodo.org/communities/trackmate/

https://github.com/fiji/TrackMate

https://imagej.net/plugins/trackmate/trackmate-v7-detectors

### Competing Interest Statement

The authors have declared no competing interest.

### Summary of Updates

- Add a supplementary figure guiding the user in the choice of detectors. - Fix the title grammar. - Add a missing affiliation for LLB.

