## Supplemental information for "Bringing TrackMate into the era of machine-learning and deep-learning"

### Supplementary Information for: Bringing TrackMate into the era of machine-learning and deep-learning.

[1]Image Analysis Hub, C2RT / DTPS, Institut Pasteur, Paris, FR
[2]Biostatistics and Bioinformatic Hub, Department of Computational Biology, Institut Pasteur, Paris, FR
[3]Turku Bioscience Centre, University of Turku and Åbo Akademi University, Turku, FI
[4]Åbo Akademi University, Faculty of Science and Engineering, Biosciences, Turku, FI
[5]Turku Bioimaging, University of Turku and Åbo Akademi University, Turku, Finland
[6]Pathogenesis of Vascular Infections unit, INSERM, Institut Pasteur, Paris, FR
[7]Université de Paris, 75006, Paris, FR
[8]MRC Laboratory for Molecular Cell Biology, University College London, London, UK
[9]The Francis Crick Institute, London, UK
[10]Department of Microbiology and Immunology, SUNY Upstate Medical University, Syracuse NY, USA
[ca]Current address: Micrographia Bio, Translation and Innovation Hub 84 Wood Lane, London, UK
[*]Equal contributors, authors listed alphabetically

##### List of Tables

Table S1. Tracking parameters used in the performance assessment . . . . 4

### Movie Legends.

***Movie 1 - Using StarDist within TrackMate to track migrating cancer cells.*** MCF10DCIS.com cells, labelled with Sir-DNA, were recorded using a spinning-disk confocal microscope and automatically tracked using StarDist integrated within TrackMate. Detected nuclei and local tracks are displayed. Color indicates ID.

***Movie 2 - Using StarDist within TrackMate to track migrating T cells.*** Activated T cell plated ICAM-1 were recorded using a brightfield microscope and automatically tracked using StarDist integrated within TrackMate. Color indicates mean speed.

***Movie 3 - Measuring ERK activity in migrating cancer cells.*** MDA-MB-231 cells expressing ERK-KTR-GFP and labelled with Sir-DNA, were recorded using a widefield microscope and automatically tracked using StarDist integrated within TrackMate. Only tracks remaining in the field of view over the whole duration of the movie are displayed. Color indicates ID.

***Movie 4 - Using Weka within TrackMate to track focal adhesions.*** Endothelial cells expressing Paxillin-GFP were recorded live using a spinning-disk confocal microscope. A custom Weka pixel classifier trained to segment focal adhesion was then loaded directly into TrackMate to track focal adhesions. In the middle panel, focal adhesions are color-coded to indicate their lifetime (red, long-lived, blue short-lived). In the right panel, track colors indicate ID.

***Movie 5 - Using ilastik within TrackMate to follow bacteria growth.*** The growth of *Neisseria meningitidis* expressing PilQ-mCherry was recorded using a spinning-disk confocal microscope. An ilastik pixel classifier trained to segment individual bacterium was loaded directly into TrackMate to follow bacteria growth.

***Movie 6 - Using cellpose and TrackMate to track stem cells.*** Mouse hematopoietic stem cells migrating in a hydrogel microwell were automatically segmented using cellpose. The resulting label images were tracked using TrackMate. In the bottom left panel, the color of the object indicates the distance travelled (red longest distance, blue shortest distance). In the bottom right panel, track colors indicate ID.

***Movie 7 - Using cellpose and TrackMate to track migrating cancer cells.*** MCF10DCIS.com cells expressing lifeact-RFP, labelled with Sir-DNA, were recorded using a spinning-disk confocal microscope. Cells were segmented using cellpose, and label images were tracked using TrackMate. Color indicates ID.

***Movie 8 - Using StarDist 2D whithin TrackMate to generate 3D labels.*** MCF10 DCIS.com spheroids were imaged using a spinning-disk confocal microscope. To generate 3D labels, nuclei were detected and tracked across the Z volume using StarDist implemented in TrackMate. The 3D rendering was performed using Arivis Vision4D.

***Movie 9 - Using cellpose 2D and TrackMate to segment 3D images of Arabidopsis thaliana floral meristem.*** Confocal images of *Arabidopsis thaliana* floral meristem were segmented using cellpose 2D implemented in ZeroCostDL4Mic. TrackMate was used to track the 2D labels across the Z volume and generate 3D labels. Arivis Vision4D was used to perform the 3D rendering.

***Movie 10 - Using cellpose 2D and TrackMate to segment 3D images of a developing Drosophila melanogaster embryo.*** Light-sheet microscopy images of a developing *Drosophila melanogaster* embryo were segmented using cellpose (2D). TrackMate was then used to track the 2D labels across the Z volume and generate 3D labels. Arivis Vision4D was used to perform the 3D rendering.

### Supplementary notes.

#### Supplementary Note 1: Tracking performance measurements.

**Introduction.** We wanted to assess whether using a detector based on Deep-Learning has a positive impact on the tracking accuracy of TrackMate. Two main frameworks can be used to assess tracking performance: the single-particle tracking challenge [1, 2] and the cell tracking challenge [3, 4]. The cell tracking challenge deals with cell image data that are very well suited for the new detectors shipped in this latest version of TrackMate. Yet, we chose to focus on the single-particle tracking (SPT) challenge data and metrics for this assessment. Indeed, the SPT challenge relies on using simulated images with varying object types, motility types, image quality, and object density. Four scenarios simulate each the motility of a different subcellular organelle: microtubule tips, vesicles, viruses, and membrane receptors. Each of these objects type has its own motility type, respectively: directed motion, random walks, and switching between these two modes. For each scenario, several movies are available with varying signal-to-noise ratios (SNR) and particle density (12 to 15 movies per scenario). This dataset will help us uncover the range of image quality or object density a detection algorithm primes over another one. However, the objects simulated in the SPT challenge are subcellular organelles: These objects are close to being sub-resolved and, therefore, mostly shapeless. For this type of object that resembles Gaussian spots, the detectors based on the Laplacian-of-Gaussian (LoG) filter are proven to be the best, especially when images are corrupted by white noise [5]. A LoG detector was already present in the previous version of TrackMate, and it is the one we will use in our comparison against a new detector based on the StarDist Deep-Learning algorithm [6]. As the data we analyze are ideal for the LoG detector, we expect these comparison settings to favor the LoG detector.

##### Methods.

***Generating a spot detector based on the StarDist algorithm.*** Particle detection algorithms started with classical computer vision (CV) approaches [7]. While they excel for well defined distinct blob-like particles, their performance was often found unsatisfactory in the conditions of low SNR, high particle density, and more complex particle geometry. The rise of DL in computer vision brought numerous new approaches in the last few years focused on particle tracking. DL approaches now readily outperform classical algorithms in conditions of low noise, unsteady illumination and heterogeneous geometry [8, 9], high-density, complex interference patterns in 3D [10], single-molecule localization [11], microtubule tracking [12], virus particles in challenging intracellular environment [13], dense particles in 3D with anisotropic PSF [14].

The generation of an efficient Deep-Learning based detector for single-particle represents a very significant work that would build upon the literature cited above. This is, however, not our purpose here. Instead, for our assessment, we created a particle detection method based on StarDist, as this algorithm is present in the last version of TrackMate. While StarDist was not directly created to detect single particles, we aimed to investigate how using StarDist to resolve overlapping objects (common at high density) and its robustness against noise would impact tracking performance. Also, using StarDist, we can build a single model that can harness all of the object types present in the SPT Challenge and more. To build this model, we generated a training dataset using the ISBI Challenge benchmark generator [15] in Icy [16].

The generated benchmark data contains simulated movies of the four scenarios of the single-particle tracking Challenge. We included movies encompassing multiple SNR and several object densities. We used the associated ground-truth images to create mask images where each object is represented by a circular spot of diameter 5 pixels. We then trained a StarDist model using the simulated images and these masks. This custom StarDist model was trained for 200 epochs on 4800 paired image patches (image dimensions: 512×512, patch size: 512×512 with a batch size of 4 and a mae loss function, using a custom StarDist 2D python script. This model was then used along with the *StarDist custom model* detector of TrackMate for the performance assessment. The generated model, the scripts to generate it and the training details are available online [17]. Importantly, the dataset used for the performance assessment is the one used in [1] while we trained the StarDist model on a different, newly simulated dataset using [15].

***The LoG detector.*** For the comparison, we used the LoG detector present in TrackMate since its first release [18]. This detector operates by filtering the source image using the LoG filter in the Fourier space then inverting it so that the object locations show as bright spots in the filtered image. The LoG filter is configured with a $\sigma$ value matching the object diameter. The objects are detected in the filtered image by looking for local maxima. If the filtered value of a spot is below a threshold set by the user, it is deemed spurious and pruned from the list of detections.

***ISBI SPT Challenge metrics.*** The ISBI SPT Challenge ships a specific evaluation dataset against which candidate tracking algorithms have been evaluated [2]. It also defines five metrics used to measure the performance of these tracking algorithms. Here, we sought to reproduce Figure 2 from [1] by comparing TrackMate LoG and StarDist detectors. We plotted three of the five metrics used in the ISBI SPT Challenge. They are described in detail in [1], but we summarize them here. **1.** The $\alpha$ value is a measure of the distance from the candidate tracks to the ground truth tracks, ignoring spurious tracks generated by the tracking algorithm that are absent in the ground truth data. Its values range from 0 to 1; higher is better, reaching 1 when the candidate track are exactly aligned to the ground-truth tracks. The $\alpha$ value decreases as the particles in the candidate tracks are found further away from the ground-truth track or missed. **2.** The $\beta$

| Scenario | LoG detector parameters | StarDist detector parameters | Linking parameters |
|---|---|---|---|
| MICROTUBULE | LoG detector; *Radius* = 5 pixels, *Threshold* = 0.30, *Sub-pixel localization* = true, *Median filtering* = false | NA | Kalman tracker; *Search radius* = 8 pixels; *Max linking distance* = 8 pixels; *Max frame gap* = 1 |
| RECEPTOR | LoG detector; *Radius* = 2.5 pixels, *Threshold* = 2.62, *Sub-pixel localization* = true, *Median filtering* = false | NA | Kalman tracker; *Search radius* = 8 pixels; *Max linking distance* = 8 pixels; *Max frame gap* = 2 |
| VESICLE | LoG detector; *Radius* = 2.5 pixels, *Threshold* = 3.02, *Sub-pixel localization* = true, *Median filtering* = false | NA | Simple LAP tracker; *Max linking distance* = 8 pixels; *Max frame gap* = 2 |

**Table S1.** Tracking parameters used in the performance assessment.

value builds upon α but includes a penalty for spurious tracks. It ranges from 0 to α; higher is better. **3.** The RMSE measures the root-mean-square error of the position of candidate objects with respect to the ground-truth positions. It is a positive scalar value, lower is better, and zero indicates a perfect match.

We adapted the existing code provided by [1] to batch computes these three metrics in a multi-threaded fashion and without any dependencies. Our modified code is available online here [19].

***Evaluating the two detectors.*** Because the Java version of StarDist only deals with 2D images, we limited ourselves to the 2D scenarios of the SPT Challenge, namely the RECEPTOR, VESICLE and MICROTUBULE scenarios. Also, because we wanted to assess the impact of a new detector on the overall tracking performance, we used the same particle linking algorithm with identical parameters in both cases. We first ran a systematic parameter sweep to find the optimal linking algorithm and parameter set. Instead of taking the optimal linking parameters for individual images in the test dataset, we retained a common detection and linking parameter set that yields overall good tracking results over a whole scenario. This approach leads to somewhat sub-optimal absolute performance but produces results that can be used to compare the two detectors. The assessment parameters are listed in the Table S1. The code that performs tracking in batch over the benchmarking dataset and imports the results of the StarDist detection is available publicly in a branch on the GitHub repository of TrackMate [20].

**Results and Discussion.** For the RECEPTOR scenario, we observe that, at low density, the LoG detector performance slightly exceeds that of the StarDist detector (Figure S1). This is expected, as for this scenario, the objects to detect closely resemble Gaussian spots, rending the LoG detector optimal for the task [5]. As the density of objects increases from 100 ('low' density) to 500 ('mid') and 1000 ('high') per frame, the performance of the StarDist detector overpasses that of the LoG detector, albeit only by a marginal value. Except for the 'mid' density and SNR of 2 case where the StarDist detector significantly outperforms the LoG detector, the two detectors perform similarly. We observed the same behavior in the VESICLE scenario (Figure S2). Again, this is expected as the object to track in the VESICLE scenario

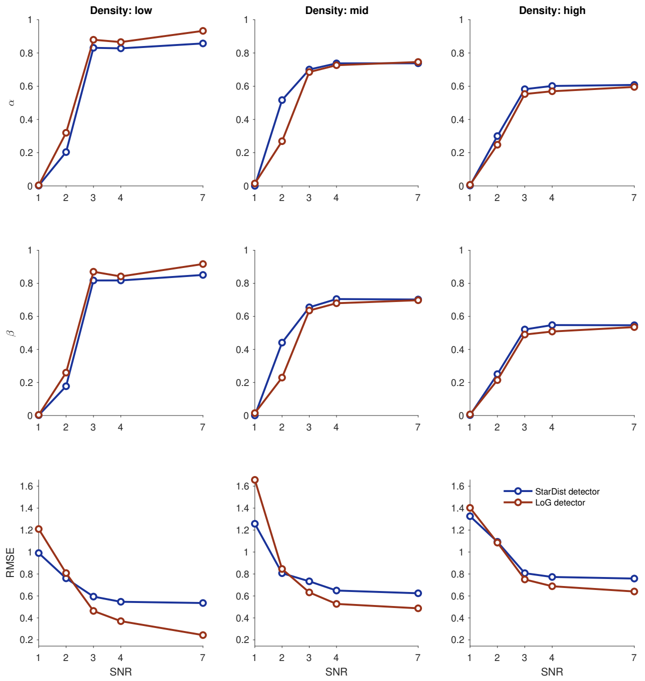

**Figure S1. Results of the tracking performance assessment for the RECEPTOR scenario.** From the left to right column, the density of particles increases from 100 to 500 and 1000 objects per frame. From top to bottom, three of the performance metrics of the ISBI SPT challenge. In each plot, the Y-axis plots the value of the metrics. For α and β, higher is better. For RMSE, lower is better (see text for the description of the metric). The X-axis plots the signal-to-noise ratio (SNR) value for individual movies of one condition. In blue: results for the new StarDist based detector. In red: results for the classical LoG detector.

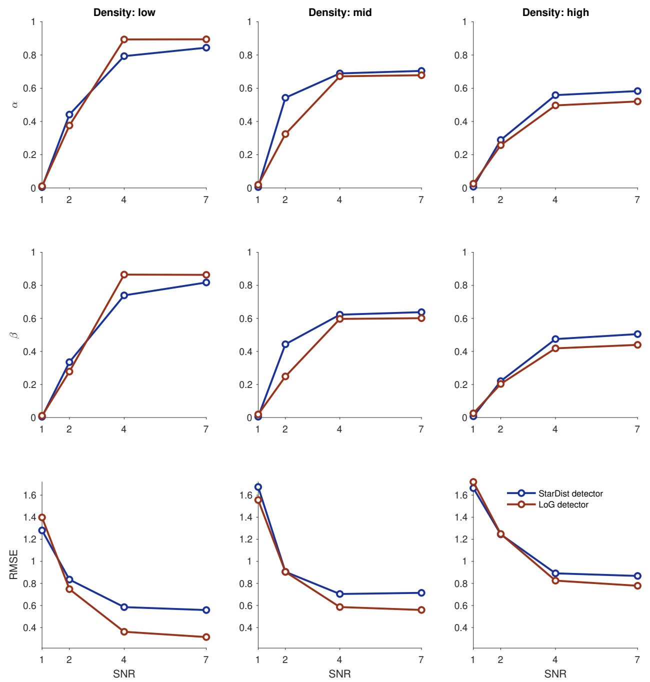

**Figure S2. Results of the tracking performance assessment for the `VESICLE` scenario.**

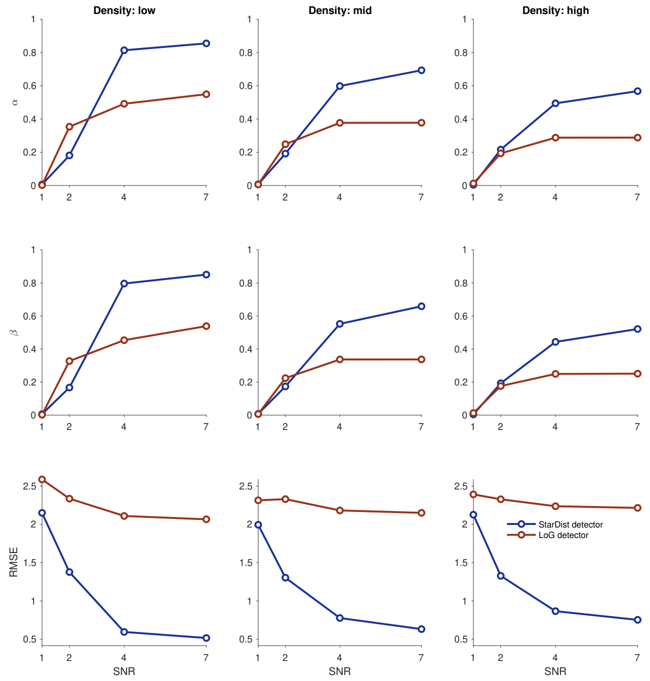

**Figure S3. Results of the tracking performance assessment for the `MICROTUBULE` scenario.**

also resembles Gaussian spots. We conclude that a detector based on StarDist can perform similarly to the LoG detector for sub-resolved objects. Interestingly, the StarDist-based detector slightly exceeds the performance of the LoG detector when the object density increases, a feature we attribute to the ability of the StarDist algorithm to resolve overlapping objects.

In the case of the `MICROTUBULE` scenario, the differences in performance are more drastic (Figure S3). Indeed, in this scenario, the StarDist detector surpasses the LoG detector by a large margin in most cases. Interestingly, this is the scenario where the objects to track are slightly more complex than a Gaussian spot as they simulate the dynamic of microtubule plus tips, and the objects to track resemble small comets with a tail protruding backward. The LoG detector offers better performance than the StarDist detector only for images with an SNR$\leq$ 2 value and with objects at 'low' and 'mid' densities. But this advantage diminishes as the density increases for this particular SNR value.

This assessment concludes that DL-based detectors positively impact tracking performance against classical detectors, in the case of complex objects, even when the comparison includes identical linking algorithms. The StarDist detector we built for this assessment is simple compared to what others have described [8–14]. That said, this simple comparison allows us to advise users to select the StarDist detector to track their object of interest when a suitable segmentation model is available. Using the StarDist detector is especially advantageous as soon as the objects to track are more complex than a Gaussian spot and their density is large.

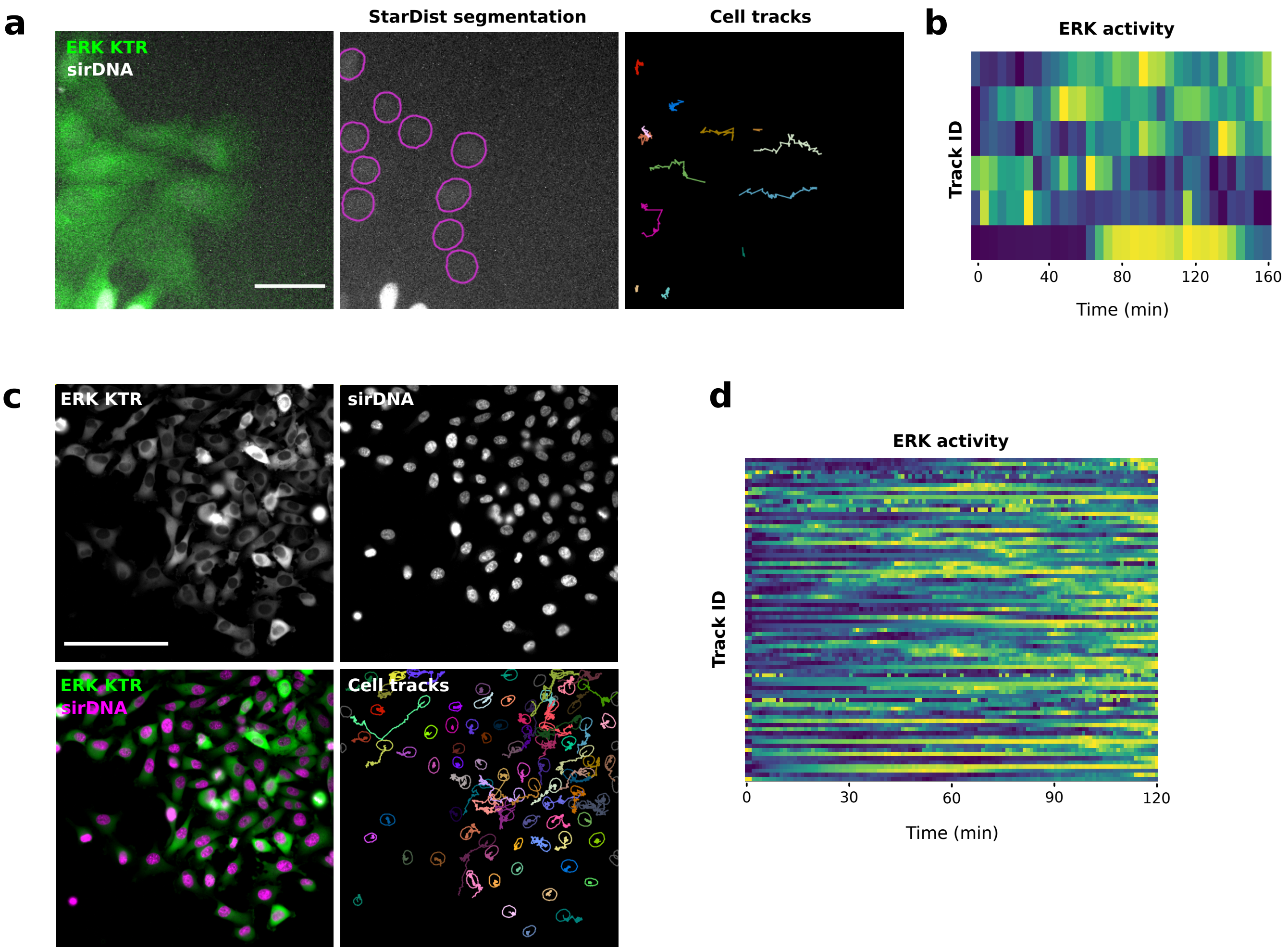

**Figure S4. Following ERK activity in migrating cells.** U2OS (**a.** and **b.**) and MDA-MB-231 cells (**c.** and **d.**) stably expressing an ERK activity reporter (ERK-KTR-Clover) and labeled using SiR-DNA were recorded live using a widefield fluorescent microscope. U2OS cells were recorded live over 3 hours (1 image every 5 minutes) and MDA-MB-231 cells were recorded live over 2 hours (1 image every minute). Cell nuclei were automatically tracked over time by using StarDist in TrackMate. A custom StarDist model was trained to detect the U2OS nuclei using the ZeroCostDL4Mic platform. The "Versatile fluorescent nuclei" StarDist model was used to track the MDA-MB-231 cell nuclei. For each tracked cell, the average intensity of the ERK reporter was measured in their nucleus over time (directly in TrackMate). Changes in ERK activity are displayed as heatmaps (blue low, yellow high). Heatmaps were generated using PlotTwist. Scale bar = 250 µm.

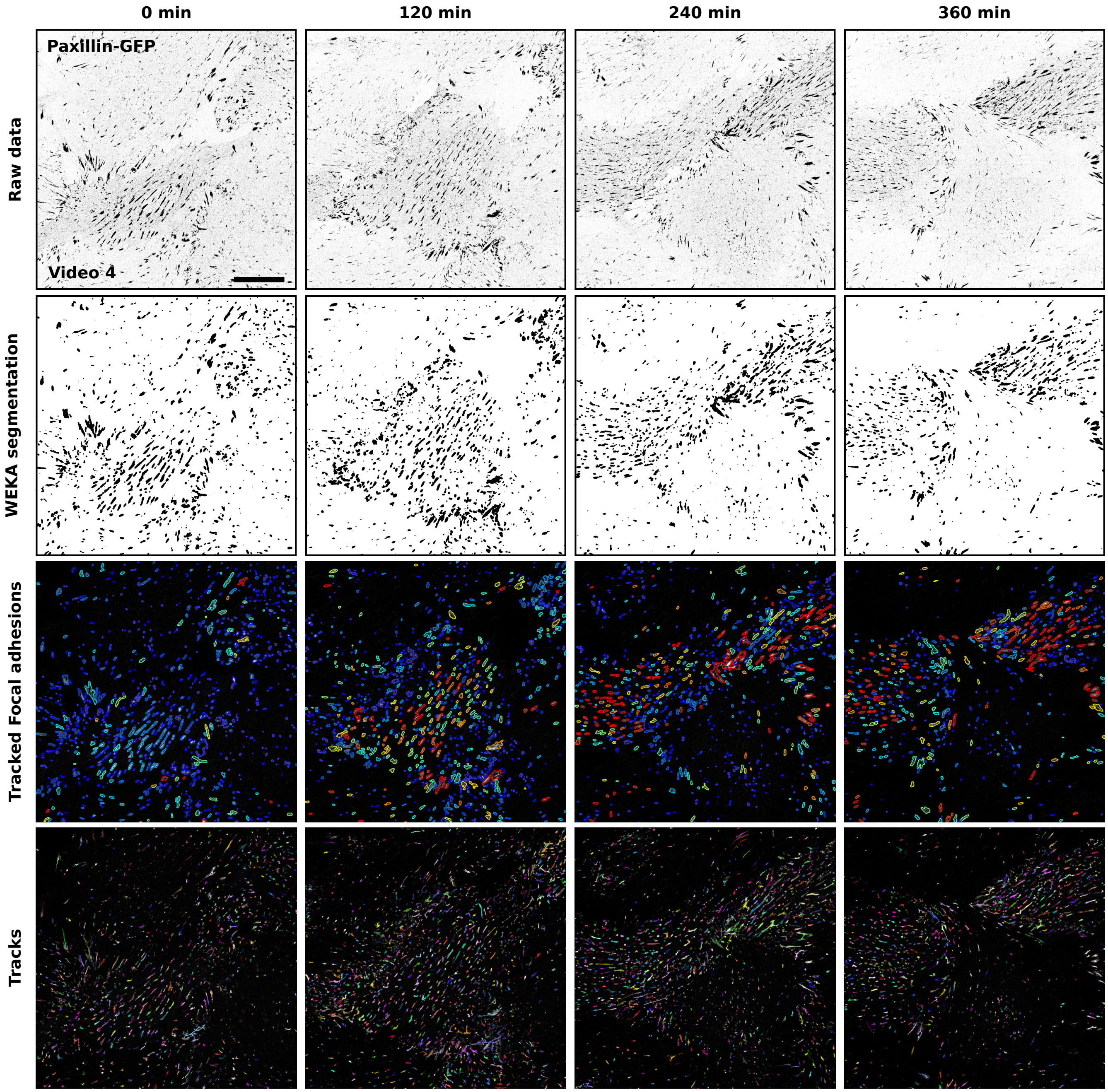

**Figure S5. Tracking focal adhesions using Weka and TrackMate.** Endothelial cells expressing paxillin-GFP were recorded live using a spinning disk confocal microscope. Focal adhesions were then segmented and tracked using Weka integrated within TrackMate (Movie 4). Raw data (inverted LUT), Weka segmentation results, tracked focal adhesion, and the focal adhesion tracks are displayed for selected time points. Tracked focal adhesions are color-coded to indicate their lifetime (red, long-lived, blue short-lived). In the bottom panel, track colors indicate ID. Scale bar = 25 µm.

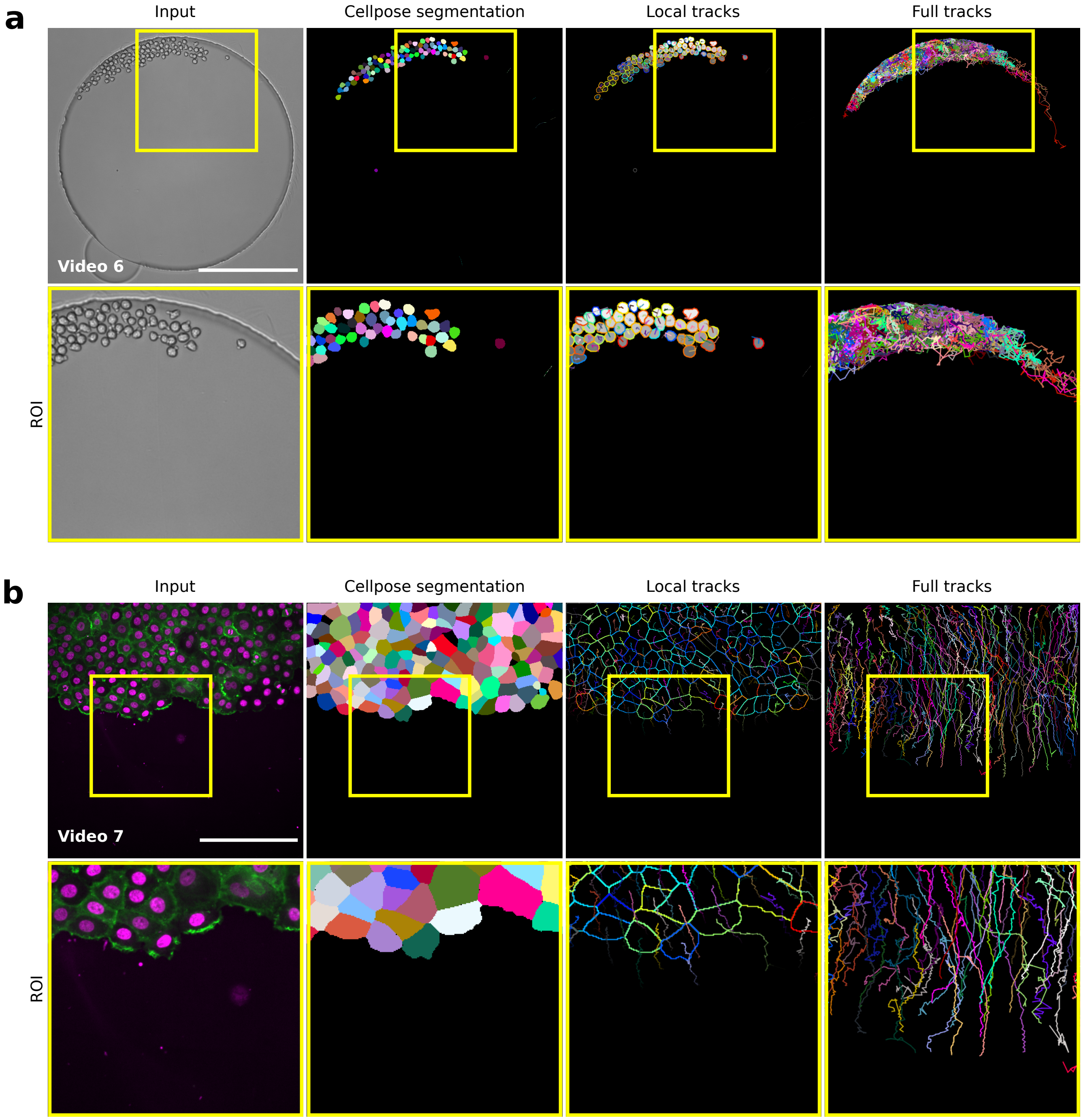

**Figure S6. Tracking label images using TrackMate. a.** Mouse hematopoietic stem cells migrating in a hydrogel microwell were automatically segmented using cellpose (Cyto model) implemented in the ZeroCostDL4Mic platform. The resulting label images were automatically tracked using TrackMate (Movie 6). Example raw and label images as well as local and full cell tracks are displayed. Yellow squares highlight regions of interest that are magnified. Scale bar = 250 µm. This dataset is available from the Cell Tracking Challenge. **b.** MCF10DCIS.com cells stably expressing lifeact-RFP and labeled with SiR-DNA were recorded live using a spinning disk confocal microscope. Cells were segmented using cellpose (Cyto model) implemented in the ZeroCostDL4Mic platform. The resulting label images were tracked using TrackMate (Movie 7). Example raw and label images as well as local and full cell tracks are displayed. Yellow squares highlight regions of interest that are magnified. Scale bar = 250 µm.

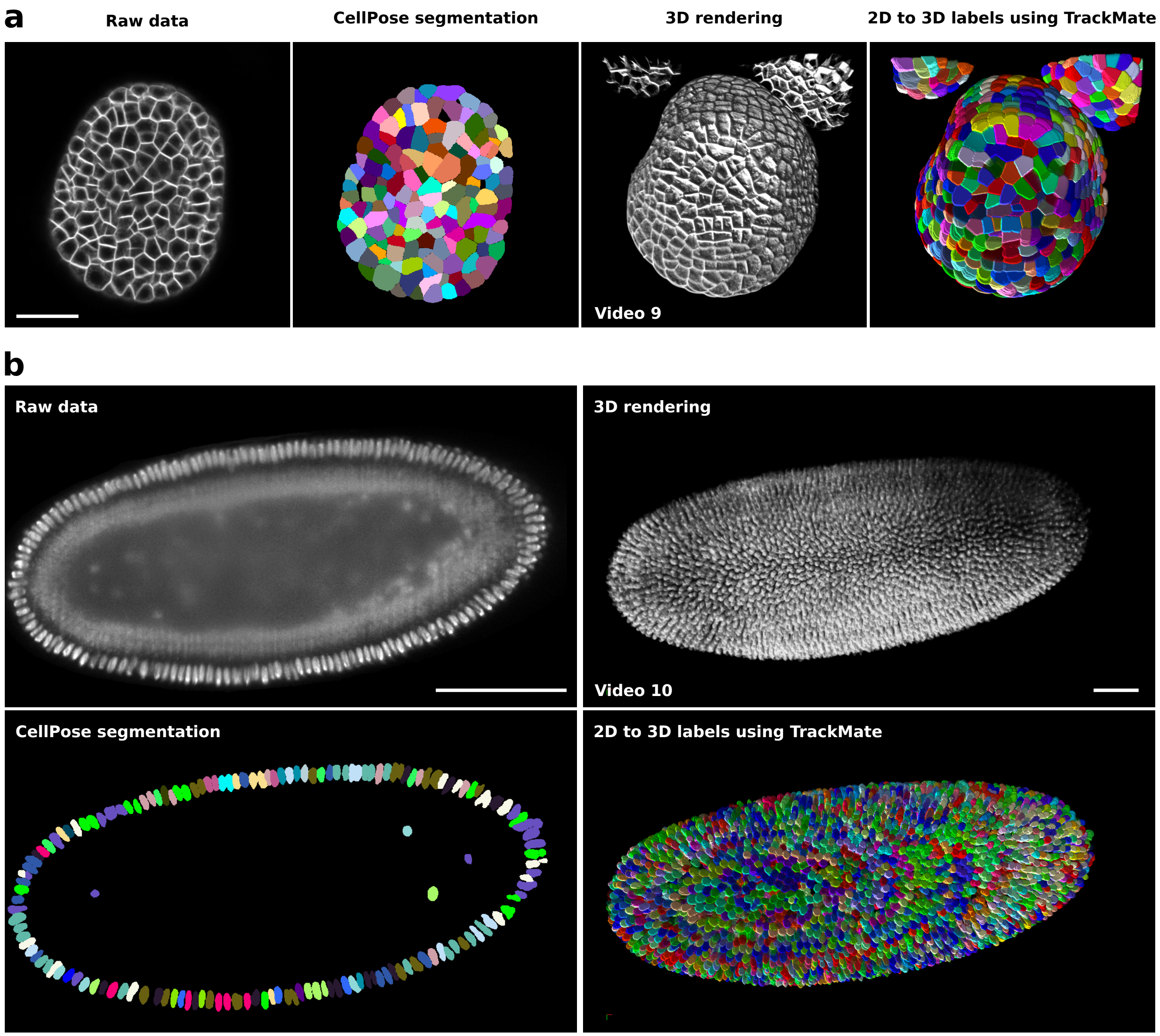

**Figure S7. Tracking 2D labels to generate 3D labels using TrackMate.** Tracking 2D labels to generate 3D labels using TrackMate. **a.** Confocal images of *Arabidopsis thaliana* floral meristem [21, 22] and **b.** light-sheet images of a developing *Drosophila melanogaster* embryo [3, 4, 23] were automatically segmented using cellpose 2D (Cyto2 model) implemented in the ZeroCostDL4Mic platform [24, 25]. Representative single Z plane and the corresponding cellpose segmentation results are displayed. To generate 3D labels, cellpose 2D segmentation results were then tracked using TrackMate. 3D rendering of the raw data and of the 3D segmentation results are also shown. Scale bars: (a) = 25 µm, (b) = 100 µm.

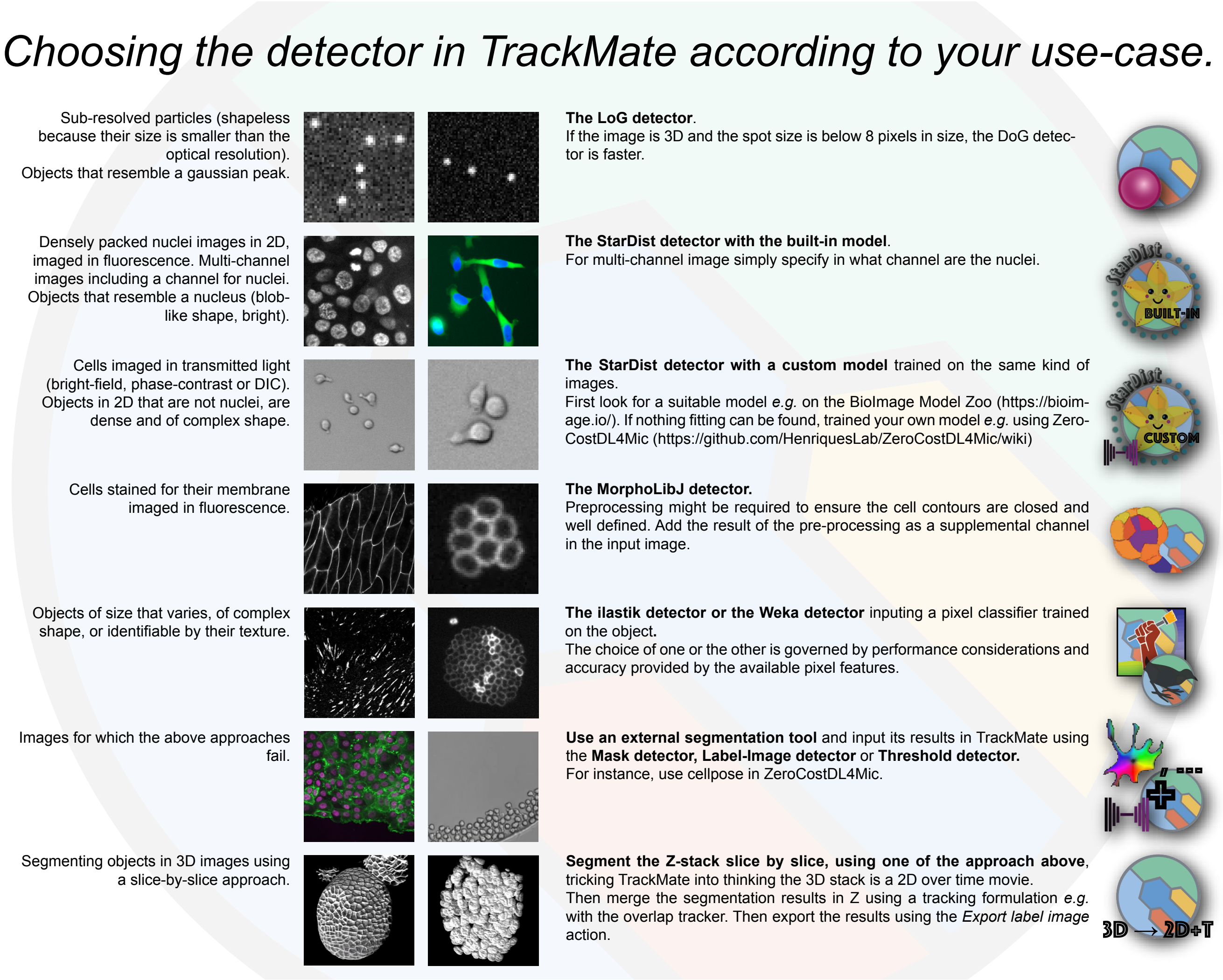

**Figure S8. Choosing a TrackMate detector according to the input image.**

### Online Methods.

#### Cells and reagents.

MDA-MB-231 and U2OS cells were engineered to express the Erk KTR by first producing lentiviral particles in HEK 293FT packaging cells (ThermoFisher, R70007). Cells were co-transfected with the third generation lentiviral packaging system composed of pMDLg / pRRE (Addgene plasmid 12251), pRSV-Rev (Addgene plasmid 12253), pMD2.G (Addgene plasmid 12259), along with the pLentiPGK Puro DEST ERK KTRClover (a kind gift from Markus Covert; Addgene plasmid 90227) transfer plasmid, using Lipofectamine 3000 (ThermoFisher) in OptiMEM (Gibco, 31985070), as per the manufacturer's protocol [26, 27]. After 24 hours the media was changed for complete growth medium and incubated for a further 24 hours, at which point the media was collected and filtered through a 0.45 µm syringe filter. MDA-MB-231 and U2OS cells were transduced with lentivirus for 48 hours in the presence of polybrene (8 µg/ml; Sigma, TR-1003-G), before washing and selection of stable positive cells using puromycin (2 µg/ml). Cells were then sorted by fluorescence-activated cell sorting (FACS) to isolate a population within a similar fluorescence range. MCF10 DCIS.COM cells were cultured in a 1:1 mix of DMEM (Sigma-Aldrich) and F12 (Sigma-Aldrich) supplemented with 5% horse serum (16050-122; Gibco BRL), 20 ng/ml human EGF (E9644; Sigma-Aldrich), 0.5 mg/ml hydrocortisone (H0888-1G; Sigma-Aldrich), 100 ng/ml cholera toxin (C80521MG; Sigma-Aldrich), 10 µg/ml insulin (I9278-5ML; Sigma-Aldrich), and 1% (vol / vol) penicillin / streptomycin (P0781-100ML; Sigma-Aldrich).

#### Tracking migrating breast cancer cells.

Migrating MCF10DCIS.com cells were tracked using either StarDist directly implemented within TrackMate (Figure 2a) or using Cellpose and then TrackMate (Supplementary figure S6b). To track MCF10DCIS.com cells labeled with sir-DNA using StarDist and TrackMate, a custom StarDist model was generated using the ZeroCostDL4Mic platform [6, 24]. This custom StarDist model was trained for 100 epochs on 72 paired image patches (image dimensions: 1024×1024, patch size: 1024×1024) with a batch size of 2 and a mae loss function, using the StarDist 2D ZeroCostDL4Mic notebook (v1.12.2). The StarDist "Versatile fluorescent nuclei" model was used as a training starting point. Key python packages used include TensorFlow (v1.15), Keras (v2.3.1), CSBdeep (v0.6.1), NumPy (v1.19.5) and Cuda (v10.1.243). The training was accelerated using a Tesla P100 GPU. This model generated excellent segmentation results on our test dataset (average Intersection over union > 0.96; average F1 score > 0.96). This model, the training dataset as well as the code used for training is available on Zenodo [28]. In TrackMate, the StarDist detector custom model (*score threshold* = 0.41 and *overlap threshold* = 0.5) and the LAP tracker (*linking max distance* = 30 µm; *track segment splitting* = 15 µm) were used. Tracks were filtered in function of their total distance traveled and tracks shorter than 80 µm were excluded.

To track MCF10DCIS.com cells expressing lifeact-RFP (cell line described here [29]) and labeled with sir-DNA, cells were first segmented using the ZeroCostDL4Mic Cellpose 2D notebook (v1.12, [24, 25]). The Cellpose model Cyto was used for the segmentation and the lifeact staining was used as the main segmentation channel. The Sir-DNA channel was used as the secondary segmentation channel. The following Cellpose parameters were used Flow threshold = 0.4 and Cell probability threshold = 0, Object diameter: 50. The quality of the segmentation was assessed visually. In TrackMate, the label image detector and the LAP tracker (*linking max distance* = 30 µm; *track segment gap closing* = 15 µm and 2 frames; *track segment splitting* = 15 µm) were used. Tracks were filtered in function of the total number of spots detected and tracks with less than 40 spots were excluded.

#### Tracking migrating T cells.

T cells migrating on ICAM-1 were automatically tracked using StarDist directly implemented within TrackMate (Figure 2b). In TrackMate, the StarDist detector custom model (Score threshold = 0.41 and Overlap threshold = 0.5) and the Simple LAP tracker (*linking max distance* = 30 µm; *gap closing max distance* = 15 µm, *gap closing max frame gap* = 2 frames) were used. The StarDist model used was described previously [30] and is publicly available on Zenodo [31].

#### Following ERK activity in migrating cells.

MBA-MD-231 or U2OS cells stably expressing clover-ERK-KTR were seeded on fibronectin-coated (1 µg /ml) Ibidi 8 well slides (Ibidi) one day before imaging. 4h before imaging, the media was supplemented with 250 nM sirDNA (Cytoskeleton Inc) and 25 mM HEPES (Sigma). Cells were then imaged live (37°C, 5% $CO_2$) using a Nikon Eclipse Ti2-E microscope (Nikon) equipped with a sCMOS Orca Flash4.0 camera (Hamamatsu) and controlled by the NIS-Elements software (Nikon, v 5.11.01). MBA-MD-231 cells were imaged using a 20x Nikon CFI Plan Apo Lambda objective (NA 0.75), either one frame per minute for 2 h or one frame every 5 minutes for 17 h. In these experiments, a camera binning of 2x2 was used. U2OS cells were imaged using a 10x Nikon CFI Plan-Fluor objective (NA 0.3) every 5 minutes for 3 hours. Cell nuclei were automatically tracked over time by using StarDist in TrackMate.

To track the nuclei of U2OS cells, a custom StarDist model was trained using the ZeroCostDL4Mic platform [24]. The training source for the model was generated from 25 manually annotated images (dimensions: 2048×2048) using the LOCI plugin in Fiji. The generated training source and target were randomly cropped into size 1024×1024, rotated, flipped and multiplied by five using the Augmentor ZeroCostDL4Mic notebook [24, 32] to generate a dataset of 120 paired images.

The custom StarDist model was trained for 200 epochs on the 120 paired image patches (image dimensions: 1024×1024, patch size: 1024×1024) with a batch size of 2 and a mae loss function, using the StarDist 2D ZeroCostDL4Mic notebook (v1.12.2) [24]). Key python packages used include TensorFlow (v1.15), Keras (v2.3.1), CSBdeep (v0.6.1), NumPy (v1.19.5) and Cuda (v11). The training was accelerated using a Tesla P100GPU. This model generated excellent segmentation results on our test dataset (average F1 score > 0.918).

In TrackMate, the StarDist detector custom model (Score threshold = 0.41 and Overlap threshold = 0.5) and the LAP tracker (linking max distance = 20 µm; Track segment gap closing = 25 µm, Gap closing max frame gap = 10 frames) were used. Tracks were filtered in function of their track duration and tracks shorter than 34 frames (2h 40min) were excluded.

To track the nuclei of MDA-MB-231 cells, the "Versatile fluorescent nuclei" StarDist model was used. In TrackMate, the StarDist detector (Score threshold = 0.41 and Overlap threshold = 0.5) and the LAP tracker (linking max distance = 40 µm; Track segment splitting = 30 µm) were used. Tracks were filtered in function of their duration and only the tracks spanning the whole movie were considered for further analysis (directly in TrackMate). For each tracked cell, the average intensity of the ERK reporter was measured in their nucleus over time (directly in TrackMate). To visualise the changes in ERK activity over time, results were uploaded to PlotTwist [33], data were rescaled between 0 and 1 and visualised as heatmaps.

#### Tracking Mouse hematopoietic stem cells migrating in hydrogel microwells.

Mouse hematopoietic stem cells migrating in a hydrogel microwell [34] were automatically segmented using Cellpose (Cyto model) implemented in the ZeroCostDL4Mic platform [24, 25]. The following Cellpose parameters were used: *flow threshold* = 0.4 and *cell probability threshold* = 0, *object diameter* = 17. The quality of the segmentation was assessed visually. The resulting label images were automatically tracked using TrackMate. In TrackMate, the label image detector and the LAP tracker (*linking max distance* = 30 µm; *track segment gap closing* = 15 µm and 2 frames; track segment splitting = 15 µm) were used. Spots were filtered in function of their circularity and area. Tracks were filtered in function of the total distance travelled tracks shorter than 80 µm were excluded. This dataset is available from the Cell Tracking Challenge website [4].

#### *Neisseria meningitidis* sample preparation and imaging.

The *Neisseria meningitidis* strain 2C43 [35] pilQ/pilQ-mCherry$_{ind}$ was grown on GCB agar plates (Difco) containing Kellog's supplements, 3 µg/ml vancomycin and 5 µg/ml chloramphenicol at 37°C in moist atmosphere containing 5% $CO_2$. The pMGC17 plasmid was designed in order to generate the 2C43 pilQ/pilQ-mCherry$_{ind}$ strain allowing IPTG-inducible expression of the type IV pilus secretin protein PilQ with a carboxy terminal fusion to mCherry expressed from the *Neisseria meningitidis* chromosome. First, pilQ was PCR-amplified from *Neisseria meningitidis* chromosomal DNA with primers pilQ-F:
TTAATTAAAGGAGTAATTTTATGAATACCAAACTGAC
AAAAATC
and pilQ-R: GTCGACTCAATAGCGCAGGCTGTTGC.
This PCR fragment was cloned in a pCRII-Blunt-TOPO vector (Invitrogen). Then, the mCherry ORF was PCR-amplified with a forward primer containing a region homologous to the 3' of pilQ (minus the stop codon) as well as a Gly-Ser-Gly linker, and a reverse primer containing a SalI restriction site and a region homologous to the TOPO vector
(MUTmChCT-F:
AGCCTGCGCTATGGTTCCGGTGTGAGCAAGGGC,
and MUTmChCT-R:
CTGCAGAATTCGCCCTTGTCGACTCACTTGTACAG).
This PCR fragment was used as a mutagenesis megaprimer to amplify pilQ from the TOPO vector [36]. Finally, this vector was digested with PacI and SalI restriction enzymes and the resulting insert ligated into pMGC10. The pMGC10 plasmid was generated by inserting the lacI gene and the lac promoter in the pMGC3 plasmid [37]. The fragment of interest was PCR amplified from the pMMB207 plasmid [38] using primers:
LacIF2: GAATTCGCTAACTTACATTAATTGCGTTGC
and LacIPR:
GTCGACGATCTTAATTAATTTCCTGTGTGAAATTGTTA
TCCG
and cloned in pMGC3 using EcoRI and SalI restriction. The pMGC17 plasmid was used to transform *Neisseria meningitidis*, generating an intermediate strain that carries both a native pilQ and pilQ-mCherry. This strain was then transformed with chromosomal DNA from a pilQ mutant strain [39, 40].

Bacteria in exponential phase from a 2 hours pre-culture in RPMI+10% FBS supplemented with 100 µM IPTG at 37°C and 5% $CO_2$ were diluted to an optical density of 0.015 ($\approx 1.5 \times 10^7$ bacteria/ml) and dropped onto a 2% agarose gel supplemented with 100 µM IPTG. Once the bacteria-containing droplet had dried up, the agar pad was flipped down onto a Fluorodish (Ibidi, 60 µm-Dish, 35 mm high Glass bottom). Fluorescently labeled proliferating bacteria were acquired using an inverted spinning-disk confocal microscope (Nikon, TI Eclipse) equipped with a 100X immersion objective (PlanFluor, NA = 0.5 – 1.3) at 37°C in the presence of 5% $CO_2$. Bacterial fluorescence was imaged in time-lapse at 5 min frame rate with an exposure time of 300 ms for 5.5 hours, and recorded with a CMOS Camera (Photometrics, 95BPrime) using Metamorph Imaging Software (Molecular Devices). The focus was maintained with the Perfect Focus System (PFS, Nikon).

#### Tracking focal adhesions in endothelial cells.

Live imaging of the endothelial cells expressing Paxillin-eGFP was described previously [41]. Briefly, human dermal microvascular blood endothelial cells expressing Paxillin were imaged using a Marianas spinning disk confocal microscope. This microscope was controlled by SlideBook 6 (Intelligent Imaging Innovations, Inc.), equipped with a Yokogawa CSU-W1 scanning unit, an inverted Zeiss Axio Observer Z1 body and a 100x, NA 1.4 oil (Plan-Apochromat, M27) objective. Images were acquired every two minutes using an Orca Flash4 sCMOS camera (chip size 2048×2048; 2×2 camera binning enabled; Hamamatsu Photonics), at 37°C and in the presence of 5% $CO_2$. Acquired images were then processed using Fiji to remove background (rolling ball radius: 10 pixels), compensate for bleaching (exponential fit method), and correct drifting (StackReg, Rigid body). A custom Weka pixel classifier was then trained in Fiji to segment the focal adhesions. In TrackMate, the Weka detector (*Threshold on probability* = 0.5) and the overlap tracker (*min IoU* = 0.3, *scale factor* = 1) were used.

#### Tracking 2D labels to generate 3D labels.

To form spheroids, MCF10 DCIS.com cells were seeded as single cells, in standard growth media, at low density (∼3,000 cells per well) on growth factor reduced (GFR) Matrigel-coated glass-bottom dishes (coverslip No. 0; MatTek). After 12 h, the medium was replaced by a normal growth medium supplemented with 2% (vol/vol) GFR Matrigel. After six days, spheroids were fixed with 4% PFA for 10 min at room temperature and labelled using Dapi. Spheroids were then imaged using a spinning-disk confocal microscope (Z step = 0.5 µm). The spinning-disk confocal microscope used was a Marianas spinning disk imaging system with a Yokogawa CSU-W1 scanning unit on an inverted Zeiss Axio Observer Z1 microscope (Intelligent Imaging Innovations, Inc.) equipped with a 100x (NA 1.4) oil, Plan-Apochromat, M27 (Zeiss). To generate 3D labels, nuclei were detected and tracked across the Z volume using StarDist implemented in TrackMate. In TrackMate, the StarDist detector (*score threshold* = 0.41 and *overlap threshold* = 0.5) and the LAP tracker (*linking max distance* = 1 µm, track merging and splitting enabled) were used. Detected spots were filtered in function of their mean intensity to exclude spots with weak intensities. Tracks were filtered in function of the number of spots per track and only the tracks with more than 3 spots were considered for further analysis (directly in TrackMate). In TrackMate, tracked nuclei were then exported as a label image to create 3D labels. 3D labels were then visualized using the FPBioimage software [42]. The video was generated using Arivis Vision4D (v 3.4).

Confocal images of Arabidopsis Thaliana floral meristem [21, 22] and light-sheet images of a developing Drosophila Melanogaster embryo [3, 4, 23] were automatically segmented using Cellpose (Cyto2 model) implemented in the ZeroCostDL4Mic platform [24, 25]. The following Cellpose parameters were used: *flow threshold* = 0.4 and *cell probability threshold* = 0, *object diameter* = 0. The quality of the segmentation was assessed visually. To generate 3D labels, the 2D label images were tracked using TrackMate. In TrackMate, the label image detector and the simple LAP tracker were used. The video were generated using Arivis Vision4D (v 3.4).

### References.

1. Nicolas Chenouard, Ihor Smal, Fabrice De Chaumont, Martin Maška, Ivo F. Sbalzarini, Yuanhao Gong, Janick Cardinale, Craig Carthel, Stefano Coraluppi, Mark Winter, Andrew R. Cohen, William J. Godinez, Karl Rohr, Yannis Kalaidzidis, Liang Liang, James Duncan, Hongying Shen, Yingke Xu, Klas E.G. Magnusson, Joakim Jaldén, Helen M. Blau, Perrine Paul-Gilloteaux, Philippe Roudot, Charles Kervrann, François Waharte, Jean Yves Tinevez, Spencer L. Shorte, Joost Willemse, Katherine Celler, Gilles P. Van Wezel, Han Wei Dan, Yuh Show Tsai, Carlos Ortiz De Solórzano, Jean Christophe Olivo-Marin, and Erik Meijering. Objective comparison of particle tracking methods. *Nature methods*, 11(3):281–289, 2014. ISSN 1548-7091. doi:10.1038/nmeth.2808.
2. Single-particle tracking challenge website. http://bioimageanalysis.org/track/, 2014. (Accessed on 06/13/2021).
3. Vladimír Ulman, Martin Maška, Klas E.G. Magnusson, Olaf Ronneberger, Carsten Haubold, Nathalie Harder, Pavel Matula, Petr Matula, David Svoboda, Miroslav Radojevic, Ihor Smal, Karl Rohr, Joakim Jaldén, Helen M. Blau, Oleh Dzyubachyk, Boudewijn Lelieveldt, Pengdong Xiao, Yuexiang Li, Siu Yeung Cho, Alexandre C. Dufour, Jean Christophe Olivo-Marin, Constantino C. Reyes-Aldasoro, Jose A. Solis-Lemus, Robert Bensch, Thomas Brox, Johannes Stegmaier, Ralf Mikut, Steffen Wolf, Fred A. Hamprecht, Tiago Esteves, Pedro Quelhas, Ömer Demirel, Lars Malmström, Florian Jug, Pavel Tomancak, Erik Meijering, Arrate Muñoz-Barrutia, Michal Kozubek, and Carlos Ortiz-De-Solorzano. An objective comparison of cell-tracking algorithms. *Nature Methods*, 2017. ISSN 15487105. doi:10.1038/nmeth.4473.
4. Cell tracking challenge website. http://celltrackingchallenge.net/, 2017. (Accessed on 06/13/2021).
5. D Sage, F R Neumann, F Hediger, S M Gasser, and M Unser. Automatic tracking of individual fluorescence particles: application to the study of chromosome dynamics. *IEEE transactions on image processing : a publication of the IEEE Signal Processing Society*, 14 (9):1372–1383, sep 2005. ISSN 1057-7149. doi:10.1109/TIP.2005.852787.
6. Uwe Schmidt, Martin Weigert, Coleman Broaddus, and Gene Myers. Cell Detection with Star-Convex Polygons. In *Lecture Notes in Computer Science (including subseries Lecture Notes in Artificial Intelligence and Lecture Notes in Bioinformatics)*, pages 265–273. Springer, jun 2018. ISBN 9783030009335. doi:10.1007/978-3-030-00934-2_30.
7. John C. Crocker and David G. Grier. Methods of digital video microscopy for colloidal studies. *Journal of Colloid and Interface Science*, 1996. ISSN 00219797. doi:10.1006/jcis.1996.0217.
8. Saga Helgadottir, Aykut Argun, and Giovanni Volpe. Digital video microscopy enhanced by deep learning. *Optica*, 6(4):506–513, Apr 2019. doi:10.1364/OPTICA.6.000506.
9. Benjamin Midtvedt, Saga Helgadottir, Aykut Argun, Jesús Pineda, Daniel Midtvedt, and Giovanni Volpe. Quantitative digital microscopy with deep learning. *Applied Physics Reviews*, 8(1):011310, mar 2021. ISSN 1931-9401. doi:10.1063/5.0034891.
10. Jay M. Newby, Alison M. Schaefer, Phoebe T. Lee, M. Gregory Forest, and Samuel K. Lai. Convolutional neural networks automate detection for tracking of submicron-scale particles in 2D and 3D. *Proceedings of the National Academy of Sciences of the United States of America*, 2018. ISSN 10916490. doi:10.1073/pnas.1804420115.
11. P. Zelger, K. Kaser, B. Rossboth, L. Velas, G. J. Schütz, and A. Jesacher. Three-dimensional localization microscopy using deep learning. *Optics Express*, 2018. ISSN 1094-4087. doi:10.1364/oe.26.033166.
12. Samira Masoudi, Afsaneh Razi, Cameron H.G. Wright, Jesse C. Gatlin, and Ulas Bagci. Instance-Level Microtubule Tracking. *IEEE Transactions on Medical Imaging*, 2020. ISSN 1558254X. doi:10.1109/TMI.2019.2963865.
13. T. Wollmann, C. Ritter, J. N. Dohrke, J. Y. Lee, R. Bartenschlager, and K. Rohr. Detnet: Deep neural network for particle detection in fluorescence microscopy images. In *Proceedings - International Symposium on Biomedical Imaging*, 2019. ISBN 9781538636411. doi:10.1109/ISBI.2019.8759234.
14. Simon Franchini and Samuel Krevor. Cut, overlap and locate: a deep learning approach for the 3D localization of particles in astigmatic optical setups. *Experiments in Fluids*, 2020. ISSN 14321114. doi:10.1007/s00348-020-02968-w.
15. Fabrice De Chaumont. ISBI Single Particle Tracking Challenge benchmark generator - Icy plugin. http://icy.bioimageanalysis.org/plugin/isbi-challenge-tracking-benchmark-generator/, Dec 2013. (Accessed on 06/13/2021).
16. Fabrice De Chaumont, Stéphane Dallongeville, Nicolas Chenouard, Nicolas Hervé, Sorin Pop, Thomas Provoost, Vannary Meas-Yedid, Praveen Pankajakshan, Timothée Lecomte, Yoann Le Montagner, Thibault Lagache, Alexandre Dufour, and Jean Christophe Olivo-Marin. Icy: An open bioimage informatics platform for extended reproducible research. *Nature Methods*, 2012. ISSN 15487091. doi:10.1038/nmeth.2075.
17. Dmitry Ershov. TrackMate Stardist ISBI SPT challenge training . https://gitlab.pasteur.fr/iah-public/trackmate-stardist-isbi-spt-challenge-training, Jun 2021. (Accessed on 08/20/2021).
18. Jean Yves Tinevez, Nick Perry, Johannes Schindelin, Genevieve M. Hoopes, Gregory D. Reynolds, Emmanuel Laplantine, Sebastian Y. Bednarek, Spencer L. Shorte, and Kevin W. Eliceiri. TrackMate: An open and extensible platform for single-particle tracking. *Methods*, 115:80–90, 2017. ISSN 10959130. doi:10.1016/j.ymeth.2016.09.016.
19. Jean-Yves Tinevez. ISBI SPT Challenge performance metrics batch code. https://gitlab.pasteur.fr/iah-public/TrackingPerformance, Jun 2021. (Accessed on 06/14/2021).
20. Jean-Yves Tinevez. Benchmarking TrackMate code. https://github.com/fiji/TrackMate/tree/paper/src/test/java/fiji/plugin/trackmate/paper, Jun 2021. (Accessed on 06/14/2021).
21. Anuradha Kar. Arabidopsis dataset, original stacks and segmented data. https://figshare.com/articles/dataset/Original_stacks_and_segmented_data/14447079/1, 2021.
22. Anuradha Kar, Manuel Petit, Yassin Refahi, Guillaume Cerutti, Christophe Godin, and Jan Traas. Assessment of deep learning algorithms for 3d instance segmentation of confocal image datasets. *bioRxiv*, 2021. doi:10.1101/2021.06.09.447748.
23. Martin Maška, Vladimír Ulman, David Svoboda, Pavel Matula, Petr Matula, Cristina Ederra, Ainhoa Urbiola, Tomás España, Subramanian Venkatesan, Deepak M.W. Balak, Pavel Karas, Tereza Bolcková, Markéta Štreitová, Craig Carthel, Stefano Coraluppi, Nathalie Harder, Karl Rohr, Klas E. G. Magnusson, Joakim Jaldén, Helen M. Blau, Oleh Dzyubachyk, Pavel Křížek, Guy M. Hagen, David Pastor-Escuredo, Daniel Jimenez-Carretero, Maria J. Ledesma-Carbayo, Arrate Muñoz-Barrutia, Erik Meijering, Michal Kozubek, and Carlos Ortiz-de Solorzano. A benchmark for comparison of cell tracking algorithms. *Bioinformatics*, 30(11):1609–1617, 02 2014. ISSN 1367-4803. doi:10.1093/bioinformatics/btu080.
24. Lucas von Chamier, Romain F. Laine, Johanna Jukkala, Christoph Spahn, Daniel Krentzel, Elias Nehme, Martina Lerche, Sara Hernández-Pérez, Pieta K. Mattila, Eleni Karinou, Séamus Holden, Ahmet Can Solak, Alexander Krull, Tim-Oliver Buchholz, Martin L. Jones, Loïc A. Royer, Christophe Leterrier, Yoav Shechtman, Florian Jug, Mike Heilemann, Guillaume Jacquemet, and Ricardo Henriques. Democratising deep learning for microscopy with ZeroCostDL4Mic. *Nature Communications*, 12(1):2276, dec 2021. ISSN 2041-1723. doi:10.1038/s41467-021-22518-0.
25. Carsen Stringer, Tim Wang, Michalis Michaelos, and Marius Pachitariu. Cellpose: a generalist algorithm for cellular segmentation. *Nature Methods*, 18(1):100–106, December 2020. doi:10.1038/s41592-020-01018-x.
26. Sergi Regot, Jacob J. Hughey, Bryce T. Bajar, Silvia Carrasco, and Markus W. Covert. High-sensitivity measurements of multiple kinase activities in live single cells. *Cell*, 2014. ISSN 10974172. doi:10.1016/j.cell.2014.04.039.
27. Takamasa Kudo, Stevan Jeknić, Derek N. Macklin, Sajia Akhter, Jacob J. Hughey, Sergi Regot, and Markus W. Covert. Live-cell measurements of kinase activity in single cells using translocation reporters. *Nature Protocols*, 2018. ISSN 17502799. doi:10.1038/nprot.2017.128.
28. Guillaume Jacquemet. Combining StarDist and TrackMate example 1 - Breast cancer cell dataset. https://doi.org/10.5281/zenodo.4034976, September 2020.
29. Guillaume Jacquemet, Ilkka Paatero, Alexandre F. Carisey, Artur Padzik, Jordan S. Orange, Hellyeh Hamidi, and Johanna Ivaska. FiloQuant reveals increased filopodia density during breast cancer progression. *Journal of Cell Biology*, 216(10):3387–3403, August 2017. doi:10.1083/jcb.201704045.
30. Elnaz Fazeli, Nathan H. Roy, Gautier Follain, Romain F. Laine, Lucas von Chamier, Pekka E. Hänninen, John E. Eriksson, Jean-Yves Tinevez, and Guillaume Jacquemet. Automated cell tracking using StarDist and TrackMate. *F1000Research*, 9:1279, December 2020. doi:10.12688/f1000research.27019.2.
31. Nathan H. Roy and Guillaume Jacquemet. Combining StarDist and TrackMate example 2 - T cell dataset. https://10.5281/zenodo.4034929, 2020.
32. Marcus D Bloice, Peter M Roth, and Andreas Holzinger. Biomedical image augmentation using Augmentor. *Bioinformatics*, 35(21):4522–4524, 04 2019. ISSN 1367-4803. doi:10.1093/bioinformatics/btz259.
33. Joachim Goedhart. PlotTwist: A web app for plotting and annotating continuous data. *PLOS Biology*, 18(1):e3000581, January 2020. doi:10.1371/journal.pbio.3000581.
34. Matthias P. Lutolf, Regis Doyonnas, Karen Havenstrite, Kassie Koleckar, and Helen M. Blau. Perturbation of single hematopoietic stem cell fates in artificial niches. *Integr. Biol.*, 1(1):59–69, 2009. doi:10.1039/b815718a.
35. Xavier Nassif, Jonathan Lowy, Paula Stenberg, Peadar O'Gaora, Amir Ganji, and Magdalene So. Antigenic variation of pilin regulates adhesion of neisseria meningitidis to human epithelial cells. *Molecular Microbiology*, 8(4):719–725, 1993. doi:10.1111/j.1365-2958.1993.tb01615.x.
36. Song-Hua Ke and Edwin L. Madison. Rapid and efficient site-directed mutagenesis by single-tube 'megaprimer' PCR method. *Nucleic Acids Research*, 25(16):3371–3372, 08 1997. ISSN 0305-1048. doi:10.1093/nar/25.16.3371.
37. Magali Soyer, Arthur Charles-Orszag, Thibault Lagache, Silke Machata, Anne-Flore Imhaus, Audrey Dumont, Corinne Millien, Jean-Christophe Olivo-Marin, and Guillaume Duménil. Early sequence of events triggered by the interaction of neisseria meningitidis with endothelial cells. *Cellular Microbiology*, 16(6):878–895, 2014. doi:10.1111/cmi.12248.
38. Victor M. Morales, Assar Bäckman, and Michael Bagdasarian. A series of wide-host-range low-copy-number vectors that allow direct screening for recombinants. *Gene*, 97(1):39–47, 1991. ISSN 0378-1119. doi:10.1016/0378-1119(91)90007-X.
39. Marie-Claude Geoffroy, Stéphanie Floquet, Arnaud Métais, Xavier Nassif, and Vladimir Pelicic. Large-scale analysis of the meningococcus genome by gene disruption: Resistance to complement-mediated lysis. *Genome Research*, 13(3):391–398, 2003. doi:10.1101/gr.664303.
40. Michaella Georgiadou, Marta Castagnini, Gouzel Karimova, Daniel Ladant, and Vladimir Pelicic. Large-scale study of the interactions between proteins involved in type iv pilus biology in neisseria meningitidis: characterization of a subcomplex involved in pilus assembly. *Molecular Microbiology*, 84(5):857–873, 2012. doi:10.1111/j.1365-2958.2012.08062.x.
41. Laura Hakanpaa, Elina A. Kiss, Guillaume Jacquemet, Ilkka Miinalainen, Martina Lerche, Camilo Guzmán, Eero Mervaala, Lauri Eklund, Johanna Ivaska, and Pipsa Saharinen. Targeting $\beta$1-integrin inhibits vascular leakage in endotoxemia. *Proceedings of the National Academy of Sciences*, 115(28):E6467–E6476, June 2018. doi:10.1073/pnas.1722317115.
42. Marcus Fantham and Clemens F. Kaminski. A new online tool for visualization of volumetric data. *Nature Photonics*, 11(2):69–69, feb 2017. ISSN 1749-4885. doi:10.1038/nphoton.2016.273.
