## Supplemental manual for "Bringing TrackMate into the era of machine-learning and deep-learning"

### TrackMate version 7 documentation and tutorials.

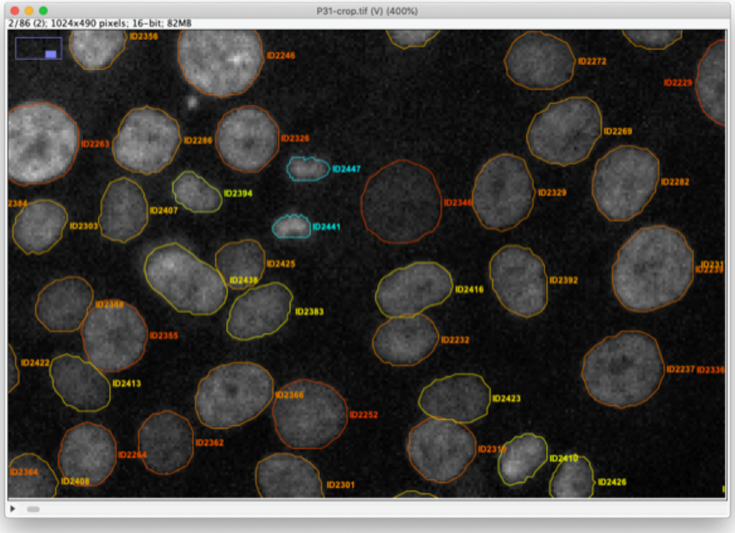

Dmitry Ershov, Minh-Son Phan, Joanna W. Pylvänäinen, Stéphane U. Rigaud, Guillaume Jacquemet, and Jean-Yves Tinevez.

*August 24, 2021*

### Introduction.

In summer 2021 we released a new major version of TrackMate, starting the version 7 series. This version introduced numerous major changes both on the user side and the developer side. These changes were prompted by several goals:

- Integrate key machine learning, deep-learning and segmentation algorithms in TrackMate. Novel algorithms that were developed in the last decade proved to be very good against difficult segmentation challenges, such as the low signal-to-noise ratio we are used to in microscopy imaging, and high density of objects that prevents their proper individualization. The defects brought by these two challenges have very negative consequences on tracking accuracy. By integrating these algorithms in TrackMate we hoped to easily offer better tracking results in difficult cases in a convenient manner.

- Because TrackMate aims at being a convenient end-user tool, we aimed for integrating some algorithms natively in TrackMate. That is, they are called directly from TrackMate and do not require intervention or manipulation from the user.

- These segmentation algorithms return the shape of objects they detected. We therefore aimed at adding object contour analysis into TrackMate. Before version 7, objects in TrackMate where shapeless spots, containing only a position and radius. Now in 2D, objects can have a contour described by a simple polygon.

- And because we now have the shape objects, we aimed at adding morphological analysis in TrackMate as well.

- TrackMate is also a platform for tracking. It aims at being used as a library or extended to quickly develop new analysis pipelines, reusing the existing coding facility. So, we added a new API that allows analysts and developers to quickly and simply integrate their own segmentation algorithm in TrackMate.

- Finally, we took advantage of this overhaul to rewrite almost all the components in TrackMate and improve their functionality and decrease their complexity.

The full changelog for the new version is here:

https://github.com/fiji/TrackMate/releases/tag/TrackMate-7.0.4

All these changes prompted for additional user and developer documentation. This document *extends* the existing TrackMate documentation, that you can find here:

https://imagej.net/plugins/trackmate/#documentation-and-tutorials

The tutorials that follow assume you are already familiar with the previous version of TrackMate, as a user or a developer.

### The new TrackMate v7 detectors and support for object shape analysis.

Starting with version 7.0.0, TrackMate offers the possibility to segment objects, and store, display and quantify their shape. We used this new API to build simple detectors that can produce objects from a *label image*, a *mask* or a *grayscale image with a threshold*. But we also looked to integrate the state-of-the-art segmentation algorithms shipped with Fiji that do so. So, we integrated the *ilastik*, *MorphoLibJ*, *StarDist* and *Weka* plugins as detectors in TrackMate. This part lists the seven detectors that have been introduced by this version and documents the shape analysis features.

#### Detectors with segmentation capabilities.

##### Mask detector.

This detector creates objects from a black and white channel in the source image. You can add the mask as an extra channel in the source image. The objects will be built based on all the pixels have a value strictly larger than 0, which solves the issue of having a mask on 8-bit, 16-bit or 32-bit images. This detector is part of the core of TrackMate.

##### Thresholding detector.

The thresholding detector creates objects from a grayscale image (it can be one channel in a multi-channel image). You must specify a threshold value to segment the objects. This detector is also part of the core of TrackMate.

##### Label image detector.

Label images are especially convenient as an output of segmentation algorithms. Indeed, in some cases you might have different objects that are so close that they touch each other. If a segmentation algorithm can detect them, but outputs a black and white mask, they will appear as one object in the mask if they share a border.

In a label image, each object is represented by different integer values. For instance, the object #1 in a label image will be made from all the pixels that have a pixel value of 1, over a black background of 0. Object #2 will have the pixel value 2, etc. This allows resolving them even if they touch each other. This detector is also part of the core of TrackMate.

##### TrackMate-Ilastik.

This detector is not part of the core Fiji distribution. You need to subscribe to *two* update sites (The `ilastik` update site and the `TrackMate-Ilastik` update site) and to install ilastik to get it. The detector installation procedure and its documentation are given in a following chapter.

##### TrackMate-MorphoLibJ.

This detector is also not part of the core Fiji distribution. You need to subscribe to *two* update sites (The `IJPB-plugins` update site and the `TrackMate-MorphoLibJ` update site) to get it. The detector installation procedure and its documentation are given in a following chapter.

##### TrackMate-StarDist.

This detector is also not part of the core Fiji distribution. You need to have StarDist installed and running in your Fiji installation. This involves subscribing to the `CSBDeep` update site and the `StarDist` update site. And to the `TrackMate-StarDist` update site. The detector installation procedure and its documentation are given in a following chapter.

##### TrackMate-Weka.

This detector is also not part of the core Fiji distribution. But since the Weka Trainable Segmentation plugin is included in the core of Fiji, we just must subscribe to the `TrackMate-Weka` update site. The detector installation procedure and its documentation are given in a following chapter.

#### Limitations.

The detection of object shape in TrackMate two some limitations now that we repeat here.

1. **Object contours are only detected for 2D images**.

Source images can be 2D + T with multiple channels, but shapes won't be detected, displayed nor analyzed in 3D. This boils down to the fact that there are no (not yet) easy and robust way to handle 3D segmentation results in Fiji. This might change in the future with the work of others, but at least with version 7 we detect contours only for 2D images.

In the meantime, when presented with a 3D image, the detectors described above either return an error message (the StarDist detector) or create a spherical spot of radius computed so that the spherical spot has the same volume that of the 3D object returned by the specific segmentation algorithm.

2. **Object contours must be simple polygons.**

Simple polygons are polygon made of one closed segmented line that does not have any self-intersection. TrackMate does not handle holes in objects, not objects made of several disconnected components. This is a limitation that allows handling computing morphological features without ambiguity.

#### Simplifying contours.

Several of the new detectors have a configuration setting that allows to simplify contours. It is an important parameter that we describe here.

Object contours are polygon that wraps around individual objects, initially following individual pixels. For instance, the initial output of the threshold detector for one spot looks like this:

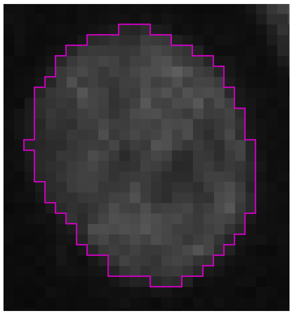

Notice that the polygon follows exactly the contour of all pixels that are above the threshold. For instance the leftmost pixel on the image above is has 3 segments for its border. And all the contour segments run along pixels horizontally or vertically.

Simplifying contour will yield a simplified shape of the object, that interpolate between pixels and return a smoother shape with fewer segments. The same algorithm running with the `Simplify contours` parameter selected will yield the following:

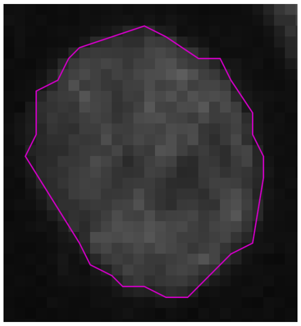

Simplifying contour generate TrackMate files that are smaller in disk space. More importantly, they yield more accurate morphological features. Indeed, the pixel-accurate contour overestimates the perimeter, because it sticks to individual pixel borders. In turn this generate contours that have an overestimated perimeter and will negatively affect the relevance of morphological feature that depend on it.

Simplifying contours somewhat tries to follow the object contour as if it would not be discretized over a pixel matrix. But it works well only if the objects are large enough. For small objects, below typically 10 pixels, the simplification generates inaccurate contours:

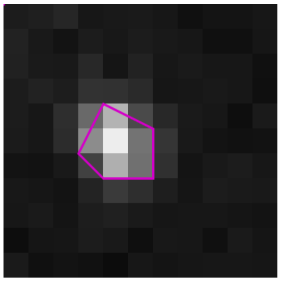

So as a rule of thumb we recommend the following:

- If your objects are big (N pixels larger than 10) and you want to measure their shape, always select the `Simplify contours` option.
- If your objects are sampled on a small number of pixels, it does not make sense to measure their morphology anyway.
- So basically, unselect the `Simplify contours` only if you want to later generate a pixel-accurate mask from the objects (with for instance the `Export label image` action).

#### Object morphology analysis.

One of the main goals of generating the object shape is to measure their shape. Detectors that return object contours trigger automatically the computation of morphological features in TrackMate. When relevant, these features use the physical calibration and units of the source image. This means for instance that if your image is calibrated with a pixel size in µm, the area of objects will be expressed in µm². These morphological features are:

##### Area.

Simply the area of the objects in spatial unit squared.

##### Perimeter.

The length of the contour in spatial unit.

##### Circularity.

The circularity is a measure of how close to a circle the shape of an object is. It has a value of 1 for circles and is getting close to 0 for very elongated objects. It is computed for 2D objects as

$$\frac{4 \times \pi \times \text{area}}{\text{perimeter}^2}$$

##### Solidity.

The solidity ranges from 0 to 1 and reports how smooth and convex is the object contour. A object with a dented contour, with many cavities will have a low solidity, close to 0. A perfectly convex object will have a value of 1. To compute it we first determine the convex hull of the object.

Intuitively, this is the contour we would get if we would wrap a rubber band around the object. It would stretch around the object contour and would not extend inside the cavities of the object. The area of this convex object is therefore always larger than the area of the initial object. Then the solidity is computed as:

$$\text{solidity} = \frac{\text{area}}{\text{convex area}}$$

#### Ellipse fit.

Several 2D morphological features are best obtained by fitting on ellipse on the object contour and reporting the ellipse parameters. In TrackMate, we first fit an ellipse to the contour using a direct fit following the Chernov method, computed using the Moore-Penrose pseudo inverse (by Kim van der Linde) for speed and robustness. TrackMate then reports the resulting ellipse parameters:

##### Ellipse center x0 and Ellipse center y0.

This the ellipse center position, with respect to the object center position. You can get the object center position by using

```
double x = spot.getDoublePosition( 0 );
double y = spot.getDoublePosition( 1 );
```

to which you need to add the ellipse center values to have the absolute position of the ellipse center.

##### Ellipse long axis and Ellipse short axis.

The length of the long and short axis of the ellipse, in physical units.

##### Ellipse angle.

The angle of the ellipse long axis with the X-axis of the image, in radians. Careful, in images the Y axis runs from top to bottom, so the positive angle direction is inverted compared to classical plots (positive angles are counterclockwise) on the image.

##### Ellipse aspect ratio.

The ellipse aspect ratio is the ratio of the major axis to the minor axis:

$$\text{ellipse AR} = \frac{\text{major axis}}{\text{minor axis}}$$

It ranges from 1 for ellipses that resembles circles and gets larger for elongated ellipses. A perfect line as a positive infinite aspect ratio.

### TrackMate Mask detector

This page describes a detector for TrackMate that creates objects from a black and white channel in the source image. You can add the mask as an extra channel in the source image. The objects will be built based on all the pixels have a value strictly larger than 0, which solves the issue of having a mask on 8-bit, 16-bit or 32-bit images.

#### Usage.

There is not much to say. You need to prepare a mask for the source image you want to track using any means that work.

For 2D images, TrackMate will obtain the contour of objects, possibly simplified, from the mask. For 3D images, it will create spherical spots of an identical volume.

It is a good idea to merge this mask as an extra channel of the source image, so that you can have both the mask and the source image in TrackMate. This allows measuring numerical features on the source image based on the contours obtained with the mask.

#### Tutorial: *C.elegans* early development.

##### The dataset.

We will use a simple and short image of a *C.elegans* embryo. You can download the data on Zenodo:

DOI 10.5281/zenodo.5132918

This movie is a maximum-intensity projection (MIP) of a longer movie used initially in:

> Tinevez JY, Dragavon J, Baba-Aissa L, Roux P, Perret E, Canivet A, Galy V, Shorte S. *A quantitative method for measuring phototoxicity of a live cell imaging microscope*. Methods Enzymol. 2012;506:291-309. doi: 10.1016/B978-0-12-391856-7.00039-1. PMID: 22341230.

This is very short movie, that stops after the 2nd cell division. We made a MIP so that this movie can be used in this tutorial.

##### The image.

Open the image `CelegansEarly_MIP.tif` in Fiji.

It is a 2 channels image. The first channel contains the raw image data. It shows the fluorescence collected from a *C.elegans* embryo, stained with eGFP-H2B, imaged on laser-scanning confocal microscope. We see the nuclei move and divide, along the with 2 smaller polar bodies. The 2nd channel contains the segmentation results of this signal, that I obtained by applying:

1. A median filter.
2. A Gaussian filter.
3. Then plain thresholding.

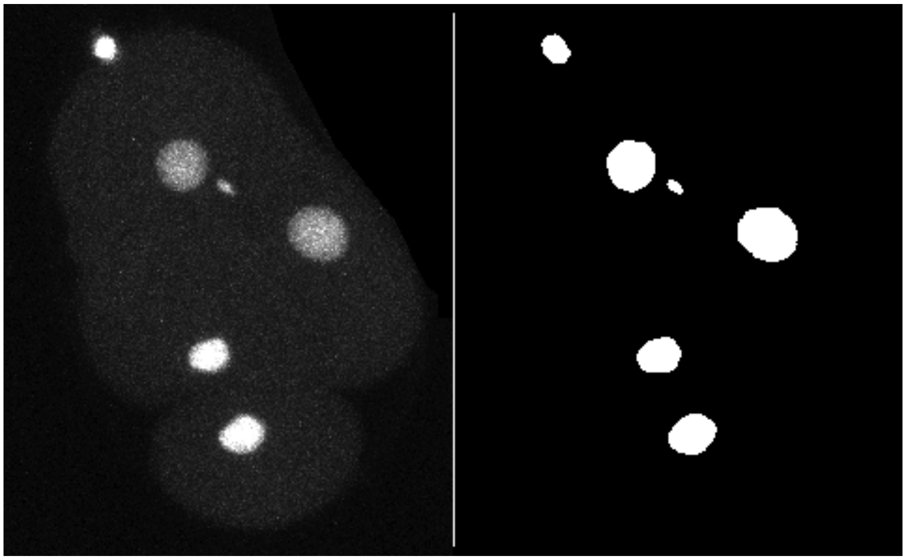

#### Running TrackMate.

Launch TrackMate: *Plugins > Tracking > TrackMate*

In the second panel, select the `Mask detector`

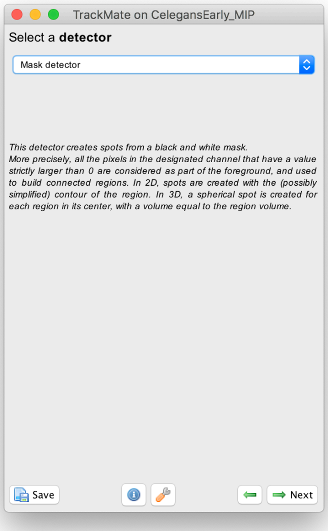

The configuration panel is very simple:

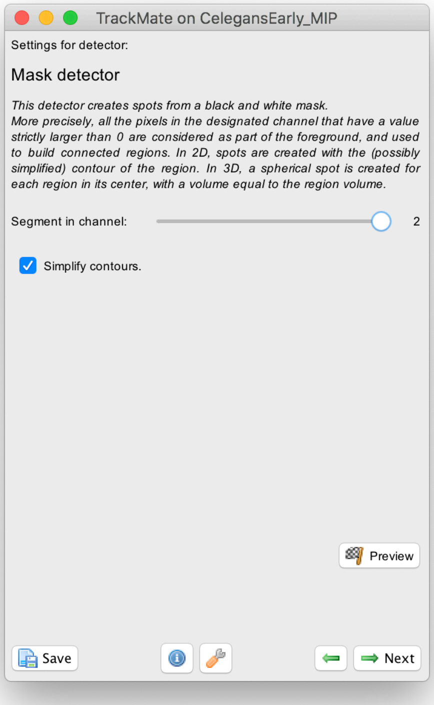

You just need to specify in what channel is the mask. In our case it is the second one. There is a checkbox that lets you simplify contours. We suggest you leave it on by default. The details of contour simplification are explained here. The `Preview` button will show the results. In frame 11 for instance, 6 objects are detected with their contour:

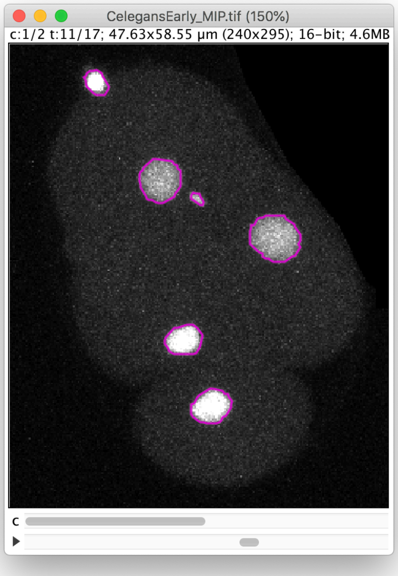

The click the `Next` button. In total 78 spots are found in this movie. You can track them with `LAP tracker`, enabling the detection of split events so that we can deal with the 2 cell divisions. In the end of tracking process, we get this:

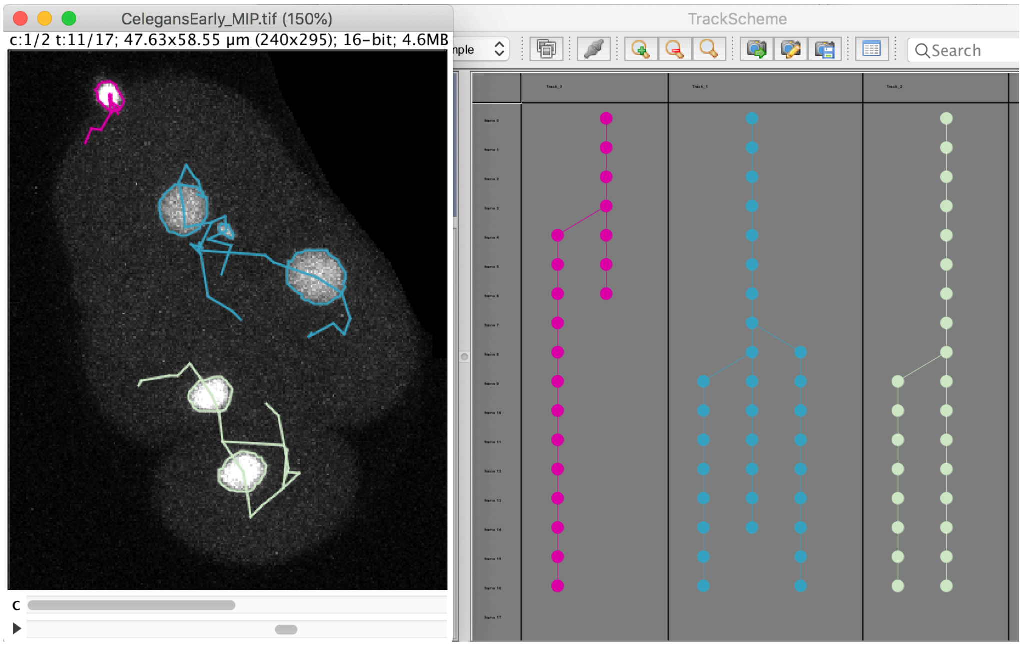

Notice that we have false divisions due to the motile polar body, that gets wrongly attached to the top cell movie. We could correct this either manually, or by filtering out the polar bodies based on their `Area`. We leave this correction as an exercise to the reader.

#### Plotting nucleus shape over time.

Now that we have the cell shape and their lineage, we can follow how the shape changes as the cell divides. For instance, let's plot the nucleus size and circularity of the bottom cell over time.

To do this, select the bottom cell by clicking in one of the spots, then moved to the `Plot feature` panel of the TrackMate wizard. Make sure you are in the `Spots` tab.

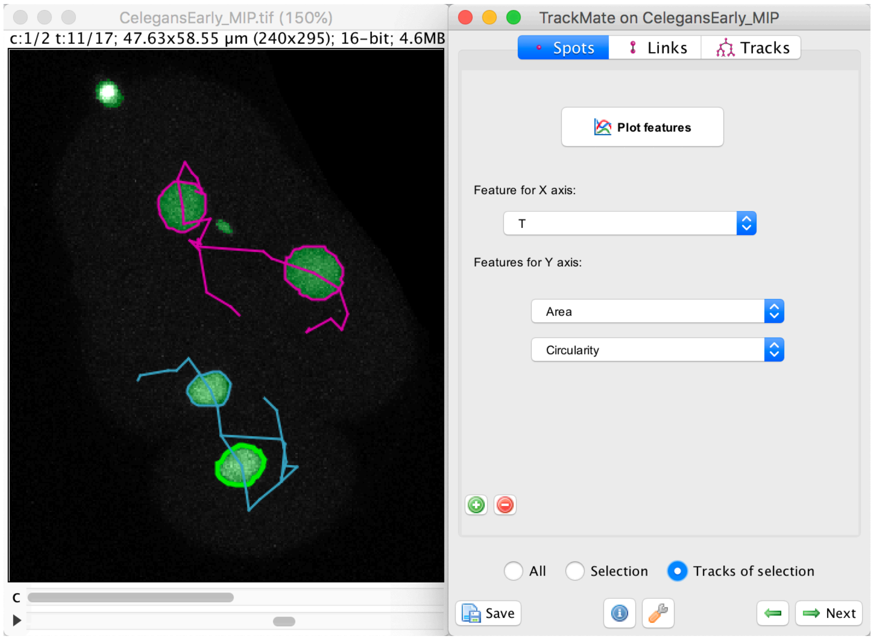

In this panel, select the `Area` in the `Feature for Y axis` first list.

Click on the green + button to add a second list of features and select the `Circularity` feature. We want to plot these feature values not for a single spot, nor for the whole dataset, but just for the track of the spot we selected. To do this, select the `Tracks of selection` radio button at the bottom. Of course, at least one spot in the track of interest must be selected.

Click on the `Plot features` button. The two graphs appear:

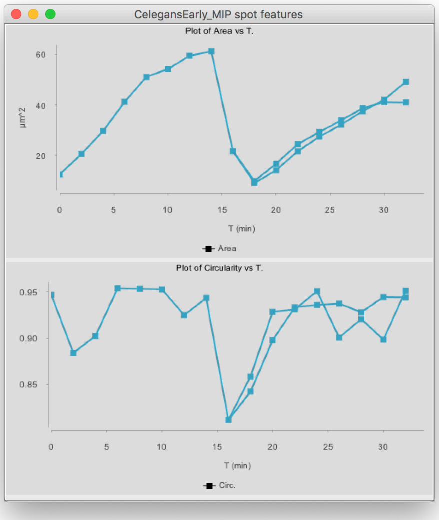

(You might have to deselect the cell by clicking elsewhere, then clicking back in the graph so that it shows the same color than above.)

The top graph shows the area over time. We see that it steadily increases until the cell divides at t=15 minutes. The area sharply decreases, and two cells are now plotted in the graph. Their area resume increasing, with almost identical rate.

The circularity is plotted on the second graph. It remains high almost all the time (close to 1), as the nuclei have a roughly spherical shape. When the cell divides, the nucleus takes an elongated shape, which results in a lower circularity.

As a side note, if you right-click on any of the 2 plots, you will have this popup menu that will let you export the graph as a pdf, png or svg image, or show its data in a table that can saved to a CSV file.

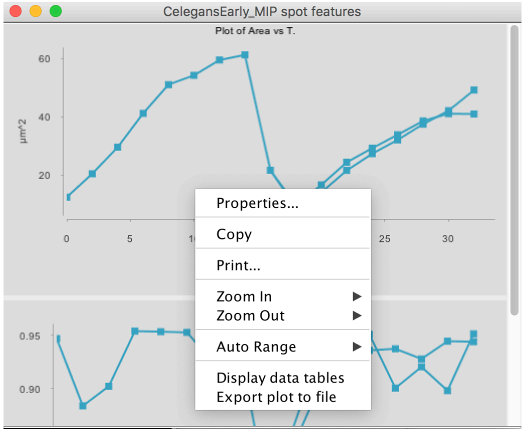

#### Editing the mask.

If we repeat the above procedure for the top cell, we have the following graphs:

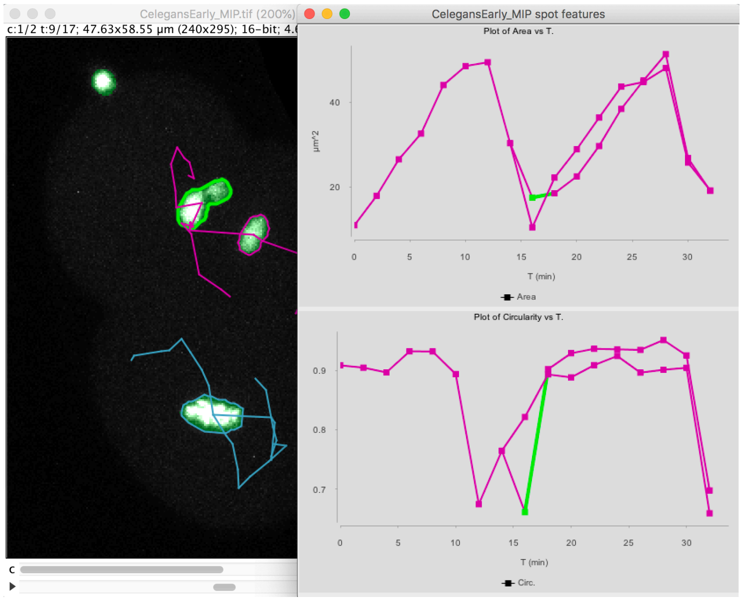

Note that at t=16 minutes, one of the daughter nuclei has a circularity that shows a sharp drop. By inspecting the image, we can see that this is caused by the object contour to be incorrect for this spot. The motile polar body went too close to the nucleus and the mask merged them together. So, we end up in having an artificially elongated object, which results in a low circularity.

Right now, there is no way to edit a spot contour. Here we would suggest a hack. We will directly edit the binary mask by cutting the object with Fiji tool.

Go back to the first panel of the TrackMate wizard and in the image display, select the 9th time-point, and make the second channel active:

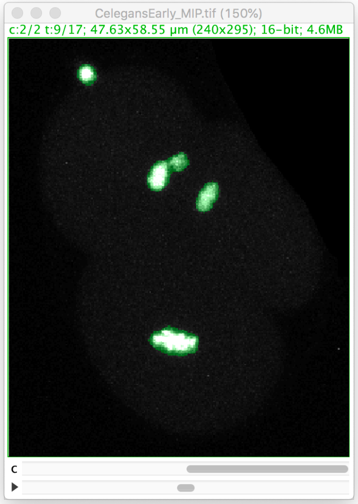

Let's zoom on the nucleus with the defective mask.

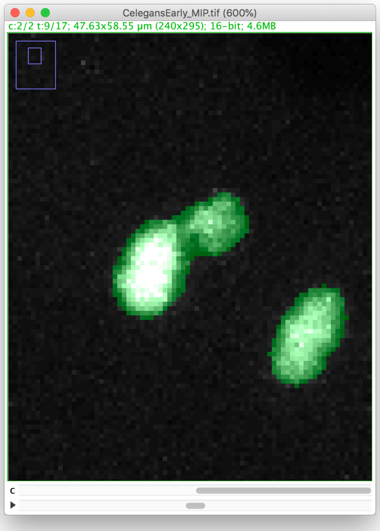

We will make a cut in the mask to separate the nucleus from the polar body. We can use the Image ROI to do so. In the ImageJ toolbar, select the line ROI tool. Double click on the color selection tool and select the black color as the foreground color. Now draw a line that goes between the nucleus and the polar body.

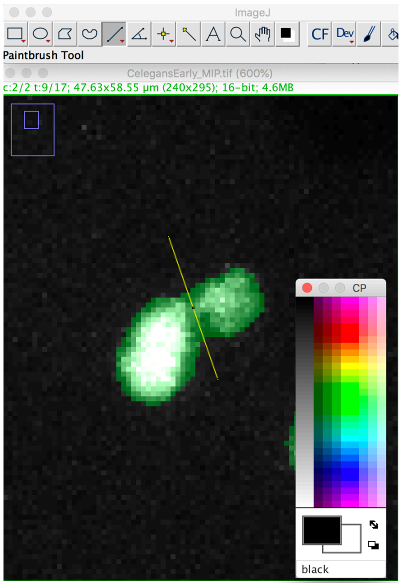

Press the `D` key to draw. Since we picked the black color, the pixels along the line ROI will be replaced by a pixel value of 0, cutting the mask into 2 components:

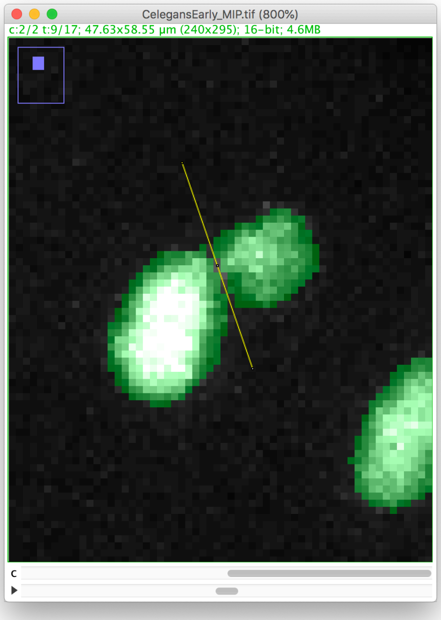

Now you can proceed with the tracking as before. The polar body and the nucleus will be segmented as two spots:

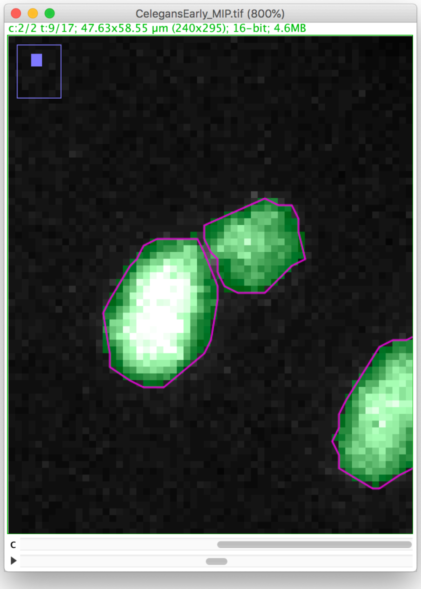

Right now, this is the only way to edit the spot contour in TrackMate. Which means it is important to get a good segmentation *result* before running TrackMate.

#### Tutorial: Cancer cells migration.

Get the tutorial dataset from Zenodo:

DOI 10.5281/zenodo.5243127

- Open Fiji.
- Open your image.
- Open TrackMate *Plugins > Tracking > TrackMate*. The start panel will launch, showing information about the image dimensions. Click `Next`.
- The “Select a detector” panel opens. From the pull-down menu, select the `Mask detector`. Click `Next`.

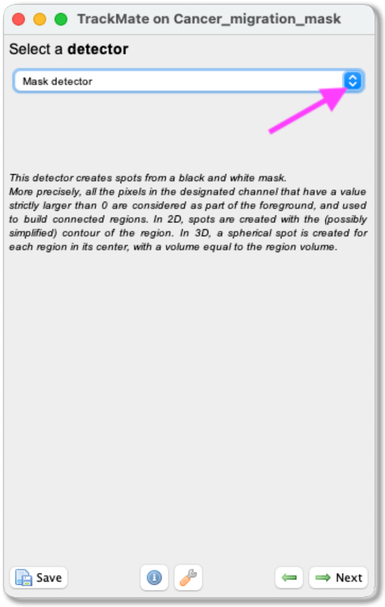

- A panel with the description of the mask detector opens. Here you can select to Simplify the contours to smooth the edges of the segmented objects. Click on `Preview` to see a preview of the segmentation. On the left, there will also be the number of detected objects in that field of view. Click `Next`. The objects are now being detected.
- When the progress bar has reached the end, click `Next`.
- A panel to filter the detected spots according to their quality opens (more information about this filtering can be found here. With our test image, this part can be ignored. Click `Next`.
- A panel to filter spots according to their properties (i.e., size, shape, location, or signal intensity) opens. With our test image, we do not need to filter any spots. Click `Next`.

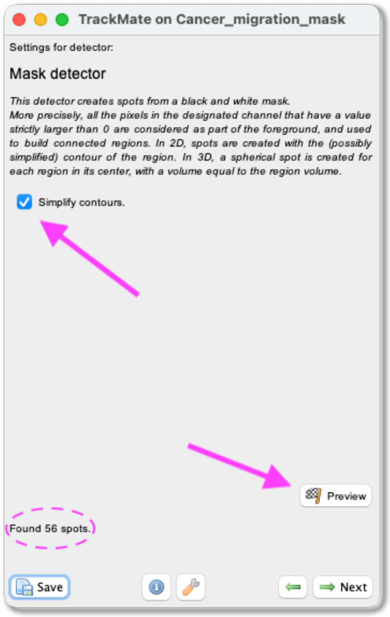

- A tracking panel opens. In this panel, you can select a method for tracking your objects. In this exercise, we use the `LAP tracker`. Please select it from the pull-down menu and click `Next`.

- A panel to choose the LAP tracker settings opens. First, with the `Frame to Frame linking` parameter, you give the maximum distance to link two objects between frames. With our test image, we use **20 microns**. Tick the `Allow gap closing` box and add values: `Max distance`: **20 microns** and `Max frame gap`: **5**. Next, you let TrackMate know if the tracks are allowed to split. Track splitting can occur, for example, due to cell division. Tick the box `Allow track segment splitting` and insert value `Max distance`: **20 microns**. Below you will also see a setting to allow Track segment merging. This box should remain unticked as we do not expect the nuclei to merge here. Click `Next`.

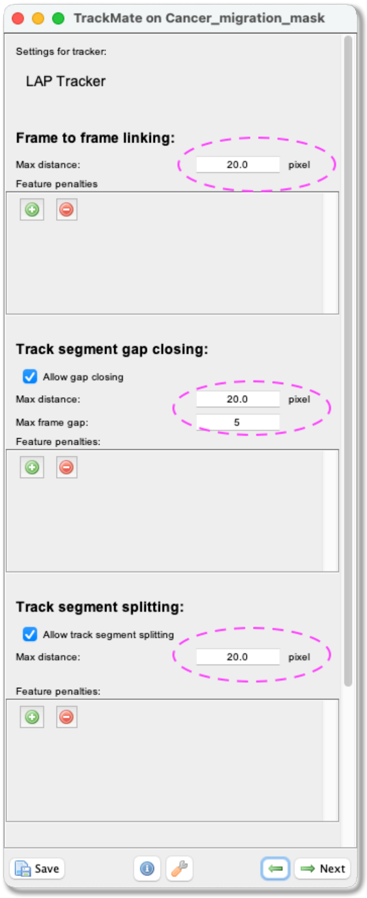

- A Track filter panel opens. In this panel, you can remove tracks according to their properties (*i.e.*, length, speed, or location). With our test image, we do not need to filter any tracks. Click `Next`.

- A window with track visualization options opens. Here it is possible to edit track or object colors according to their properties. We will label the tracked objects according to their mean speed and the tracks according to the total distance travelled.

  – First, make sure that the `Display spots` and `as ROIs` boxes are ticked.

- Select `Track mean speed` from the Color spots by clicking on the pull-down menu and then click `auto`.
- To fill the objects with the indicative color, click on `Edit settings`. This will open a new panel with display settings. Here, tick the box `draw spots filled`.
- From the same panel, also set the `line thickness` as **2**.
- Close the Display settings window.
- Make sure the `Display tracks` box is ticked.
- From the pulldown menu, select `Show tracks forward in time`. This will show the future trajectories of the object movements.
- From the Color tracks pull-down menu, select `Total distance travelled`. And click `auto`.

- In this panel, you can also export the results as .csv files. Please do so by clicking on **Tracks** at the bottom of the panel. A window with a lot of data will open. Ensure you export the **Spots** (information about the spot) and the **Tracks** (information about the tracks) files. Close the results window and click **Next**.

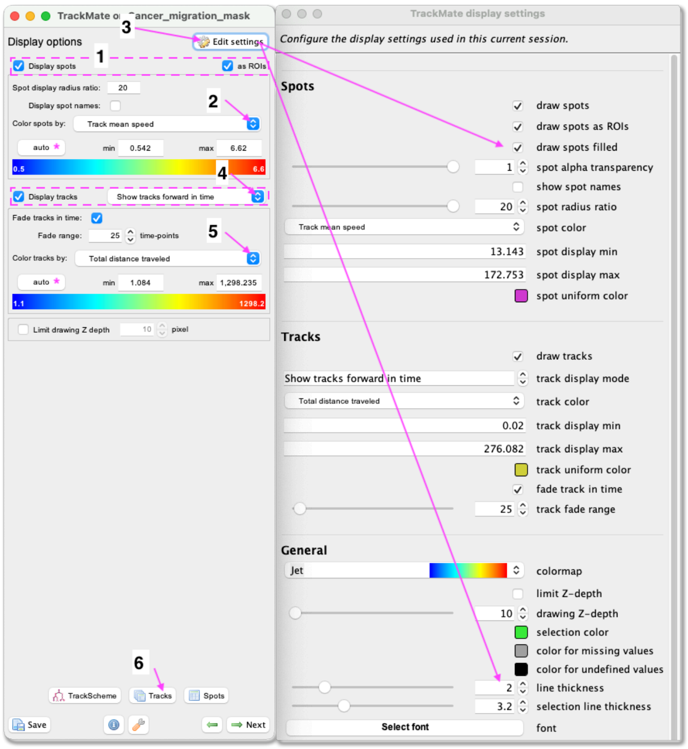

- In this panel, you have access to many plotting features- again, click `Next`.

- Here you have the possibility to do different actions. For this exercise, we will export a tracking image of our experiment. Then from the pull-down menu, select `Capture overlay`. After clicking on `Execute`, a pop-up window opens. Here you can define the time interval you want to save and click `OK`. TrackMate will generate a video of your experiment. Remember to save the image from *File > Save as...*.

You can see on the resulting movie that some touching cells were incorrectly identified as one object. This is a problem with mask images. You can fix this by editing the mask as we did in the first tutorial above or moving back to the raw image and use the StarDist detector we introduce later.

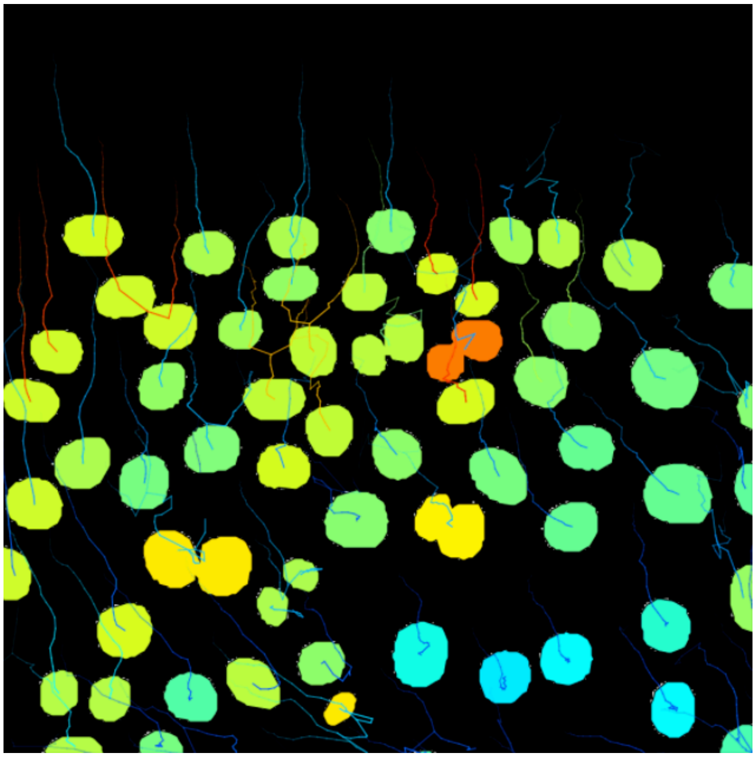

### TrackMate Thresholding Detector.

#### Usage.

This page describes a detector for TrackMate that creates objects from a grayscale image (it can be one channel in a multi-channel image). You must specify a threshold value to segment the objects.

#### Step-by-step tutorial.

Download the tutorial dataset from Zenodo:

DOI 10.5281/zenodo.5220796

- Open Fiji.
- Open your image in Fiji.
- Open TrackMate *Plugins > Tracking > TrackMate*. The start panel will open and display information about your image dimensions. Click `Next`.
- The *Select a detector* panel opens. From the pull-down menu, select `Thresholding detector`. Click `Next`.

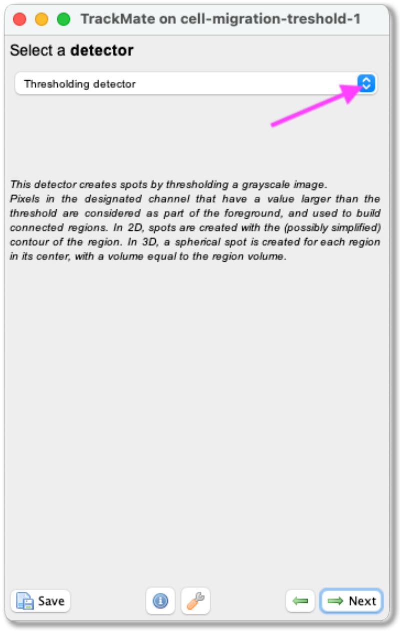

- A panel with the description of the detector opens. You can set the threshold yourself or click on the `Auto` button for automatic thresholding. With our test image, the automatic thresholding value is 92. You can also select to `Simplify the contours` to smoothen the edges of the segmented objects. By clicking on the `Preview` button, you will see a preview of the segmentation. On the left, the number of detected objects is displayed. Once a suitable threshold has been chosen, click on `Next`.

- The objects will be detected. Click `Next` again.

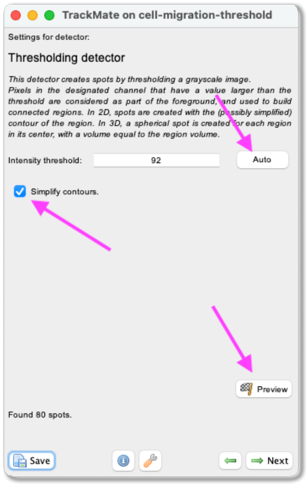

- A panel to filter the detected spots according to their quality opens (more information about this filtering can be found here). With our test image, this part can be ignored. Click `Next`.

- Next, a panel to filter spots according to their properties (*i.e.* size, shape, location, or signal intensity) opens. In this exercise, do the following:

  - Click the green `+` sign at the bottom of the panel - a filter appears.

  - Click the pull-down menu and select `Area`. Here we will filter out the smallest detected objects. Make sure the `Above` button is selected.

  - Drag the horizontal line (pink dashed line) to 20. The small objects become invisible.

- Click `Next`.

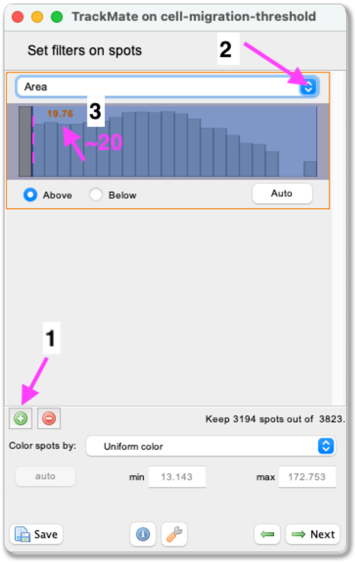

- A tracking panel opens. In this panel, you can select a method for tracking your objects. In this exercise, we use the `LAP tracker`. Please select it from the pull-down menu and click `Next`.

- A panel for the LAP tracker settings opens. First, with the `Frame to Frame linking` parameter, you give the maximum distance to link two objects between frames. Here use **20 microns**. Then, you can choose how many spots can be missing, and they could still be the same track. Tick the `Allow gap closing` box and add values: Max distance: **20 microns** and `Max frame gap`: **4**. Next, you let TrackMate know if the tracks are allowed to split. Splitting can be caused, for example, due to cell division. Tick the box `Allow track segment splitting` and insert value Max distance: **20 microns**. Below you will also see settings for Track segment merging. Here this box should remain unticked. Click `Next`.

- A Track filter panel opens. In this panel, you can filter tracks according to their properties (*i.e.*, length, speed, or location). With our test image, we do not need to filter any tracks. Click `Next`.

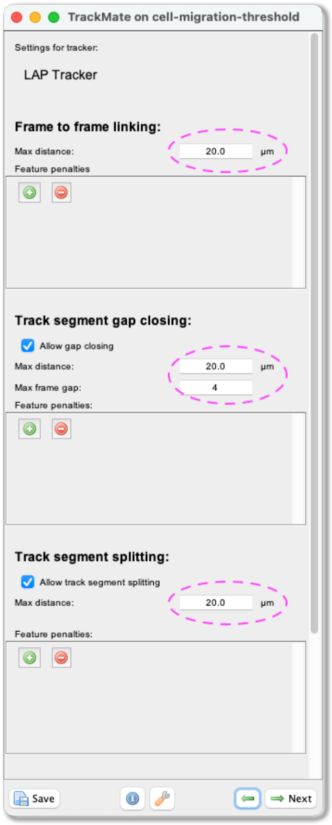

- A Track filter panel opens. In this panel, you can filter tracks according to their properties (i.e., length, speed, or location). With our test image, we do not need to filter any tracks. Click `Next`.

- Next, a panel with multiple display options appears. Here you can define how the objects and tracks are labelled. In this exercise, we will label the objects and tracks according to their mean speed.

- First, make sure that the `Display spots`, `as ROIs` and `Display tracks` boxes, are ticked.

- Select `Track mean speed` from the `Color spots by` and `Color tracks by` pull-down menus and click `auto`.

- In this panel, you can also export the results as CSV files. Please do so by clicking `Tracks` at the bottom of the panel. A window with a lot of data shows up. Ensure you export the `Spots` (information about the spot) and the `Tracks` (information about the tracks) files. Close the results window and click `Next`.

- You will see a plot features panel - again, click `Next`.

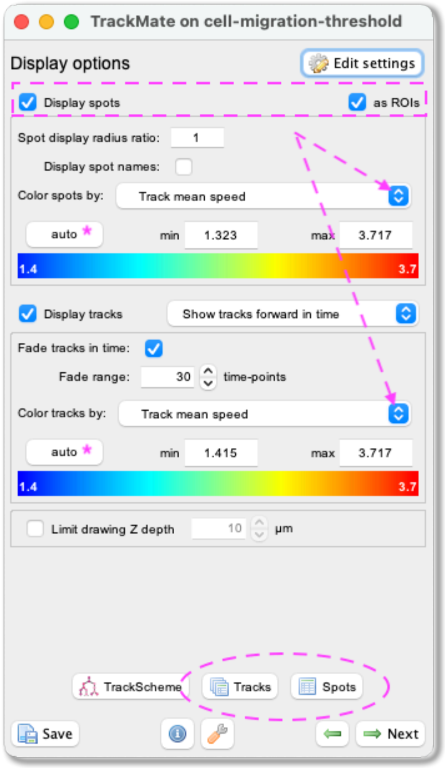

- In the next panel, you can perform multiple operations. For this exercise, we will export a tracking image of our experiment. Then from the pull-down menu, please select `Capture overlay`. As you click on `Execute` below, a pop-up window opens. Here you can define the time interval you want to save. You can hide the image if you wish by ticking the box `Hide image` and click `OK`. TrackMate will generate a video of the experiment. Remember to save the image with *File > Save as...*

### TrackMate Label-Image Detector.

This page describes a detector for TrackMate that creates objects from a label image.

#### Usage.

Label images are especially convenient as an output of segmentation algorithms. Indeed, in some cases you might have different objects that are so close that they touch each other. If a segmentation algorithm can detect them, but outputs a black and white mask, they will appear as one object in the mask if they share a border. In a label image, each object is represented by different integer values. For instance, the object #1 in a label image will be made from all the pixels that have a pixel value of 1, over a black background of 0. Object #2 will have the pixel value 2, etc. This allows resolving them even if they touch each other.

You can use a label image as a channel in your source image, since TrackMate can harness multi-channel image. The image type does not have to be of integer type; but the labels need to be integers even on a float type. This makes it easier to combine the label image with the raw image.

If you have only one channel the preview looks like this:

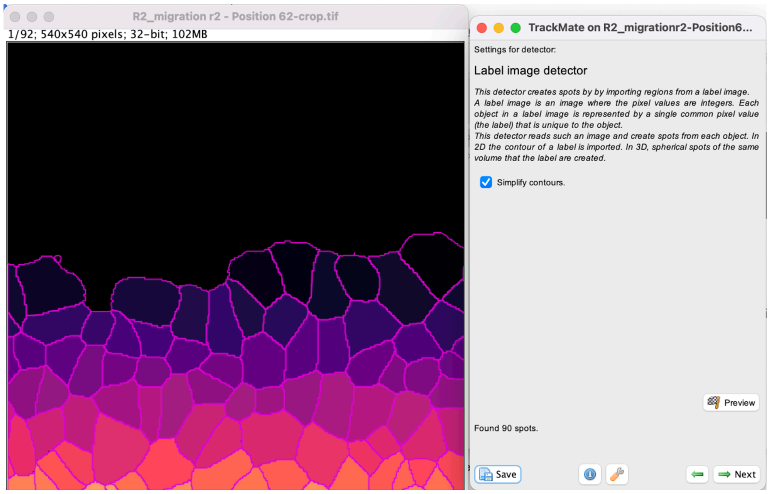

#### Step-by-step tutorial.

First download the tutorial dataset on Zenodo:

DOI 10.5281/zenodo.5221424

- Open Fiji.
- Open your image in Fiji.
- Open TrackMate *Plugins > Tracking > TrackMate*. The start panel will open and display information about your image dimensions. Click `Next`.

- The “Select a detector” -panel opens. From the pull-down menu select the Label Image detector. Click `Next`.

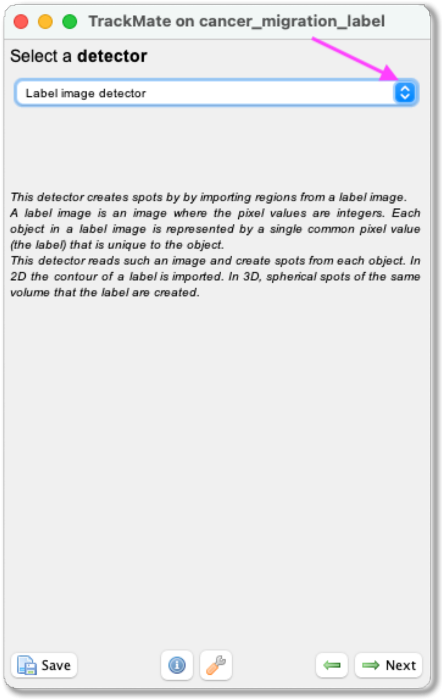

- A panel with the description of the detector opens. Here you can also select to Simplify the contours to smooth the edges of the segmented objects. Click on `Preview` to see a preview of the segmentation. On the left, there will also be the number of detected objects in that field of view. Click `Next`. The objects are now being detected.

- When the progress bar has reached the end, click `Next`.

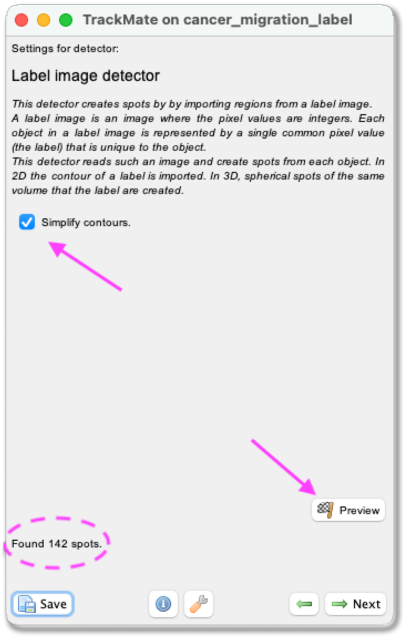

- A panel to filter the detected spots according to their quality opens (more information about this filtering can be found here. With our test image, this part can be ignored. Click `Next`.

- A panel to filter spots according to their properties (i.e., size, shape, location, or signal intensity) opens. With our test image, in frame 65 you will see small floating cells that were segmented. To get rid of this, you can add a spot filter to remove anything with an `Area` lower than 230. Click `Next`.

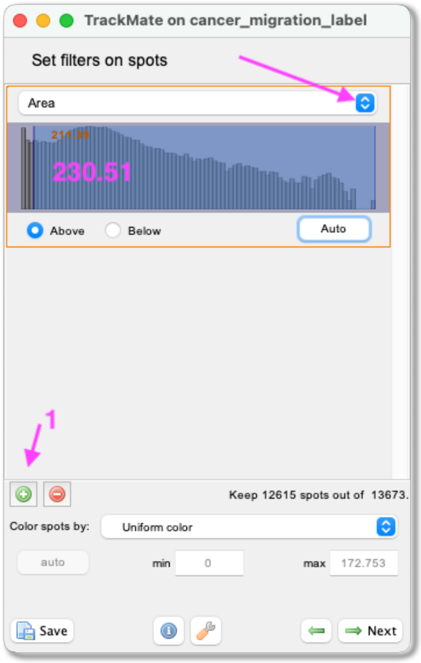

- A tracking panel opens. In this panel, you can select a method for tracking your objects. In this exercise, we use the `LAP tracker`. Please select it from the pull-down menu and click `Next`.

- A panel for the LAP tracker settings opens. First, with the `Frame to Frame linking` parameter, you give the maximum distance to link two objects between frames. With our test image, we used **20 microns**. Then, you can choose how many spots can be missing, and they could still be the same track. Tick the `Allow gap closing` box and add values: `Max distance`: **20 microns** and `Max frame gap`: **5**. Next, you let TrackMate know if the tracks are allowed to split. Splitting can be caused, for example, due to cell division. Tick the box `Allow track segment splitting` and insert value Max distance: **30 microns**. Below you will also see settings for Track segment merging. Here this box should remain unticked. Click `Next`.

- A Track filter panel opens. In this panel, you can filter tracks according to their properties (*i.e.*, length, speed, or location). With our test image, we do not need to filter any tracks. Click `Next`.

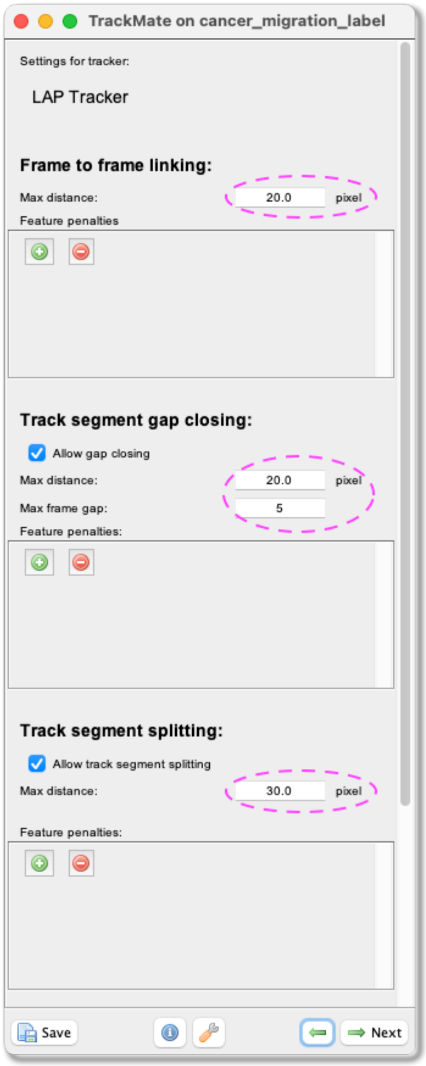

- A panel with the Display options appears. Here you can define how the objects and tracks are labelled.
- First make sure that the `Display spots`, `as ROIs` and `Display tracks` boxes are ticked.
- Select from the pull-down menu `Color spots by Area`, and click `auto`. This allows you to visualize cell size by color (blue smaller cells, red the largest).
- From the pull-down menu, select `Show tracks forward in time`. This will show the future trajectories of the tracked objects.
- from the Color tracks by pull-down menu select `Track mean speed`. And click `auto`.
- Click `Next`.

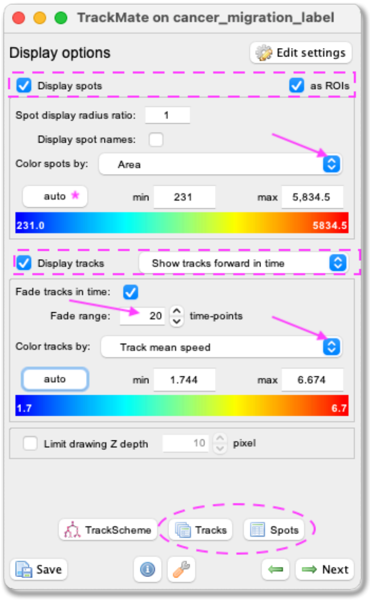

- In this panel you can also export the results as .csv files, please do so by clicking `Tracks` at the bottom of the panel. A window with a lot of data shows up. Make sure you export the `Spots` (information about the spot) and the `Tracks` (information about the tracks) files. Close the results window and click `Next`. You will see a plot features panel - again click `Next`. You will see a plot features panel, where you can generate plots of certain features if you wish - again click `Next`.

- In the last panel, there is a possibility to do different actions. For this exercise, we will export a tracking image of our experiment. From the pull-down menu please select `Capture overlay`. As you click `Execute` below a pop-up window shows up. Here you can define the time interval you want to save and click `OK`. TrackMate will generate a video of the experiment.

- Remember to save the image from [File - Save as...].

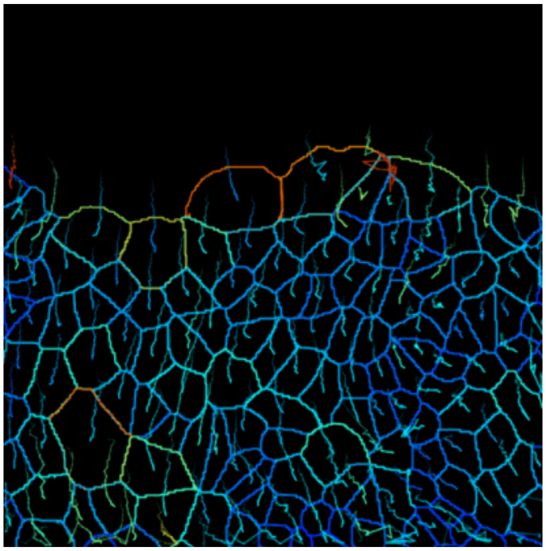

Cool, no?

### TrackMate MorphoLibJ.

This page describes a detector module for TrackMate that relies on the Morphological Segmentation plugin shipped with the MorphoLibJ library. It is not included in the core of TrackMate and must be installed via its own update site.

#### Installation.

We need to subscribe to both the MorphoLibJ library and the TrackMate-MorphoLibJ update sites.

1. First, subscribe to the **IJPB-plugins** update site, that contains the MorphoLibJ library:

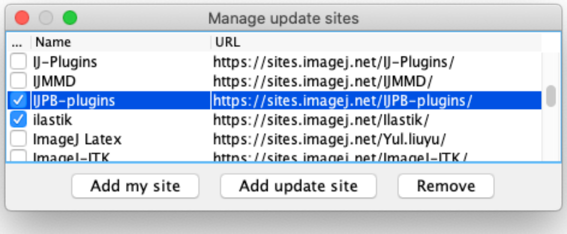

2. Then subscribe to the **TrackMate-MorphoLibJ** update site:

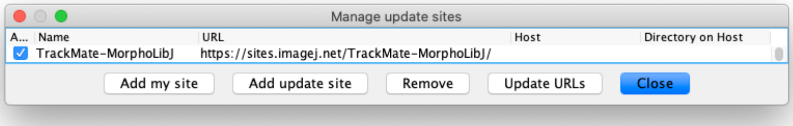

Relaunch Fiji. A new item should appear in the list of detectors in the 2nd panel of the TrackMate UI:

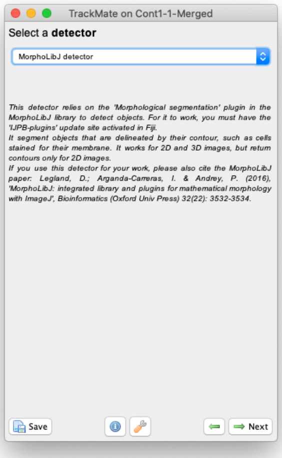

#### Usage.

This detector module literally implements the Morphological Segmentation plugin. It will therefore have the same strengths and limitations. Also, it limits itself to dealing with **Border images**. That is: images in which the objects are labelled for their contours. For instance, cells stained for their membranes.

The best documentation for this segmentation tool is found on its dedicated page, but we will repeat the description of the key parameters in the tutorial below. One of the key advantages compared to using directly the plugin is that the TrackMate module version will properly process time-lapses, while the core plugin treats everything as a Z-stack.

#### Tutorial. Tracking cells in Xenopus tissue.

Download the tutorial dataset from Zenodo.

DOI 10.5281/zenodo.5211585

It contains a movie of cells in a Xenopus tissue, imaged by and courtesy of Jakub Sedzinski (University of Copenhagen).

You will find two movies in the zip file: `Cont1-1.tif` and `Cont1-1-Merged.tif`. The second one is simply a duplicate of the first one, but with a second channel contained a processed version of the raw image. We will use this one for this tutorial.

- Lauch Fiji and open the `Cont1-1-Merged.tif` file. Then launch TrackMate *Plugins > Tracking > TrackMate*

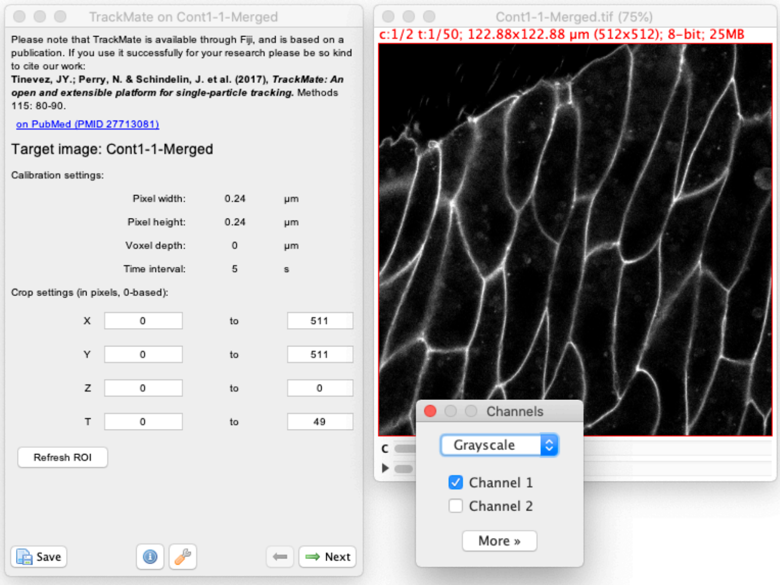

- Click `Next` and select the `MorphLibJ detector` in the detector list (see the image above). Click `Next` again.

- You now see the configuration panel for the detector.

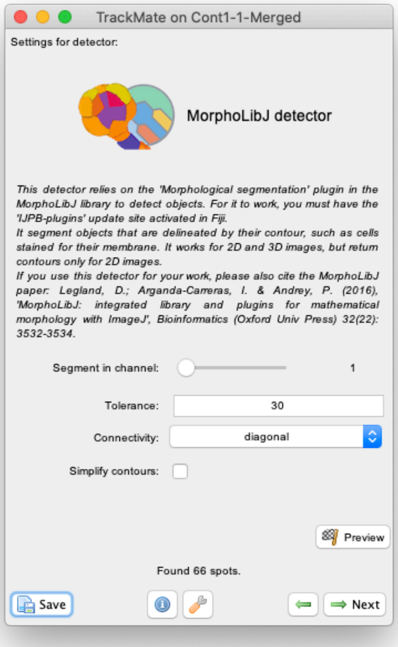

- Beyond the `Segment in channel` slider and the `Simplify contours` checkbox that are specific to TrackMate, it offers to tune the two key parameters of the Morphological Segmentation plugin:

  - The `Tolerance` parameter. Copying from the plugin page: This is the dynamic of intensity for the search of regional minima (in the extended-minima transform, which is the regional minima of the H-minima transform, value of h). Increasing the tolerance value reduces the number of segments in the result, while decreasing its value produces more object splits. Note: since the tolerance is an intensity parameter, it is sensitive to the input image type. A tolerance value of 10 is a good starting point for 8-bit images (with 0-255 intensity range) but it should be drastically increased when using image types with larger intensity ranges. For example, to ~2000 when working on a 16-bit image (intensity values between 0 and 65535).

  - The `Connectivity` selector specifies the desired pixel connectivity. `Diagonal` specifies the 8-connectivity (in 2D) and `Straight` specifies the 4-connectivity. This parameter has a limited impact for large objects. Selecting non-diagonal connectivity (4 or 6) usually provides more rounded objects.

The `Tolerance` parameter is the one that matters the most. If we play with some values on the first channel, you will see that we are kind of stuck in a not so nice place. For large values of the tolerance, we end in missing some borders, fusing cells together. For smaller values, we have the opposite effect and have many spurious small objects that appear.

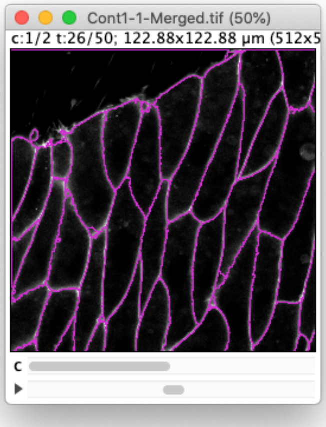

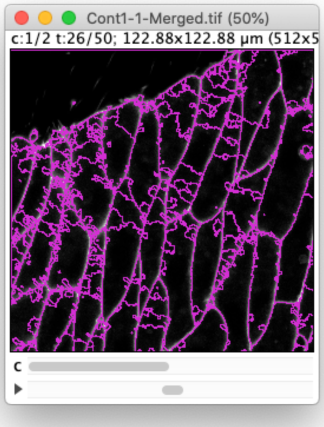

(Results of the preview with a tolerance of 30 and 10 respectively.)

Ideally, we want to find a value in between where there is not missed border and no spurious objects, but despite the high quality of this movie, this is not perfectly possible. (A value of 20 gives satisfactory results, but not through the whole movie). Indeed, there is some noise in the image that negatively impacts the resulting segmentation. A solution is to exploit the integration of TrackMate within Fiji to pre-process the movie to temper the effect of noise. I have found out that a 3x3 median filter followed by a gaussian blue with σ=0.5 pixels give satisfactory results with the least effort. (It is still not good enough to have 100% perfect segmentation results. For this you want to have a cleverer edge-enhancing strategy such as using a LoG filter properly tuned. But we move forward.)

Because we want to go quick on this tutorial series, I have included the results on the preprocessing in the image we have been using in the second channel. Therefore, simply set the `Segment in channel` slider to 2, use a tolerance value of 15 and select the `Simplify contour` checkbox. Even with this our results are not 100% perfect. We will have to discard some objects and tracks later.

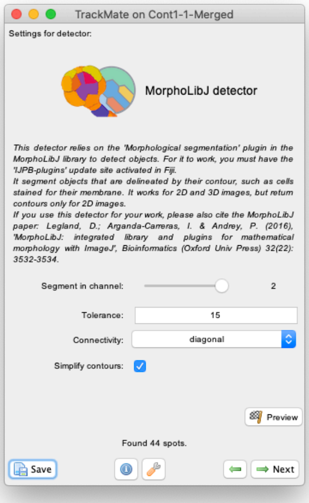

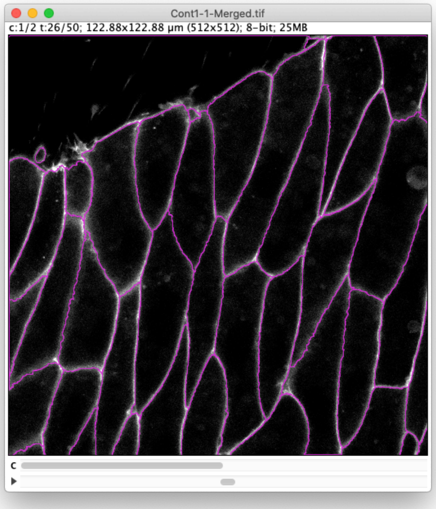

- Then click `Next` to let the processing of all frames proceed. In the end you should have about 2000 spots.

- In the `Initial thresholding` panel, we can discard objects of low quality. For the MorphoMorphoLibJlibJ detector **the object quality is simply the object area in pixels**. So we can use this step to prune small objects resulting to catchment basins caused by remaining noise. Try to put the slider around 766 as below, which will leave about 1800 objects in the results.

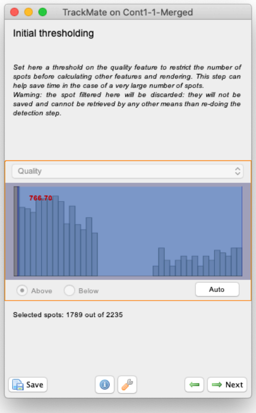

- Then click `Next`. TrackMate will first compute all the features of the retained objects, including morphological features. As our cells are relatively big this may take several seconds.

- In many cases, such as this one, you will have in all time-points a very large object corresponding to the background. We want to filter it out so that it is not part of the tracking. In this panel, click on the greent + button to add a filter, and in the filter panel that appear, choose `Area` in the top list. The histogram of all the objects area appears. You can see that it is made of two parts well separated, one for small area around 500 µm² and a peak around 2000 µm². (Note that now the area is expressed in µm² and not in pixels as in the *Initial thresholding* panel.) The latter corresponds to the background objects, as you can check by setting the threshold line in between the two peaks. If the radio button below the histogram is set to `Above`, TrackMate will hide all the smaller objects and only one object per frame should remain. We want to keep the smaller objects, so select the `Below` radio button to hide the background objects.

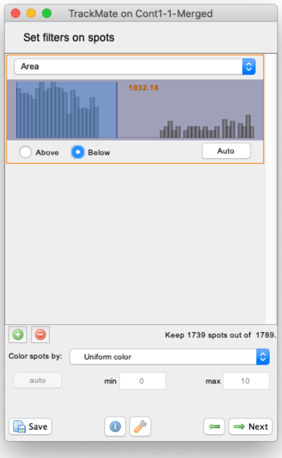

- Click `Next`. We need now to select a tracker. For this kind of large objects, that moves by less than their diameter, the `Overlap tracker` is well suited. Select it and click `Next`.
- The default parameter values are good enough in our case (`Precise`, `Min IoU` of 0.3 and `Scale factor` of 1), so click `Next` directly.
- We find 38 tracks with a quite large mean number of spots:

```
Starting tracking process.
Tracking done in 1.8 s.
Found 38 tracks.
  - avg size: 45.6 spots.
  - min size: 2 spots.
  - max size: 50 spots.
```

- Let's skip filtering them and move on to the *Display options* panel, then to the *Plot features* panel.
We will inspect the tracking results and check what is the impact of the over-segmentation we had with our segmentation parameters. For instance, select the large object which is roughly 3rd starting from the top-right (see below). At the frame 10, this cell was cut in two because of over-segmentation. Let's plot the resulting cell are over time.
  - Make sure the `Spots` pane is currently selected in the grapher panel.
  - Make sure that `T` is selected as a feature the X axis.
  - As feature for the Y axis, select `Area`.

– We don't want to plot the values for all cells, just for the one we selected. So, at the bottom of the grapher panel, click on the `Track of selection` radio button.
  We see that around T=50s, the area drops brutally, showing that the corresponding object was cut in half at this time-point.

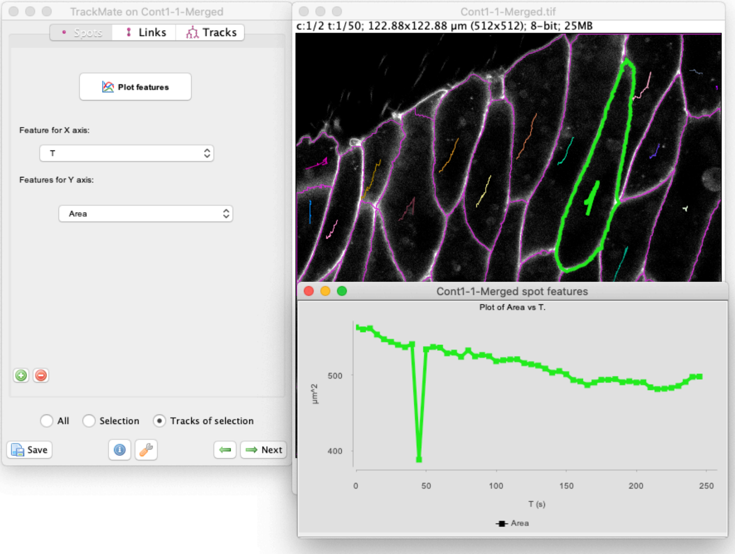

A proper way to fix this would be to merge the two objects belonging to the same cell before tracking.
But as of today TrackMate does not offer a way to do so, so we are left to post-processing results, or finding a pre-processing pipeline that gives a perfect segmentation.

But apart from the few segmentation errors in this movie, the resulting segmentation is pleasing:

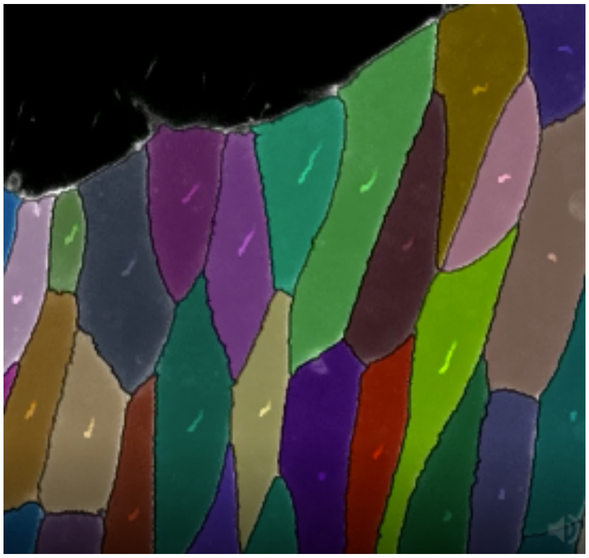

Here is how to generate this movie.

- Click the green left arrow key at the bottom of the TrackMate panel to move back to the `Display options` panel.
- Click on the `Edit settings` button in the top-right corner. The following display config panel shows up:

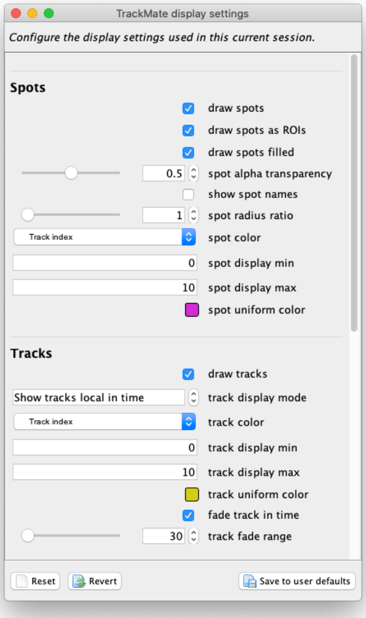

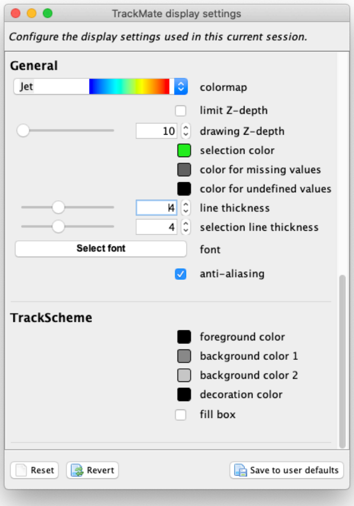

Let's configure it as follow. From top to bottom:

- Select the `draw spot filled` checkbox so that in the overlay the objects are painted filled.
- For `spot alpha transparency`, put 0.5, so that we see the raw image by transparency.
- In the `spot color` list, select `Track index`, so that objects are colored by the track they belong to.
- In the `track display mode`, select `Show tracks local in time`, so that we see tracks as a "dragon tail" backward and forward in time.
- In `line thickness`, put a value of 4, to stress the objects contour.

Now we also want to only display one channel in grayscale instead of the two channels in composite at once.
In the Fiji menu, show the channels tool dialog *Image > Color > Channels Tool...*In the dialog that appears, select `Grayscale` and the `Channel 1`. The image and TrackMate overlay should look like this:

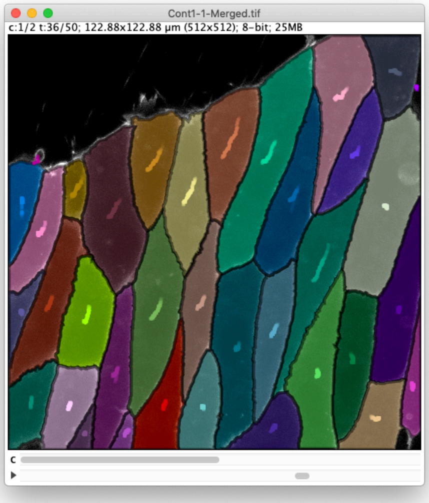

- Close the *TrackMate display settings* panel.
- Now move back to the last panel of TrackMate, by clicking on the `Next` button two times in the TrackMate wizard. This should bring you to the *Select an action* panel.
- Select the `Capture overlay` action and click the `Execute` button.
- In the dialog that appears, click `OK`. This will generate a new RGB movie that you can save to AVI for instance.

And that's it for this tutorial.

### TrackMate Weka.

This page describes a detector module for TrackMate that relies on the Trainable Weka Segmentation plugin to segment objects in 2D or 3D. It is not included in the core of TrackMate and must be installed via its own update site.

#### Installation.

For this module to work, you just need to install the TrackMate module. Subscribe to the **TrackMate-Weka** update site.

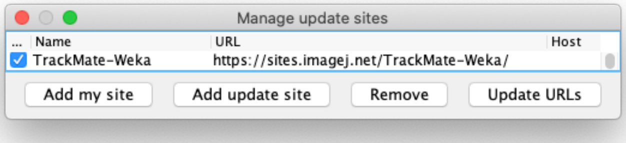

#### Tutorial: Tracking focal adhesions.

Trainable Weka Segmentation is a machine learning pixel-based segmentation, a classifier for the objects of interest in the image is trained from user annotation, the classifier is then used for segmentation. The following image is employed to demonstrate the usage of the Weka classifier within TrackMate and can be downloaded from Zenodo:

DOI 10.5281/zenodo.5226842

The movie included show human dermal microvascular blood endothelial cells expressing Paxillin, imaged with a spinning-disk confocal microscope.

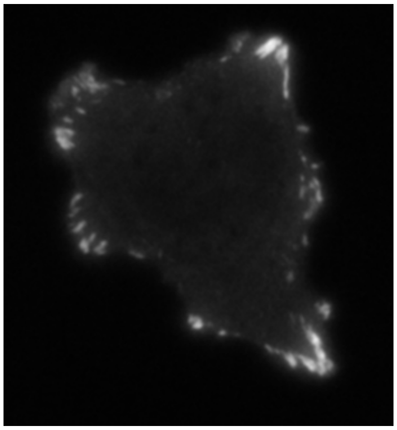

The goal is to track the focal adhesions staying at the cell periphery. They are in general brighter than the cell body and the image background, but with a high variance in their intensities.

#### Weka classifier.

We provide in the tutorial dataset an already trained Weka classifier for detection of the focal adhesions.

Only the first image frame was extracted to do the annotation. The default features plus the Gabor filter were selected for training. The figure below illustrates the Weka GUI with two classes: one in red for focal adhesions and one in green for the others (cell body, image background). More details on training the classifier can be found from on the plugin documentation page.

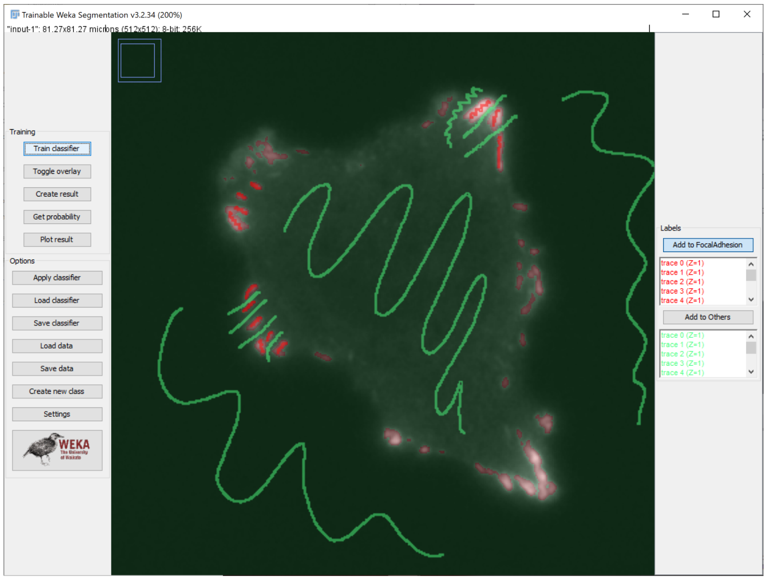

#### Using Weka in TrackMate.

Now open the image in Fiji and launch TrackMate, click Next and select Weka detector from the dropdown menu

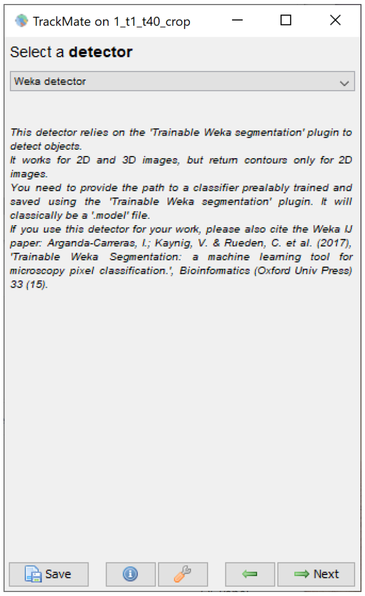

In the next panel, browse the classifier file, choose the target class as `FocalAdhesion` and set the threshold probability as 0.5.

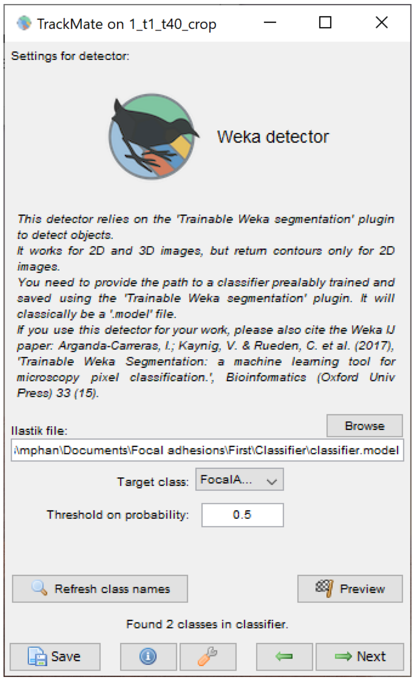

Click on `Preview` button to check the segmentation result, then click Next to run the segmentation for all time frames. This step takes about 8 minutes on a standard pc.

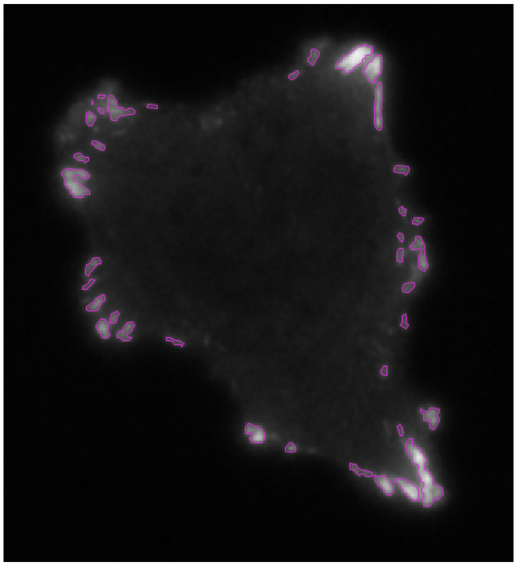

Once the detection process is done, click `Next` until the step `Select a tracker`, here we choose the `Simple LAP tracker` and set the `Linking max distance` and the `Gap-closing max distance` are both equal 2.0 microns, the `Gap-closing max frame gap` is set as 2. Click `Next` and wait for the tracking process to finish.

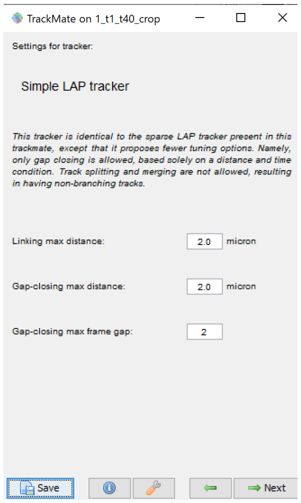

One purpose of tracking the focal adhesions is to study how they shrink or expand over time. TrackMate provides an option to color the tracks by time. Click Next until the step Display options and select Frame in Color tracks by, then click auto.

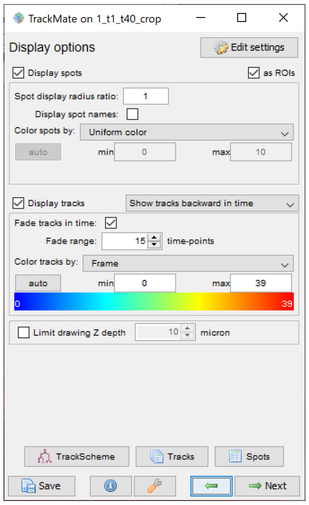

Looking at the tracking result shown in the movie below, the focal adhesions localizing at the top are shrinking whereas the ones at the bottom right are expanding.

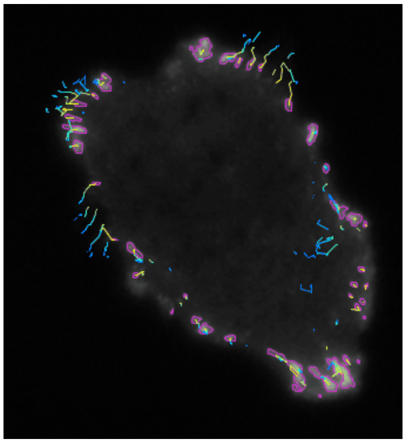

### TrackMate StarDist.

This page describes a detector module for TrackMate that relies on StarDist to segment objects in 2D. It is not included in the core of TrackMate and must be installed via its own update site.

#### Installation.

You need to subscribe to the **CSBDeep** update site and to the **StarDist** update site first, then to the **TrackMate-StarDist** update site:

The CSBDeep update site:

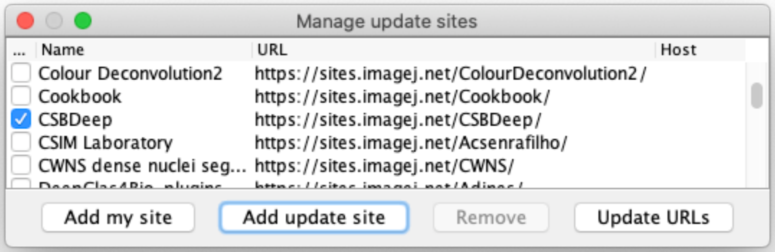

The StarDist update site:

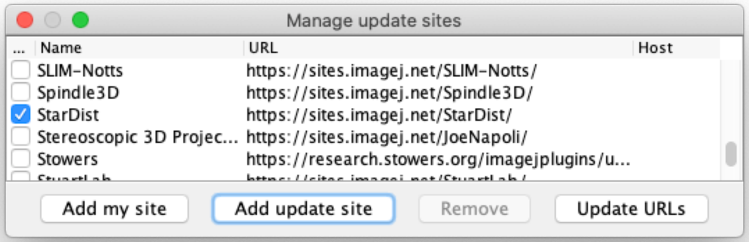

And finally, the TrackMate-StarDist update site.

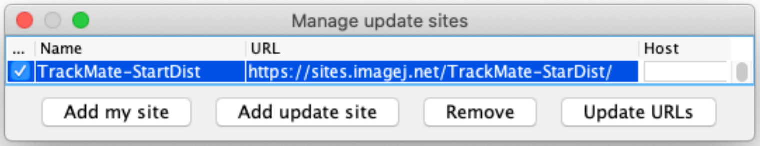

Once you have them all, please restart Fiji.

We suggest that you test StarDist by itself just to make sure it runs after installation. This way we can better find what is wrong in case it does not work. Follow for instance the instructions on the StarDist page.

TrackMate-StarDist ships two detectors that will appear in TrackMate. After installing TrackMate-StarDist and restarting Fiji, these two detectors will be integrated in TrackMate in a transparent manner. We describe how to use them in the four tutorials below. They describe in order:

1. Using TrackMate-StarDist on a single-channel image.
2. Using TrackMate-StarDist on a multi-channel image and exploiting the intensity information.
3. Using TrackMate-StarDist with a custom Deep-Learning model.
4. Using TrackMate-StarDist to segment a 3D image using a slice-by-slice approach.

If you use this detector for you research, please be so kind as to cite the StarDist and the TrackMate papers:

> Schmidt, U., Weigert, M., Broaddus, C., & Myers, G. (2018). Cell Detection with Star-Convex Polygons. In *Medical Image Computing and Computer Assisted Intervention – MICCAI 2018* (pp. 265–273). Springer International Publishing. doi:10.1007/978-3-030-00934-2_30

#### StarDist detector with builtin versatile nuclei model on a single channel image.

The StarDist plugin comes with a very efficient model that can segment nuclei imaged in fluorescence in 2D, generated from the dataset in the Kaggle challenge of 2018:

> Caicedo, J. C., Goodman, A., Karhohs, K. W., Cimini, B. A., Ackerman, J., Haghighi, M., … Carpenter, A. E. (2019). *Nucleus segmentation across imaging experiments: the 2018 Data Science Bowl. Nature Methods*, *16*(12), 1247–1253. doi:10.1038/s41592-019-0612-7

We use this model in the first StarDist detector.

In the first tutorial we will use a movie following the migration of cancer cells, labelled for their nuclei. You can download it from Zenodo:

DOI 10.5281/zenodo.5206107

First launch Fiji and open the tutorial image in Fiji. We use here the `P31-crop.tif` file, provided in the tutorial data.

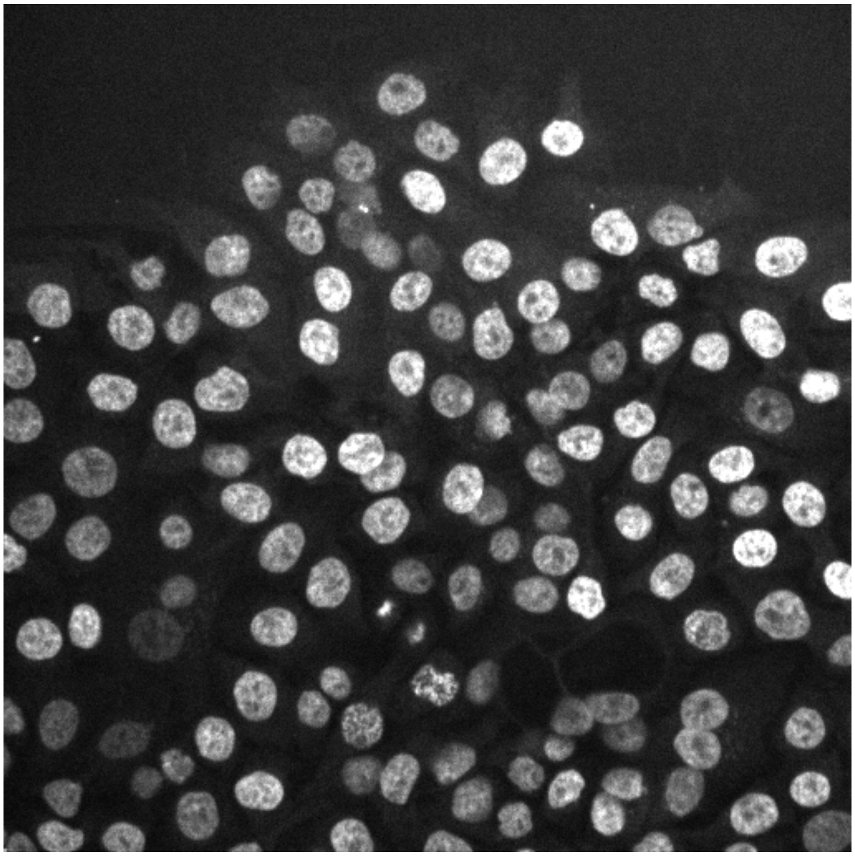

Then launch TrackMate: *Plugins>Tracking>TrackMate*.

In the second panel titled **Select a detector**, you should see two new choices in the list, and one of them is **StarDist detector**.

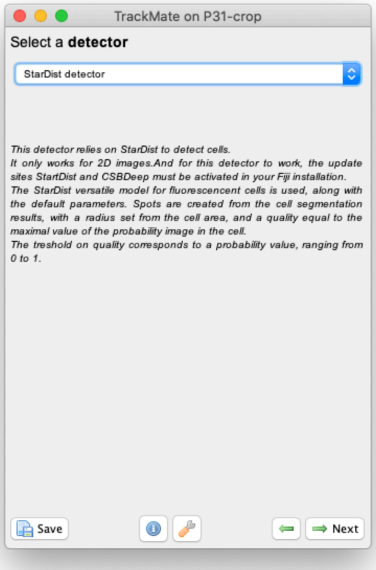

Select it and click **Next**. This simple panel appears.

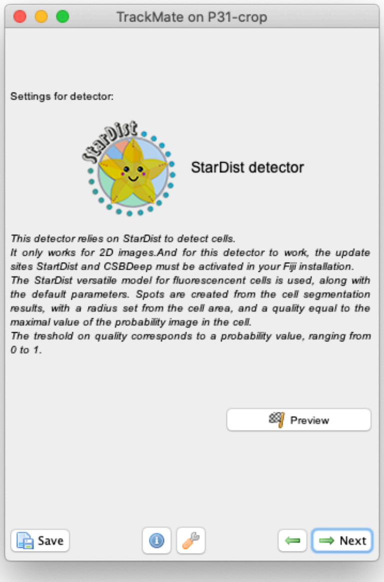

Note that in this panel there is no configuration for Stardist detector: we use the default values of this model for the score threshold and overlap threshold. We observed, that in cases when this model with the default values does not work well with some dataset, changing the score and overlap threshold have little to no positive impact on the results. We reasoned therefore that if the model and the default values do not work well for your data, it is best then to train a specific StarDist model for your problem.

Check the results of segmentation by clicking on the **Preview** button. Here is what I get on the first time-point of the tutorial image:

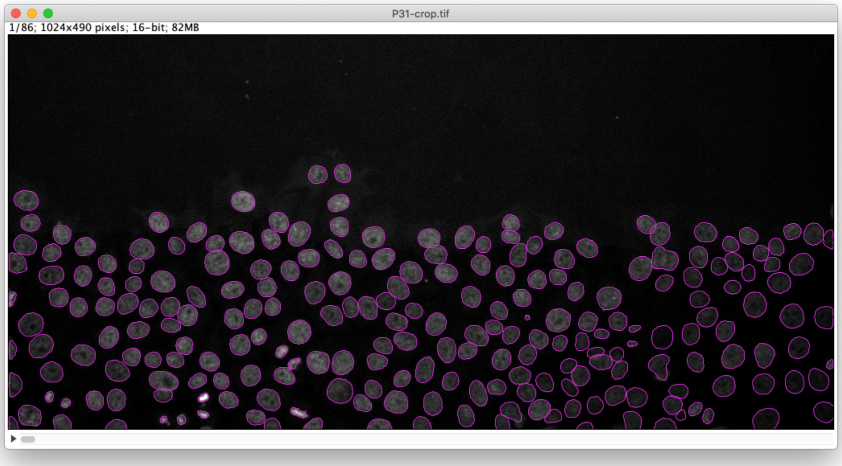

The StarDist model works really well with this kind of data.

After that you follow through the next steps in TrackMate to segment all cells in all time-points then track them. Using the default tracker and default parameters each time we get this result:

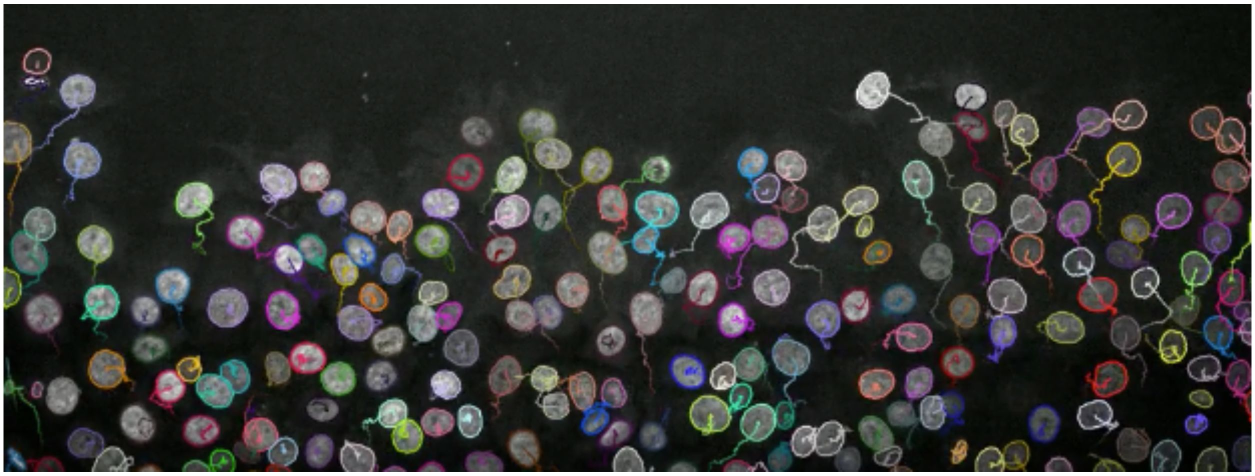

It was simple and fast, which means that careful inspection for missed cell divisions and false links is on order.

#### ERK signaling and motility assay with a multi-channel image.

In this part of the tutorial, we will correlate the translocation of an ERK reported in the nuclei with cell motility. We will use images from a cell migration assay, where cells are expressing an ERK reported in the first channel and are stained for their nuclei in the second channel. The analysis will consist in segmenting and tracking the cells in the nuclei channel and analyzing intensities in the ERK channel. In a second part we will investigate whether the instantaneous speed correlates with the ERK signal in the nuclei.

You can find the source image and additional files on Zenodo:

DOI 10.5281/zenodo.5205955

##### Tracking with TrackMate.

Step by step:

- Open Fiji.
- Open your image. This image has two channels, the ERK reporter in channel 1 (green) and a DNA staining in channel 2 (grey).
- Open TrackMate: *Plugins>Tracking>TrackMate*. The TrackMate start panel will open, showing information about the image dimensions. Click `Next`.

- The “Select a detector” panel opens. From the pull-down menu, select the `StarDist detector`. The description of the detector method will appear in the panel. Click `Next`.

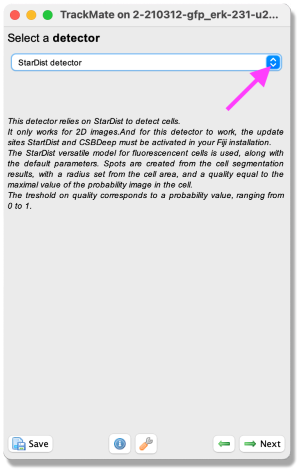

- A panel with a description of the StarDist detector opens. By clicking on the `Segment in channel` slider, select the channel to segment. Here, select channel 2, which contains the DNA label.
- By clicking the `Preview` button, you can test the detector on the current frame.
- When you are happy with the result, click `Next`.
- The detector will detect all nuclei in the selected channel for all time frames. As the movie has many time-points, this can take a few minutes.
- When the progress bar has reached the end, click `Next`.
- A panel to filter the detected spots according to their quality opens (more information about this filtering can be found here). In this exercise, this part can be ignored. Please click `Next`.

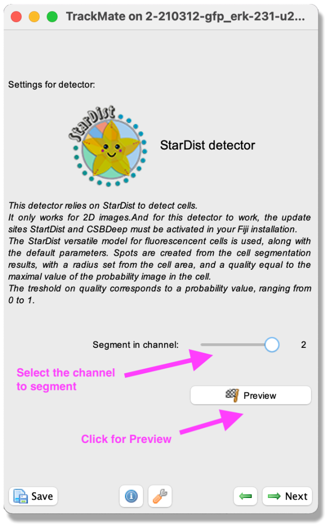

- A panel to filter spots according to their properties (*i.e.*, size, shape, location, or signal intensity) opens. In this exercise, do the following:
  - Click the green plus sign at the bottom of the panel - a filter appears.
  - Click the pull-down menu and select `Area`. Here we will filter out the smallest detected objects. Make sure the `Above` button is selected.
  - Drag the horizontal line (pink dashed line) to value 30.92 (approximatively). If you want to enter an exact value as threshold, just click inside the filter panel to make it active (the text becomes red) and type the number you want with keyboard. After 1 second it will be set as threshold.
  - Click again on the green plus sign at the bottom of the panel to activate a second filter.
  - Click the pull-down menu and select `Mean intensity ch2`. Here we will filter out the detected objects with low intensity in Channel 2. Make sure the `Above` button is selected. Drag the horizontal line (pink dashed line) to value 120.73. Click `Next`.

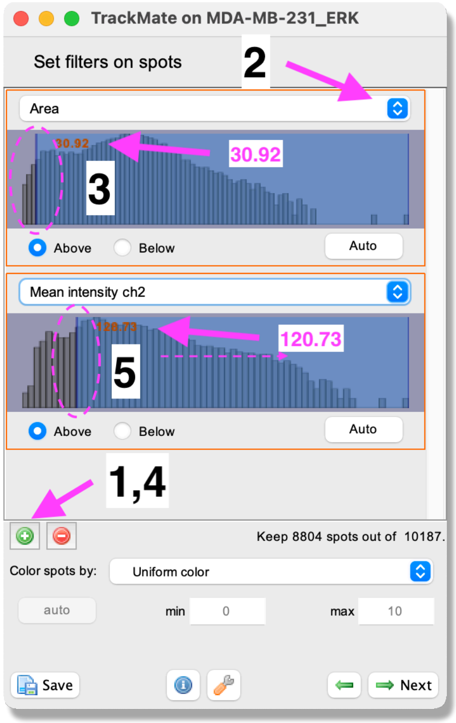

- Next, a tracking panel opens. In this panel, you can select the methods for tracking objects. Here, we use the `LAP tracker` to account for possibly dividing cells. Please select it from the pull-down menu and click `Next`.

- A panel for the LAP tracker settings opens. Int this panel, you choose how to track the cells. First, with the `Frame to Frame linking` parameter, you give the maximum distance to link two objects between frames. Here use 30 microns. Then, you can choose how many spots can be missing, and they could still be the same track. Tick the `Allow gap closing` box and add values: `Max distance`: 30 microns and `Max frame gap`: 5. Next, you let TrackMate know if the tracks are allowed to split. Splitting can be caused, for example, due to cell division. Tick the box `Allow track segment splitting` and insert value `Max distance`: 15 microns. Below you will also see settings for `Track segment merging`. Here this box should remain unticked. Click `Next`.

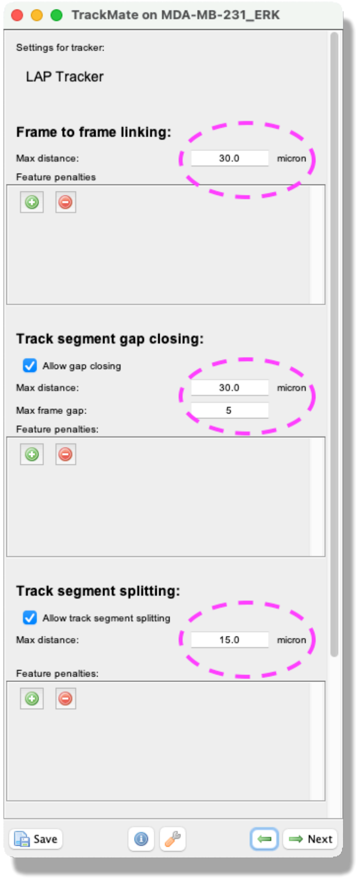

- After tracking, a track filter panel opens. In this panel, you can remove tracks according to their properties (i.e., length, speed, or location). Here, we filter out short tracks to remove cells that migrate in or out of the field of view. As for the object filtering step described above, click the green plus sign to add a filter. Next, click on the pull-down menu and select `Track duration` from the list. Make sure that `Above` is ticked and set the slider to about 30k. Click `Next`.

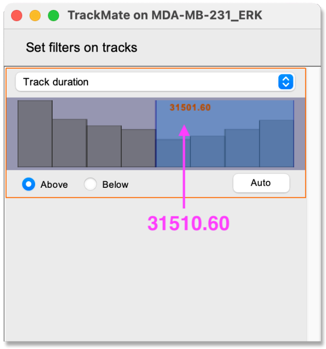

- A window with track visualization options opens. Here it is possible to edit track or object colors according to their properties.
  - First, make sure that the `Display spots` and `as ROIs` boxes are ticked.
  - One option is to label the spots (nuclei) according to the standard deviation of channel 1 intensity. In this case, a blue color will indicate that the ERK activity reporter signal is stable over time. In contrast, a red color will show that the ERK activity reporter signal fluctuates over time. With this coloring, it is possible to visualize cells displaying ERK activity oscillations.
  - To do this, select from the pulldown menu `Color spots by Mean intensity ch1`. Click `auto` below to spread out the signal color.
  - For the tracks, we will visualize the track displacement. First, make sure you have the `Show tracks local in time` selected from the pull-down menu after `Display tracks`.
  - Next, tick the `Fade tracks` in the time box and select a `Fade range` of 14. Next, choose from the pull-down menu: `Color tracks by Track displacement`. This will show the tracks that move a short distance in blue and the longest travelled track in red. Now you can visualize if the presence of ERK oscillation corresponds to the distance travelled by the cells.
- In this panel, you can also export the results as .CSV files. Please do so, we will need them for the analysis in MATLAB in the second part. Click on the `Tracks` button at the bottom of the panel. A window with a lot of data shows up. Ensure you export the `Spots` (information about the spot) and the `Edges` (information about the links between 2 spots) files. Close the results window and click `Next`.

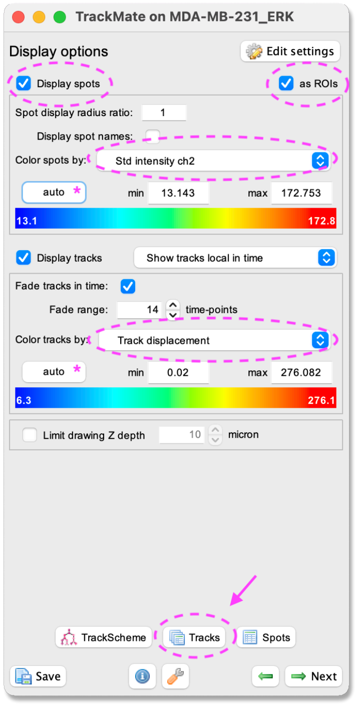

- In the next panel `Plot features`, it is possible to Plot some of the features of the tracks, for example, the changes of the signal intensity over time. However, we cannot readily correlate the instantaneous speed with the ERK intensity in nuclei, because the former is a feature of edge objects and the latter of spot objects. We will address this in a second part with a MATLAB analysis script. Click `Next`.

- In the next panel, there is a possibility to do different actions. For this exercise, we will export a tracking image of our experiment. From the pull-down menu, please select `Capture overlay`. As you click on `Execute` below, a pop-up window opens. Here you can define the time interval you want to save. Please tick the box `Hide image` and click `OK`. TrackMate will generate a video of the experiment.

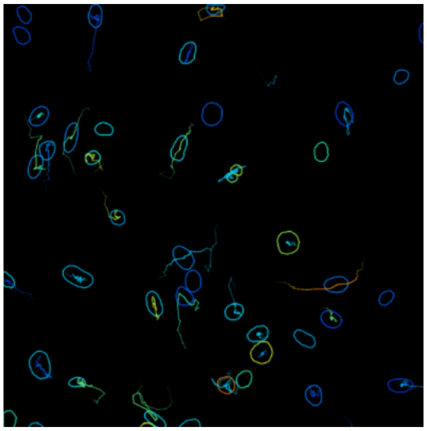

#### Track analysis with MATLAB.

The zip file of the dataset contains the 2 CSV files generated from the first part above, and a MATLAB script that reads them and investigate whether the speed correlates with ERK intensity. We outline the script here.

First let's load the data.

```
>> clear
>> close
>> clc

% Path to the CSV files, exported from TrackMate.
>> spots_csv_file = 'CellMigration_WithERK-spots.csv';
>> edges_csv_file = 'CellMigration_WithERK-edges.csv';

%% Load CSV files into MATLAB tables.
>> spot_table = readtable( spots_csv_file );
>> edge_table = readtable( edges_csv_file );

%% Display info on tables.
>> fprintf( 'Header of the spot table:\n' )
>> head( spot_table )

>> fprintf( '\nHeader of the edge table:\n' )
>> head( edge_table )

Header of the spot table:

ans =

  8×38 table

Var1           ID      TRACK_ID    QUALITY    POSITION_X …
___________    _____   ________    _______    __________
{'ID20994'}    20994      0        0.87727      467.92
{'ID10754'}    10754      0        0.83571      451.67
{'ID25602'}    25602      0        0.89615      490.18
{'ID20995'}    20995      0        0.87711      482.46
{'ID4611' }     4611      0        0.85884      458.63
{'ID7684' }     7684      0        0.83752      468.28
{'ID29701'}    29701      0        0.87407       480.8
{'ID15878'}    15878      0        0.83024      461.84
```

```
Header of the edge table:

ans =

  8×13 table

Var1                     TRACK_ID    SPOT_SOURCE_ID  …
____________________     ________    ______________
    {'ID31545 → ID31681'}       0            31545
    {'ID19256 → ID19400'}       0            19256
    {'ID14361 → ID14477'}       0            14361
    {'ID17824 → ID18039'}       0            17824
    {'ID17160 → ID17216'}       0            17160
    {'ID7684 → ID7827'  }       0             7684
    {'ID10506 → ID10604'}       0            10506
    {'ID22343 → ID22544'}       0            22343
```

As said above, we need to correlate a value that belongs in two different tables: the mean ERK intensity in the spot table, and the speed in the edge table. The speed is defined as an edge feature, because you need two spots to define a displacement and a time interval, hence a speed. But we want to plot the speed and intensity for the same spot. The trick is to use the spot `ID` value, which is unique in a tracking session. The edge table has two columns that store the ID of the 2 spots it links: `SPOT_SOURCE_ID` and `SPOT_TARGET_ID`.

So, we need to join the 2 tables, using the `ID` column in the spot table and (for instance) the `SPOT_TARGET_ID` in the edge table. In MATLAB, this is done as follow:

```
>> T = join( edge_table, spot_table, ...
    'LeftKeys', 'SPOT_TARGET_ID', 'RightKeys', 'ID', ...
    'LeftVariables','SPEED', 'RightVariables','MEAN_INTENSITY_CH1');
>> head(T)

ans =

  8×2 table

      SPEED      MEAN_INTENSITY_CH1
    _________    __________________

    0.0047378          917.44
     0.010687          787.33
    0.0077653          862.49
    0.0037531          907.65
     0.030507          1016.8
    0.0028973          1606.6
    0.0064104          1060.7
     0.030591          1063.9
```

And finally, we can visualize whether we have some correlation with a scatter plot.

```
>> figure
>> scatter( T.SPEED, T.MEAN_INTENSITY_CH1, 'k.' )
>> xlabel( 'Speed (µm/sec)' )
>> ylabel( 'ERK nuclei intensity' )
```

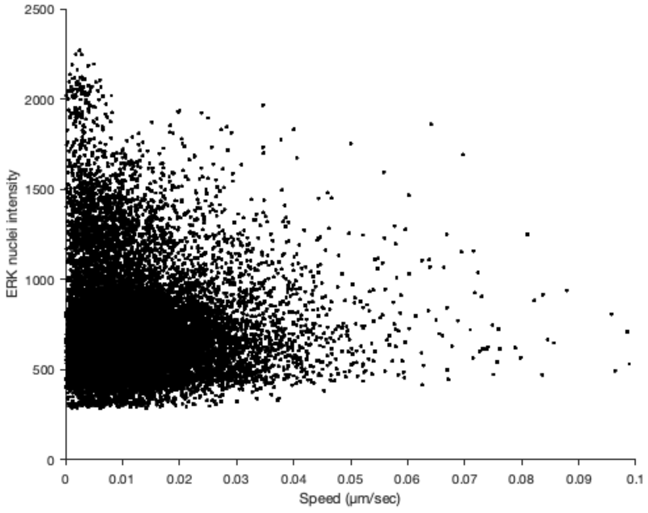

It's rather unclear with that many points. Let's try to plot the density of points.

```
>> figure
>> histogram2( T.SPEED, T.MEAN_INTENSITY_CH1, 'FaceColor', 'flat' )
>> view( 2 )
>> grid off
>> box off
>> xlabel( 'Speed (µm/sec)' )
>> ylabel( 'ERK nuclei intensity' )
```

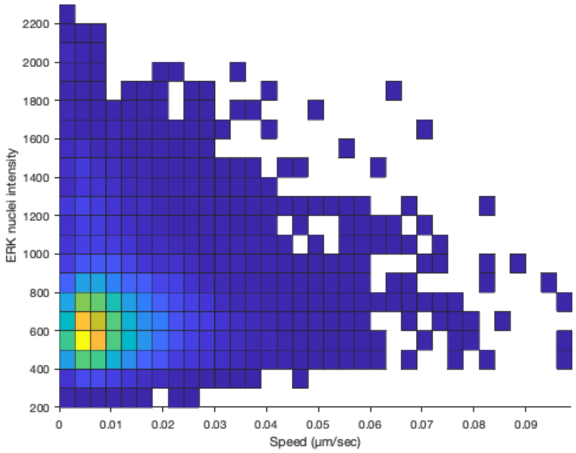

It's not clearer. Let's analyze the correlation coefficient.

```
>> [ R, P ] = corrcoef( T.SPEED, T.MEAN_INTENSITY_CH1 );

>> fprintf( 'Correlation coefficient: %.2e\n', R( 1, 2 ) )
>> fprintf( 'P-value for the correlation: %.2f\n', P( 1, 2 ) )

Correlation coefficient: 4.00e-03
P-value for the correlation: 0.56
```

We must conclude that this dataset does not show a correlation between instantaneous speed and ERK intensity in the nucleus.

#### Tracking T-cells imaged in bright-field with a custom model in the StarDist detector.

You can also use a custom model, that you have trained yourself and packaged as a zip file. We recommend using the dedicated notebooks on the *ZeroCostDL4Mic* platform to do so. Check the ZCDL4M wiki page dedicated to training StarDist to generate such a model.

In this tutorial we will track T cells imaged in bright-field with a model we trained ourselves. You can find the image and the model (packaged as a zip file) on Zenodo:

DOI 10.5281/zenodo.5206119

First open the tutorial image called `TCellsMigration.tif` in Fiji provided in the tutorial data.

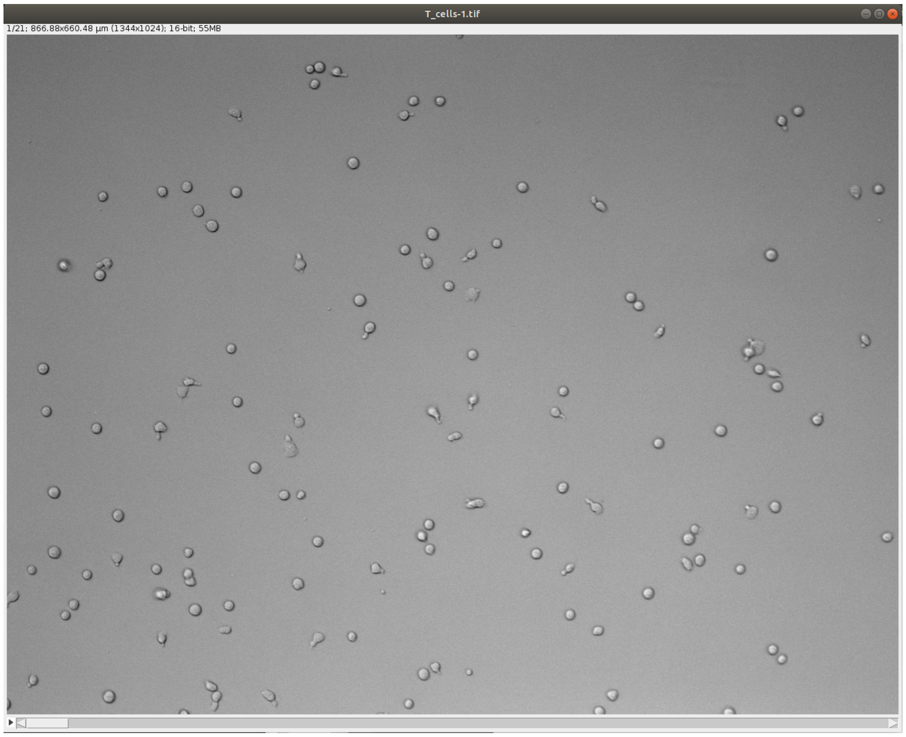

Launch TrackMate. In the second panel titled **Select a detector**, choose **StarDist detector custom model** and click **Next**. Its configuration panel requires several parameters:

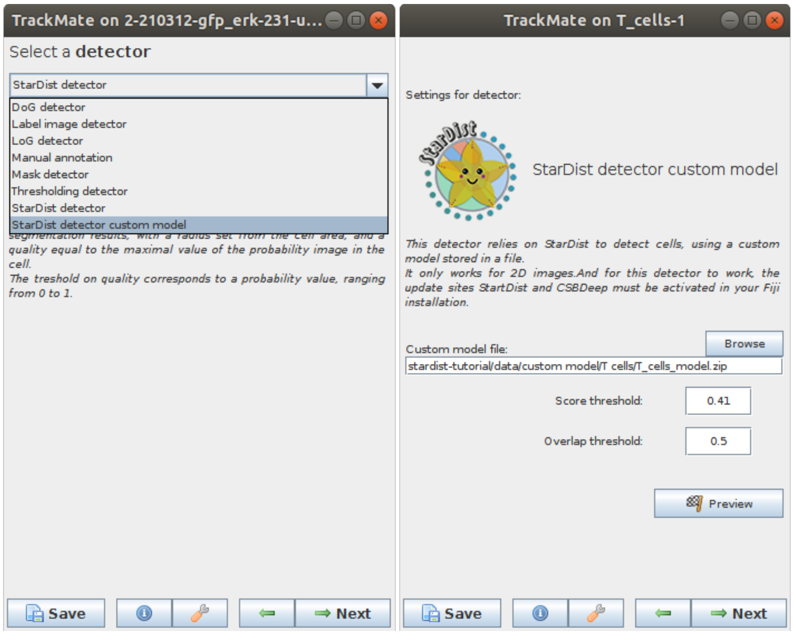

- In the **Custom model file** text field, you need to enter the path to a StarDist model packaged as a zip file (or use the **Browse** button to navigate to the folder). In the tutorial dataset the file is called `StarDistModel-TCellsBF.zip`.
- **Score threshold** correspond to the threshold on the probability map to identify object. It accepts values from 0 to 1.
- **Overlap threshold** correspond to the threshold used in non-maxima suppression step used to separate touching/overlapping objects; it accepts values from 0 to 1.
- Set these parameters and click **Preview** button to test the detector on the current frame. Here is an example of what we get on the first time-point of the tutorial image (using default parameters):

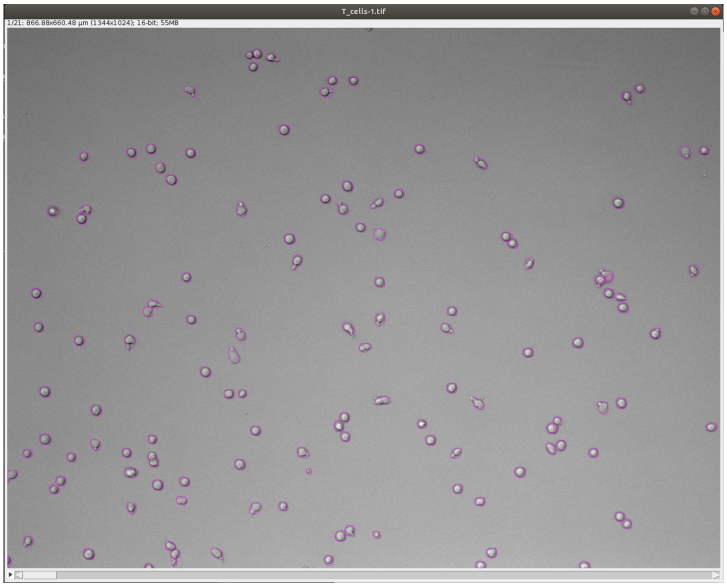

- In case the results are satisfying, click **Next** to perform detection in the full time-series. After the detection is finished, continue with the following steps same as in standard TrackMate workflow. Using the default tracker (Simple LAP tracker) with the default parameters we get this result:

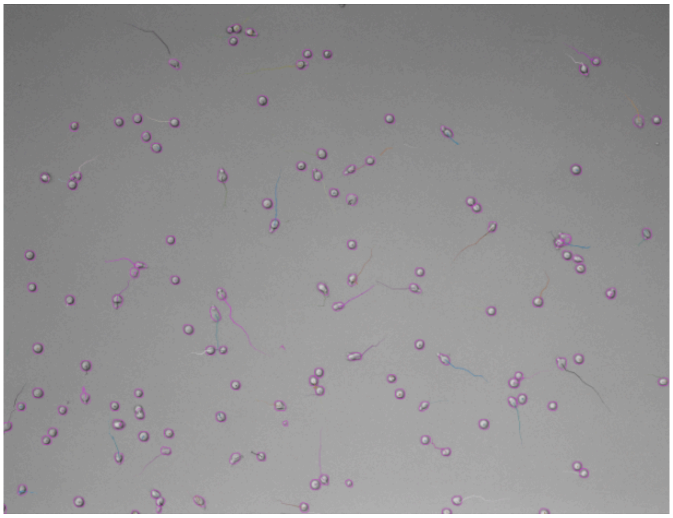

#### Generation of 3D labels by tracking 2D labels using TrackMate.

In this tutorial, you will learn how to generate 3D labels using TrackMate. The segmentation of 3D objects can be very hard. Deep-Learning proves to be very efficient but there are still many models and algorithms that only work for the 2D case.

In this tutorial, we "hack" TrackMate to segment 3D objects using the 2D StarDist segmentation algorithm. We trick TrackMate into thinking a 3D image is a 2D movie over time. We track the fake 2D objects in Z and use the resulting track information to rebuild the 3D segmentation of objects. The following step-by-step tutorial shows how to do this.

First download the dataset from Zenodo:

DOI 10.5281/zenodo.5220610

The dataset contains the raw data but also the intermediate label images in case you want to play with them directly.

- Open Fiji.
- Open the `Spheroid-3D.tif` image in Fiji. This image is a Z stack of MCF10DCIS.com 3D spheroids that have been stained with DAPI to visualise their nuclei.
- Open TrackMate *Plugins > Tracking > TrackMate*. As the image is a 3D Z stack, TrackMate will ask you to swap the Z and T dimensions. This is what we want; click `Yes`.

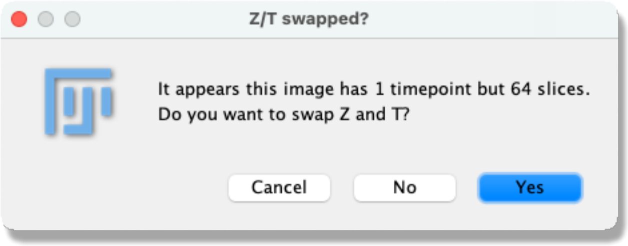

- The start panel will open, showing information about the image dimensions. Click `Next`.
- The `Select a detector` panel opens. From the pull-down menu select the `StarDist detector`. Click `Next`.
- A panel with a description of the StarDist detector opens. By clicking the `Preview` button, you can test the StarDist detector on the current frame. When you are happy with the result, click `Next`.

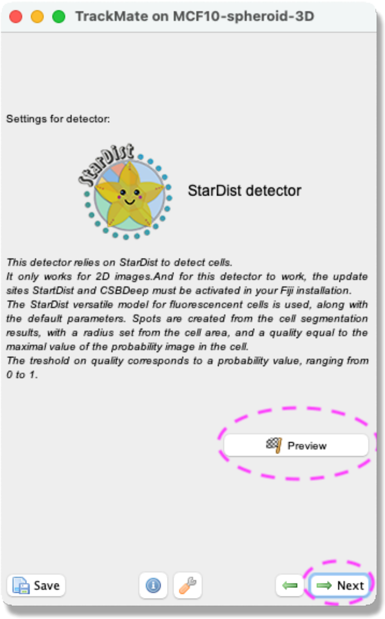

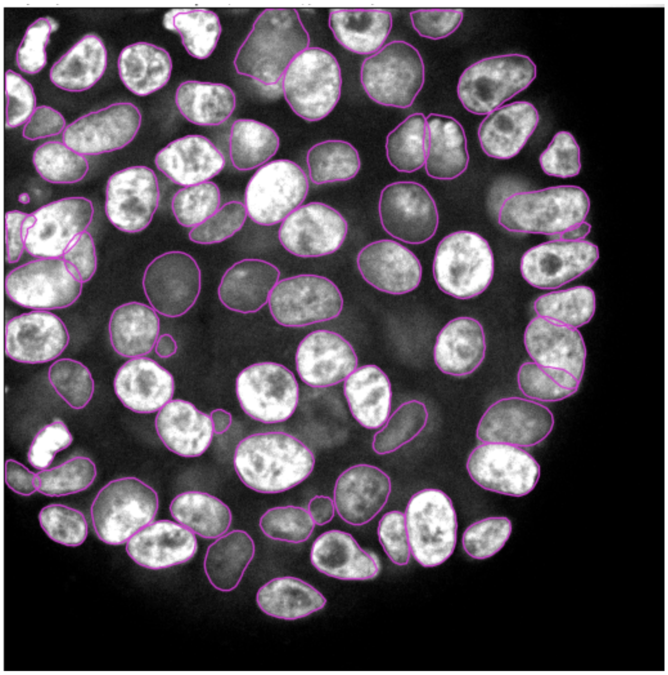

- The detector will detect all nuclei in the selected channel for all time frames. This can take a few minutes.
- When the progress bar has reached the end, click `Next`.
- A panel to filter the detected spots according to their quality opens (more information about this filtering can be found here). In this exercise, this part can be ignored. Click `Next`.
- A panel to filter spots according to their properties (i.e., size, shape, location, or signal intensity) opens. In this exercise, do the following:
  - Click on the green plus sign at the bottom of the panel - a filter appears at the top of the panel.
  - Click on the pull-down menu and select `Area`. Here we will filter out the smallest detected objects. Make sure the `Above` button is selected.
  - Drag the horizontal line (cyan dashed line) to value 31.47.
  - Click on the green plus sign at the bottom of the panel again - a new filter appears.
  - Click on the pull-down menu and select `Max intensity ch1`. Here we will filter out the objects that have low intensity. Make sure the `Above` button is selected. Drag the horizontal line (cyan dashed line) to value 45524.62.
  - Click on `Next`.

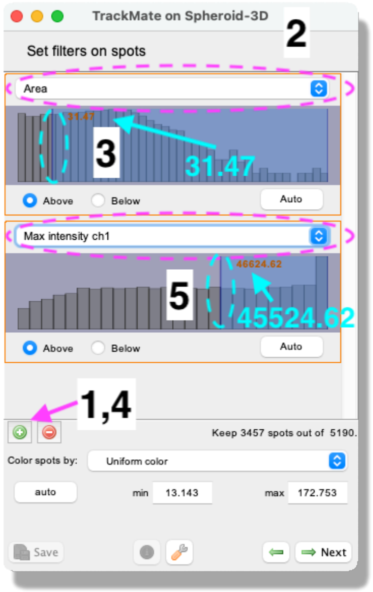

- Next, a tracking panel opens. In this panel, you can select the methods for tracking objects. Here, we use the `LAP tracker`. Please select it from the pull-down menu and click `Next`.

- A panel to choose the LAP tracker settings opens. With this panel, you choose how to track the cells. First, with the `Frame to Frame linking` parameter, you give the maximum distance to link two objects between frames. Here use 4 microns. Next, you tell Trackmate how many spots can be missing and still be the same track. Tick the `Allow gap closing` box and add values: `Max distance`: 4 microns and `Max frame gap`: 3. Click `Next`.

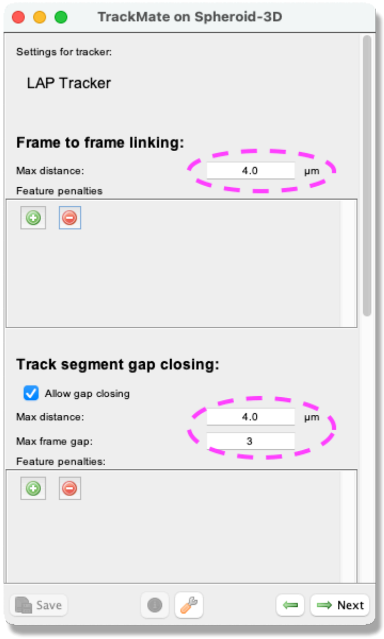

- A Track filter panel opens. In this panel, you can remove tracks according to their properties (i.e., length, speed, or location). Here, we filter out the shortest tracks to remove possible artefacts. Similarly, as in the object filtering above, click the green plus sign to add a filter. Click the pull-down menu and select `Track duration` from the list. Make sure that the `Above` option is ticked and set the slider to 3.45. Click `Next`.

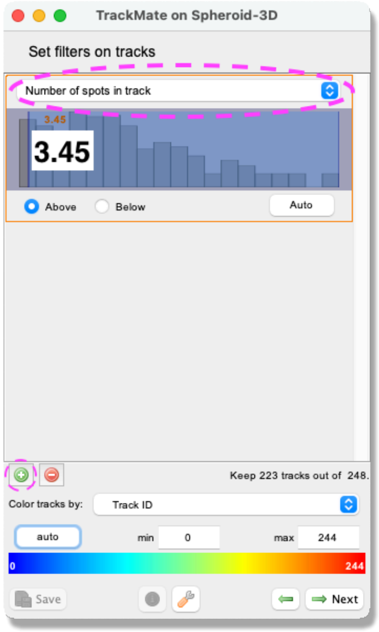

- A window with track visualization options opens. Here it is possible to edit track or object colors according to their properties. We don't need to use it in this exercise. Click on `Next`.

- Click on Next again.

- You should have reached the `Select an action` panel. In this panel, there is a possibility to do different actions. For this exercise, we will export a label image of the tracked objects. From the pull-down menu, select `Export label image` and click on `Execute`.

- From the pop-up window, tick the box `Export only spots in tracks` to eliminate any object not linked to a track and click `OK`.

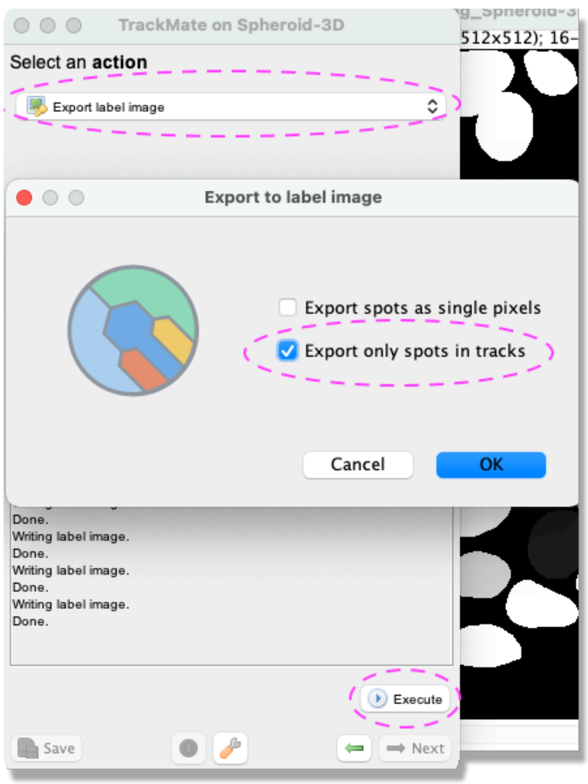

- The label image is now exported. Remember to change the dimensions back from T to Z in Fiji *Image > Properties* and to save your image *File > Save as...*.
- The label image can be viewed in 3D for instance using FPBioimage, and further analyzed using the 3D ImageJ Suite.

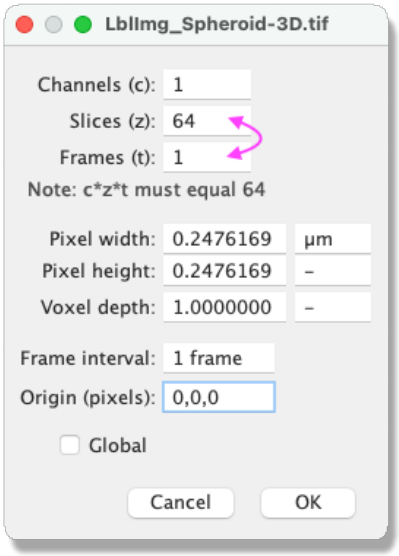

### How to use the new API to create spots with ROIs in TrackMate.

#### Introduction.

Up to the version 7 of TrackMate, detection algorithms were limited to return the position of spots and their radius. All the detection were represented by a tuple in the shape of `frame, x, y, z, radius, quality`. This is well suited to implement *detection algorithms*, that return the position of an object but omit its shape. The previous tutorial series (online) showed how to implement such an algorithm as a detector for TrackMate.

With version 7, we rewrote a large part of TrackMate to remove this limitation, at least in 2D. We changed the data model so that `Spot`s in TrackMate could *possibly* store the shape, while not affecting the existing detectors. We made a new API to facilitate implementing *segmentation algorithms* in TrackMate, that can return the shape of objects. Their shape is later used to compute morphology features or to measure intensity within the object contour. We used this API to implement 7 new segmentation algorithms in TrackMate described above, integrating the some of the best segmentation tools available in Java. This page documents how you can use this API to implement your own segmentation algorithms yourself and make it a first-class-citizen of TrackMate, like any of the other TrackMate modules we document in this series.

We need to first review the new API itself, which basically boils down to one class offering static methods. Then, as for the other detectors, we need to add some flags in the detector factory to tell TrackMate that what we build is a segmentation algorithm that returns the object shape. Finally, we will also see some find tuning of the multithreading for your detectors, made to accommodate the various existing segmentation tools you might want to integrate in TrackMate. We will use the examples of the 7 segmentation algorithms mentioned above to base this tutorial.

#### Creating spots that store object contours.

##### What changed in the `Spot` class.

Starting in version 7, the Spot class in TrackMate has a new field: a SpotRoi object.

It is basically made of the 2D polygon that stores the object contour, relative to the spot center (the `x, y, z` tuple). Its only fields are two `double[]` arrays for the `xp` and `yp` coordinates of the contour, ordered along the contour. The last point connects to the first. The `Spot` objects returned by detection algorithms have the `SpotRoi` set to `null`. The sole difference of a segmentation algorithm in TrackMate is that it returns `Spot` objects with a non-`null` `SpotRoi` object.

#### Limitations.

Let's start with the bad news: We can deal with object contours only for 2D images. The main reason is that we do not have yet a good way to store 3D contours in Fiji. This might change but for now all of what you will see only applies to 2D contours, so polygons.

Also, the object shape must be represented by a simple polygon. That is: the contour must not have holes, and the line of the polygons should not intersect each other. If you create a spot with an intersecting contour, the results are undefined. If you have objects with holes and you try to create spots for them, the holes will be ignored.

#### Creating a `Spot` from a polygon.

The two `double[]` arrays for the `xp` and `yp` must be set relative to the spot X and Y position. You can compute them yourself, but it is easier to use the static method `createSpot()` that returns a spot from `x` and `y` in physical coordinates:

```
double[] xp = ...;
double[] yp = ...;
double quality = ...;
Spot spotWithShape = SpotRoi.createSpot(xp, yp, quality);
```

As for all the coordinates stored in `Spot`, the X and Y must be in physical units (*e.g.* microns if your image is calibrated in microns). This method will take care of computing the spot `x` and `y` center, compute a radius from the polygon, and store the contour points correctly.

#### Example: How spots are created in the StarDist detector.

The Fiji StarDist implementation returns the objects it found as polygons. We therefore just used this method to bridge it to TrackMate. In StarDistDetector.java you will find the following lines:

```
/*
 * We received the 'polygons' object from the StarDist runner.
 * As for all the other detectors, this instance have to return
 * a list of Spot 'spots' containing all the objects segmented in
 * the current time-point.
 */

// Create spots from output.
for ( final Integer polygonID : polygons.getWinner() )
{
    // Collect quality = max of proba.
    final PolygonRoi roi = polygons.getPolygonRoi( polygonID );
    proba.setRoi( roi );
    final double quality = proba.getStatistics( Measurements.MIN_MAX ).max;

    // Create ROI.
    final Polygon polygon = roi.getPolygon();
    final double[] xpoly = new double[ polygon.npoints ];
```

```
    final double[] ypoly = new double[ polygon.npoints ];
    for ( int i = 0; i < polygon.npoints; i++ )
    {
        /*
         * Here we convert the polygon points in pixel coordinates to
         * physical coordinates (multiplication by the calibration).
         * We also need to offset them by the interval top-left corner
         * in case the user ask to perform segmentation in a sub-region
         * of the source image.
         */
        xpoly[ i ] = calibration[ 0 ] * ( interval.min( 0 ) + polygon.xpoints[ i ] );
        ypoly[ i ] = calibration[ 1 ] * ( interval.min( 1 ) + polygon.ypoints[ i ] );
    }
    Spot spot = SpotRoi.createSpot( xpoly, ypoly, quality );
    // 'spots' is the list of Spot this detector will return.
    spots.add( spot );
}
```

As you can see it is fairly simple. It illustrates how you can plug anything that returns a polygon in TrackMate and make a new detector out of it. If you want to integrate a technique that returns instead a mask, a probability map or a label image, we also made utility methods for these cases.

#### Creating a collection of spots from a mask image or a threshold image.

##### The `MaskUtils.fromThresholdWithROI()` method.

The above method is what you could use to implement your own segmentation algorithm. We also provide a simple API to facilitate integrating existing segmentation algorithms in TrackMate. Many of the existing algorithms either return a mask image, a label image or some sort of probability map to threshold to generate objects. This API consists mainly of utility methods in the class MaskUtils that accept such inputs and output a collection of `Spots`, containing the object contour. Let's start with how to input mask images.

For the maximal flexibility, a mask image in TrackMate can be of any type provided it is using a scalar, real pixel type. We simply say that the objects are made by connecting all the pixels that have a value strictly larger than 0. So the method that creates spots from such a mask is the one for importing a threshold image: MaskUtils.fromThresholdWithROI()

```
    /**
     * Creates spots <b>with their ROIs</b> from a <b>2D</b> grayscale image,
     * thresholded to create a mask. A spot is created for each
     * connected-component of the mask, with a size that matches the mask size.
     * The quality of the spots is read from another image, by taking the max
     * pixel value of this image with the ROI.
     *
     * @param <T>
     *            the type of the input image. Must be real, scalar.
     * @param <S>
     *            the type of the quality image. Must be real, scalar.
     * @param input
```

```
     *            the input image. Must be 2D.
     * @param interval
     *            the interval in the input image to analyze.
     * @param calibration
     *            the physical calibration.
     * @param threshold
     *            the threshold to apply to the input image.
     * @param simplify
     *            if <code>true</code> the polygon will be post-processed to be
     *            smoother and contain less points.
     * @param numThreads
     *            how many threads to use for multithreaded computation.
     * @param qualityImage
     *            the image in which to read the quality value.
     * @return a list of spots, with ROI.
     */
    public static final < T extends RealType< T >, S extends NumericType< S > > List<
Spot > fromThresholdWithROI(
                final RandomAccessible< T > input,
                final Interval interval,
                final double[] calibration,
                final double threshold,
                final boolean simplify,
                final int numThreads,
                final RandomAccessibleInterval< S > qualityImage )
```

This method is suitable for 2D images. It will create spots with object contours. Let's review a bit its input:

- `input` is the mask input. It must be a `RandomAccessible` of type `T`, which is the classical frame that TrackMate automatically provides to its detectors.
- `interval` is the interval in the input to analyze. Like for all other detectors, TrackMate returns the pixel infor as an unbounded `RandomAccessible` and we need to specify what part of the image we have to analyze. Again, this is common to all detectors and provided by TrackMate.
- `calibration` is a `double[]` array of 3 elements that contains the pixel size of the image (pixel width, height and depth). It is read from the input calibration that you can set in Fiji. Again, all of this is common to all detectors.
- `threshold` is a double value above above which intensities will be considered par of an object. This is spefic to this detector and is set by the user. For mask images, it is 0.
- `simplify` is a boolean flag that states whether the user requested the contours to be smoothed and simplified. This is important for measuring correctly morphological features and we document it fully elsewhere.
- `numThreads`. The creation of spots in this way is multithreaded and you can set here how many threads to use. Again, if you declare your detector to be `Multithreaded`, the `numThreads` value of your detector will be set automatically by TrackMate and you use it here. If your detector is not multithreaded, use a value of 1.

- The `qualityImage` is an image from which we will read a quality value for the spots created. It must be defined on the same interval that of the `interval` parameter, and the pixels must be of `NumericType` and scalar. If you cannot set the quality from a channel or an image (like for a mask), just pass `null` to this parameter and the quality valueof an object will be set to be its area. Otherwise, it will be the maximal pixel value within the object contour in the quality image.

##### Example: the mask detector.

Let's see how it is used on the mask detector. Since the mask images are treated simply as grayscale images with a threshold of 0, the mask detector is actually implemented in the ThresholdDetector class. (The MaskDetectorFactory returns a `ThresholdDetector` with a threshold value set to 0. See this line.) Here is the content of the `process()` method:

```
@Override
public boolean process()
{
	final long start = System.currentTimeMillis();
	if ( input.numDimensions() == 2 )
	{
		/*
		 * 2D: we compute and store the contour.
		 */
		spots = MaskUtils.fromThresholdWithROI( input, interval, calibration,
							threshold, simplify, numThreads, null );
	}
	else if ( input.numDimensions() == 3 )
	{
		/*
		 * 3D: We create spots of the same volume that of the region.
		 */
		spots = MaskUtils.fromThreshold( input, interval, calibration, threshold,
										numThreads );
	}
	else
	{
		errorMessage = baseErrorMessage + "Required a 2D or 3D input, got "
						+ input.numDimensions() + "D.";
		return false;
	}
	final long end = System.currentTimeMillis();
	this.processingTime = end - start;
	return true;
}
```

Note that the detector treats differently 2D and 3D images. As stated above, the new API supports object contour only for 2D images. The method `MaskUtils.fromThreshold()` is the complement of the `MaskUtils.fromThresholdWithROI()` for 3D images, but that returns `Spot` objects that do not have a contour. Here the spots created have a radius that is such that the sphere with this radius have the same volume that of the segmented object.

The use of this API makes the detector code very short. You can imagine adapting the same approach to integrate a segmentation algorithm to would output a mask image or a threshold image. For instance, this is what we did to integrate the *Traininable Weka segmentation* plugin and the *ilastik* pixel classifier.

#### Example: the Weka detector.

The Weka detector is not very complicated. The bulk of calling Weka is done in the WekaRunner class. Running Weka is done in the WekaRunner.computeProbabilities() method. It returns the probability of the classification for the specified input and the specified class. We won't detail it. But creating spots from this probability is simple. The method WekaRunner.getSpots() method, which resembles the method described in the previous paragraph:

```
public List< Spot > getSpots( final RandomAccessibleInterval< T > proba,
     final double[] calibration, final double threshold, final boolean simplify )
{
       final List< Spot > spots;
       if ( isProcessing3D )
       {
              spots = MaskUtils.fromThreshold(
                            proba,
                            proba,
                            calibration,
                            threshold,
                            numThreads,
                            proba );
       }
       else
       {
              spots = MaskUtils.fromThresholdWithROI(
                            proba,
                            proba,
                            calibration,
                            threshold,
                            simplify,
                            numThreads,
                            proba );
       }
       return spots;
}
```

Here the threshold value to segment the probability map is set by the user. Since we have a probability map, we can use it to compute a quality value derived from this probability.

#### Example: the ilastik detector.

The ilastik detector works exactly in the same way. It has a IlastikRunner class that is in charge of calling ilastik and convert the results to a spot collection. The ilastik detector just makes a simple call to it.

However, we use a special slicing of time-points for this algorithm. Indeed, the ilastik runner expects to receive *all* the time-points to process at once, runs ilastik on them, and then return. We will discuss this in the next section.

Here is some explanation on the runner code:

```
/*
 * This corresponds roughly to the lines 94-110 of the IlastikRunner class.
 */

// Path to the ilastik project, provided by the users.
final File projectFile = new File( projectFilePath );

// Create an ilastik pixel classifier, configured with the classifier in the specified project.
final PixelClassification classifier = new PixelClassification(
  executableFilePath,
  projectFile,
  logService,
  statusService,
  numThreads,
  maxRamMb );
final PixelPredictionType predictionType = PixelPredictionType.Probabilities;

// Run the classifier on the 'cropped' source image. This will result in the PixelClassification
// Actually RUNNING ilastik in the background, passing input and output images as files saved
// in a temp folder. But this is transparent to us.
final ImgPlus< T > output = classifier.classifyPixels( cropped, predictionType );

// The output has one channel per class in the classifier, so we need to get the channel that
// contains the probability for our object of interest only (specified by the user via the classID
// parameter).
final ImgPlus< T > proba = ImgPlusViews.hyperSlice( output, output.dimensionIndex( Axes.CHANNEL ),
classId );

// Etc.
...

// Not we just have to import this probability map as TrackMate ROIs. Since we received the proba
// for ALL time-points at once, we need to process it time-point by time-point:

for ( int t = 0; t < proba.dimension( timeIndex ); t++ )
                {
                        final List< Spot > spotsThisFrame;
                        final ImgPlus< T > probaThisFrame = TMUtils.hyperSlice( proba, 0, t );

                        if ( DetectionUtils.is2D( probaThisFrame ) )
                        {
                                /*
                                 * 2D: we compute and store the contour.
                                 */
                                final boolean simplify = true;

        // In 2D we again use the MaskUtils utilities. Note that we use the '...WithROI()'
        // method version. In 2D we can import objects with their contours.
                                spotsThisFrame = MaskUtils.fromThresholdWithROI(
                                                probaThisFrame,
                                                probaThisFrame,
                                                calibration,
                                                probaThreshold,
```

```
                                        simplify,
                                        numThreads,
                                        probaThisFrame );
                    }
                    else
                    {
                        /*
                         * 3D: We create spots of the same volume that of the region.
                         * So we use the methods without the '...WithROI()'.
                         */
                        spotsThisFrame = MaskUtils.fromThreshold(
                                        probaThisFrame,
                                        probaThisFrame,
                                        calibration,
                                        probaThreshold,
                                        numThreads,
                                        probaThisFrame );
                    }
// etc...
```

#### Controlling the slicing of time-points.

Normally TrackMate automates the multi-threading processing of several time-points. When you develop a detector, a `SpotDetector` instance is supposed to operate only on one time-point. The processing logic in TrackMate will take care of providing a single time-point image to this detector and incorporate the detection results into the correct place in the `SpotCollection`.

But because we wanted to integrate with algorithms and tools such as ilastik, we needed to provide another way of handling time-points. For instance, ilastik expects to receive all the time-points at once, process them, and return the probabilities for all time-points as well. This is considerably faster than calling ilastik several times, once per time-point.

So starting with v7, there is a new hierarchy for `SpotDetectorFactory` in TrackMate:

##### `SpotDetectorFactory`

SpotDetectorFactory is the initial interface for spot detector factories that generate one detector per time-point. It is well suited to detection and segmentation algorithms that can run concurrently on several time-points at once without a penalty too large. All detectors I know of, except the ilastik detector, derive from this interface.

Note that its only specific method is the following:

```
/**
 * Returns a new {@link SpotDetector} configured to operate on the given
 * target frame. This factory must be first given the <code>ImgPlus</code>
 * and the settings map, through the <code>#setTarget(ImgPlus, Map)</code>
 * method.
 *
 * @param interval
 *            the interval that determines the region in the source image to
 *            operate on. This must <b>not</b> have a dimension for time
 *            (<i>e.g.</i> if the source image is 2D+T (3D), then the
 *            interval must be 2D; if the source image is 3D without time,
```

```
 *            then the interval must be 3D).
 * @param frame
 *            the frame index in the source image to operate on
 */
public SpotDetector< T > getDetector( final Interval interval, int frame );
```

So it will generate a `SpotDetector` instance per time-point. Such a detector has for output a `List< Spot >`, which is expected to contain only the spots it found in the frame it is configured to run on. TrackMate will take care of assembling all the `List< Spot >` from different time-point in a `SpotCollection`.

##### `SpotGlobalDetectorFactory`

SpotGlobalDetectorFactory is a new interface that does not slice time-points. Its unique specific method returns a `SpotGlobalDetector` that is expected to process *all* time-points at once. Such a detector is instantiated only once per detector run.

```
/**
 * Returns a new {@link SpotDetector} configured to operate on all the
 * time-points. This factory must be first given the <code>ImgPlus</code>
 * and the settings map, through the <code>#setTarget(ImgPlus, Map)</code>
 * method.
 *
 * @param interval
 *            the interval that determines the region in the source image to
 *            operate on. This must <b>not</b> have a dimension for time
 *            (<i>e.g.</i> if the source image is 2D+T (3D), then the
 *            interval must be 2D; if the source image is 3D without time,
 *            then the interval must be 3D).
 */
public SpotGlobalDetector< T > getDetector( final Interval interval );
```

The `SpotGlobalDetector` outputs a `SpotCollection`, that is expected to contain all the spots in all time-point of the movie.

#### The base `SpotDetectorFactoryBase` interface and the `has2Dsegmentation` flag.

The two interfaces above derive from a mother interface that contains common methods: SpotDetectorFactoryBase. It contains a very important method if are building a segmentation algorithm for TrackMate:

```
/**
 * Return <code>true</code> for the detectors that can provide a spot with a
 * 2D <code>SpotRoi</code> when they operate on 2D images.
 * <p>
 * This flag may be used by clients to exploit the fact that the spots
 * created with this detector will have a contour that can be used
 * <i>e.g.</i> to compute morphological features. The default is
 * <code>false</code>, indicating that this detector provides spots as a X,
 * Y, Z, radius tuple.
 *
```

```
 * @return <code>true</code> if the spots created by this detector have a 2D
 *         contour.
 */
public default boolean has2Dsegmentation()
{
      return false;
}
```

If your detector can return the spot shape for 2D images, override this method so that it returns true. This is important to let TrackMate know it should compute morphological features on the spot this detector created. If you don't see the morphological features in the GUI (Area, ...) this is most likely because of this method.

As you can see in the examples above in this page, the new detectors return true for this method.
